# Human-mouse cross-species comparison identifies common and unique aspects of intestinal mesenchyme development

**DOI:** 10.1101/2025.09.03.674033

**Authors:** Kelli Johnson, Xiangning Dong, Zhiwei Xiao, Hamza Islam, Meghan Anderman, Ian Glass, Jason R. Spence, Katherine D. Walton

## Abstract

Using single-cell RNA sequencing (scRNA-seq) and histological approaches, we examine cross-species cellular identity, diversity and organization of mesenchymal (fibroblast) populations in the developing mouse and human intestines. In both species, we defined 7 fibroblast populations. Using cross-species integration and label transfer approaches we find that each mesenchymal cell subtype in the murine intestine is highly concordant to a cell type/state in the human intestine, and vice-versa, suggesting a strong conservation of mesenchymal cell types/states across species. Despite this conservation, we also observe that individual lineage-defining genes are not always shared and can be found in different mesenchymal populations. High resolution spatial analysis via fluorescent in situ hybridization (FISH) and immunofluorescence (IF) confirmed these findings and revealed that transcriptionally-defined sub-types of intestinal mesenchymal cells in mice and humans are organized within similar spatial domains.

## Introduction

Mouse models are frequently used in biomedical research due to their genetic tractability, and physiological similarity to humans. Studying development in the mouse and extrapolating the results to human physiology can bridge gaps in our knowledge that cannot be addressed directly in humans; however, there are also unique differences across species^1^. By understanding the similarities and differences between developing gastrointestinal systems of human and mouse, we gain insight into the common and divergent mechanisms of intestinal development, homeostasis and disease, which can be leveraged to improve disease diagnosis and the treatment of gastrointestinal disorders.

During early development (embryonic day (E)7.5 in the mouse and 16-17dpc (days post conception in the human), the small intestine begins as a flat sheet of endoderm and associated mesoderm^2,3^. By mouse E8 and human 18dpc, the flat sheet begins fold and close into a tube of endoderm surrounded by a tube of mesoderm by E9.5/22-30dpc^3,4^. The endoderm will give rise to the epithelial lining of the intestine, while the mesoderm eventually becomes the various mesenchymal components (fibroblasts, smooth muscle, vasculature, pericytes and resident immune cells). Regionalization of this tube into the stomach, small intestine and large intestine is noticeable by E8.5/22dpc; with distinct domains of region-specific epithelial markers^3,5,6^. Finally, the third embryonic germ layer, the ectoderm, gives rise to the enteric nervous system (ENS). Neural crest cells migrate from the neural tube to populate the intestine between E8.5-13.5/22-49dpc^7,8^.

Reciprocal signaling between the epithelium and mesenchyme is essential for coordinated patterning of the absorptive unit into crypts and villi ^9,10^. Emergence of the villi from a flat epithelium is coordinated through reciprocal epithelial-mesenchymal signaling that leads to the aggregation of mesoderm-derived fibroblasts directly adjacent to the epithelium, marked by high expression of PDGFRA (also called subepithelial cells (SECs) or telocytes^11^) into mesenchymal clusters^12,13^. Mesenchymal clusters are post-mitotic and act as signaling centers, secreting a variety of ligands to the epithelium and neighboring mesenchyme^14-17^. When cluster formation is inhibited, villi fail to emerge and when the pattern of cluster formation is altered, emerging villi are improperly patterned^12,13^. This underscores the importance of the mesenchyme during intestinal development and villus morphogenesis.

Recent single-cell approaches have attempted to define the heterogeneity and spatial organization of the developing and adult murine^18-24^ and human^25-34^ intestines. A particular strength of these studies has been resolving the heterogeneity within the mesenchymal cell populations (also referred to in the literature as ‘fibroblasts’ or ‘stromal cells’), which generally encompasses the cells that help to establish and maintain intestinal structure. Despite these recent advances, some ambiguity between the species still exists based on cell nomenclature and the associated molecular identities of individual cell populations. For example, cells lining the murine epithelium are referred to as ‘telocytes’ while they are called ‘subepithelial cells’ (SECs) in the human intestine; nonetheless, both populations are identified by high levels of PDGFRA and FOXL1 expression. While the molecular identities of heterogeneous cell types have emerged (e.g. murine ‘trophocytes’ defined by expression of R-SPONDIN ligands) in both species, there are still some gaps in our knowledge about the heterogeneity of mesenchyme, molecular identities and species specificity. Here, we aimed to address two major gaps by: 1) defining mesenchymal cell heterogeneity during mouse intestinal development (E13.5-E17.5); 2) directly comparing mesenchymal/fibroblast cell populations present in the mouse to the human fetal intestine using multiple integrated published datasets^28,29,35,36^.

In both species, we identified all major cell classes (epithelium, mesenchyme/fibroblasts, smooth muscle, nerves, endothelium, immune cells, and mesothelium). Mesenchyme and smooth muscle populations were further interrogated to define sub-populations of cells based on their molecular identity, and follow-up imaging analysis was used to define the position of each sub-population within the intestine. Leveraging the single-cell data, we used label transfer approaches to determine similarities and differences between the murine and human cell population. Through our analysis, we identified new/novel cell populations within the mesenchyme/fibroblast compartment in both species, which will be the subject of future investigation, and we found that all the major mesenchyme/fibroblast sub-populations are conserved between species. However, even though we observed conservation at the population level, we found divergence at the individual gene level, where important cell-defining marker genes in one species may be expressed in the other species in a different population and vice versa. Taken together, we define mesenchyme/fibroblast heterogeneity in the developing mouse and human intestine, and find that while populations are conserved across species, individual genes that act as important markers for a population in one species may not be expressed in the corresponding population in the other species.

## Results

### Interrogating cellular heterogeneity in the developing mouse and human intestine

We generated single-cell RNA sequencing (scRNA-seq) data from dissociated whole mouse small intestine (proximal duodenum through distal ileum) each day from E13.5-E17.5 (**Supplemental Figure 1A**). This encompasses critical stages before, during and after villus morphogenesis and lead to the identification of epithelial, mesenchymal (also known as fibroblast/stroma), ENS, mesothelium, immune and endothelial cells (**Supplemental Figure 1B**). Cells were identified based on expression of previously defined and well accepted markers (**Supplemental Figure 1C**)^25,37,38,39,20,22,28,29,19,27,32,40^. For each timepoint analyzed, we plotted the proportion of cells contributing to each cluster as a percent of the total sample at each stage (**Supplemental Figure 1D**) or as a percent of each sample that contributed to the total cluster (**Supplemental Figure 1E**,**J**).

**Figure 1.**
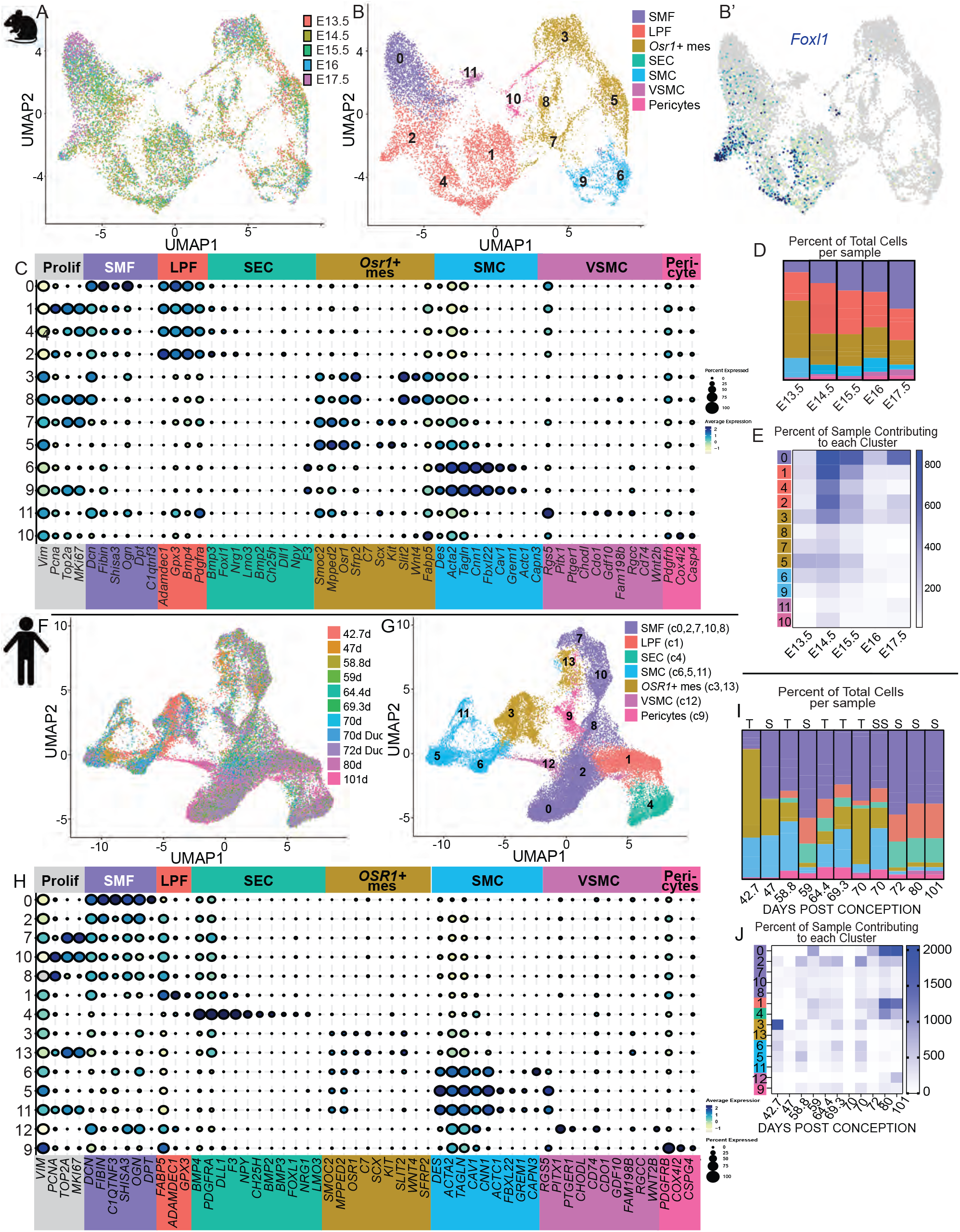
Comparison of single-cell sequencing of all mouse mesenchymal cells from E13.5 to E17.5 and all human cells from 42.7-101dpc. A,F) UMAP rendering of mesenchymal samples. B,G) UMAP rendering of cells clustered by transcriptional similarity. B’) Featureplot rendering of Foxl1+ expressing cells on the UMAP from B. C,H) Dotplot of average expression of transcripts in each cluster. D,I) Parts of the whole distribution plot of the contribution each cluster per stage. E,J) Heat map of the percent of the sample contributing to a cluster at each stage. Color coding of clusters is the same in B, C, D, E, G, H, I and J.

To interrogate human data that corresponds to the key morphologic stages before, during and after villus morphogenesis, we integrated two different published human fetal datasets^28,29^ from 42.7-101dpc (**Supplemental Figure 1F-J**). After quality filtering, 11 different samples combined for a total of 62,545 cells. Previously described cell types were identified based on highly expressed canonical genes (**Supplemental Figure 1H**), demonstrating epithelial, mesothelial, mesenchyme/fibroblast, smooth muscle, endothelial, immune and neural crest populations in the data (**Supplemental Figure 1G**,**H)**. For each sample analyzed, we plotted the proportion of cells contributing to each cluster as a percent of the total sample at each stage (**Supplemental Figure 1I**) or as a percent of each sample that contributed to the total cluster (**Supplemental Figure 1J**).

### Interrogating epithelial cell types in the developing mouse and human intestine

To ensure the quality of our datasets, we interrogated the epithelial cells from both species since they have been well characterized in the past using lineage tracing, classic histological^41-46^ and single-cell sequencing approaches^20,26,28,29,31,32,34^ (**Supplemental Figure 2**). Epithelial clusters from both mouse and human were extracted, re-clustered and re-annotated them to confirm the presence of cell types in our dataset (**Supplemental Figure 2A-B, F-G**). In both the mouse and human, we were able to identify distinct epithelial sub-populations using well-defined marker genes (**Supplemental Figure C, H**) including progenitor populations, absorptive cells (enterocytes) and secretory cells (goblet, enteroendocrine). For each sample analyzed, we plotted the proportion of cells contributing to each cluster as a percent of the total sample at each stage (**Supplemental Figure 2D**,**I**) or as a percent of each sample that contributed to the total cluster (**Supplemental Figure 2E, J**).

### Interrogating mesenchymal cell heterogeneity in the developing mouse and human intestine

The understanding of mesenchymal heterogeneity is continually evolving and a wide array of nomenclature has been used to describe various cell types identified (^27,37,47,19,48^). Here, we define mesenchymal cell populations based on transcriptional markers that distinguish them. Mouse mesenchyme/fibroblasts and smooth muscle populations were extracted (**Supplemental Figure 1B**, clusters (c)2, 3, 5, 8 and 9) and re-clustered into 12 clusters that we could categorize into 7 cell types (**Figure 1A-B**).

Clusters could be partitioned into two broad groups based on their function: mesenchyme/fibroblasts (c0,1, 2, 4 and 10) and smooth muscle (c3,5, 6, 7, 8, 9 and 11). Fibroblasts in the murine intestine were annotated in part based on recent work from our group that defined human fibroblast populations in the human intestine using spatial transcriptomics^49^. Fibroblasts (c0) that highly express *Fibin, Shisa3* and *Ogn*, are named submucosal fibroblasts (SMFs; **Figure 1C**, c0). Lamina propria fibroblasts (LPFs; **Figure 1C**, c1, 4, 2) are identified by *Adamdec1, Gpx3, Bmp4* and *Pdgfra*. A previously described population of cells lining the intestinal epithelium have been identified, well characterized and called ‘telocytes’^11,23,48,50^. We did not identify a distinct telocyte population within these clusters; however, within the LPF population (c2), we identified a small population of cells that expressed the telocyte-specific marker *Foxl1* (**Figure 1B’**). We therefore further subclustered Figure 1B, c2 and observed a small population defined by the telocyte marker *Foxl1* as well as *Pdgfra*^*HI*^, *Nrg1, Dll1* and *F3* (**Supplemental Figure 3**). We also identified four clusters that are defined by expression of *Osr1* and *Smoc2*, referred to as *Osr1+* mesenchyme (**Figure 1B**,**C**, c3,8,7,5). *Osr1*+ mesenchyme can be identified by expression of either *Scx* and *Kit* (c7, c5) or *Slit2* and *Wnt4* (c3, c8) and each of the *Osr1*+ sub-types possess one cluster that is proliferating (c7 and c8 - *Pcna, Top2a*, and *MKi67*). Mature smooth muscle cells (SMCs – **Figure 1C**, c6 and c9) express high levels of the smooth muscle markers *Acta2, Tagln* and *Cnn1*, and one cluster (c9) expresses proliferation markers. The vascular smooth muscle cell (VSMC) markers *Rgs5* and *Rgcc* are highly expressed by c11, while the pericyte markers *Pdgfrb, Cox4i2*, and *Cspg4* are expressed by c10.

Human mesenchyme/fibroblast and smooth muscle cells (**Supplemental Figure 1G**, c0, 2, 3, 6, 8 and 11) were extracted and re-clustered into 14 clusters representing 7 cell types (**Figure 1F-H**). As in the mouse, the intestinal mesenchyme was divided into two main categories: fibroblasts and smooth muscle (**Figure 1G**). The fibroblasts, some of which have recently been defined in the developing human small intestine using spatial transcriptomics^49^, were divided into an SMF population (**Figure 1G-H**, c0,2,7,10 and 8) expressing *SHISA3, FIBIN, C1TNF3, OGN* and include 3 clusters of proliferating cells (*PCNA, TOP2A* and *MKI67* in c7, 8 and 10). There is also an LPF population expressing *ADAMDEC1, GPX3* and *FABP5* (**Figure 1G-H**, c1). Unlike in the mouse, the human telocyte population, previously called subepithelial cells (SECs), clustered separately from the LPF population (likely due the abundance of these cells in the dataset) and expressed *F3, DLL1, NPY, FOXL1 and NRG1* (**Figure 1G-H**, c3).

Similar to the mouse, we also identified *OSR1*+ mesenchyme in the human dataset, defined by expression of *OSR1, SMOC2, MPPED2, OSR1, KIT*^51^, *SLIT2*^52^ *and C7* (**Figure 1G-H**; c3, 13). The *OSR1*+ mesenchyme, could be further divided into proliferative (c13) and non-proliferative (c3). Finally, we identified SMCs and vascular SMCs (VSMCs) (**Figure 1G-H**; c5, 6, 11, 12). The mature smooth muscle populations express high levels of smooth muscle markers *ACTA2, TAGLN*, and *CNN1* (c5, 6 and 11), and can be further divided by expression of *ACTC1*^*HI*^ (c5-non-proliferative; c11-proliferative) and *CAPN3*^*HI*^ (c6). VSMCs (c12) are identified by *PITX1*^*53*^, *PTGER1*^*54*^, and *CD74*^*55*^ are expressed by cells in cluster 12. Finally, pericytes expressed *PDGFRB, COX4I2*, and *CSPG4* (c9).

### Cross-species comparison of mesenchymal/fibroblast populations in the developing intestine

Annotation of cell populations within the mouse and human datasets based on enriched and highly expressed genes suggested that many cell populations within the murine intestine had an analogous population within the human intestine. To directly compare similarities and differences between the developing human and mouse intestine, we used a label prediction approach^56^. To do this, we first converted mouse gene names to their human homologs, then integrated the complete dataset to ensure the homology mapping was successful. After brief validation of the homology-translated data to verify that populations and markers remained largely unaffected, the complete dataset was re-separated by species-of-origin (**Figure 2A-B**) before being re-annotated using expression markers identified pre-integration (**Supplemental Figure 1**) and the similarity of each cell class or mesenchymal population between the two species was calculated.

**Figure 2.**
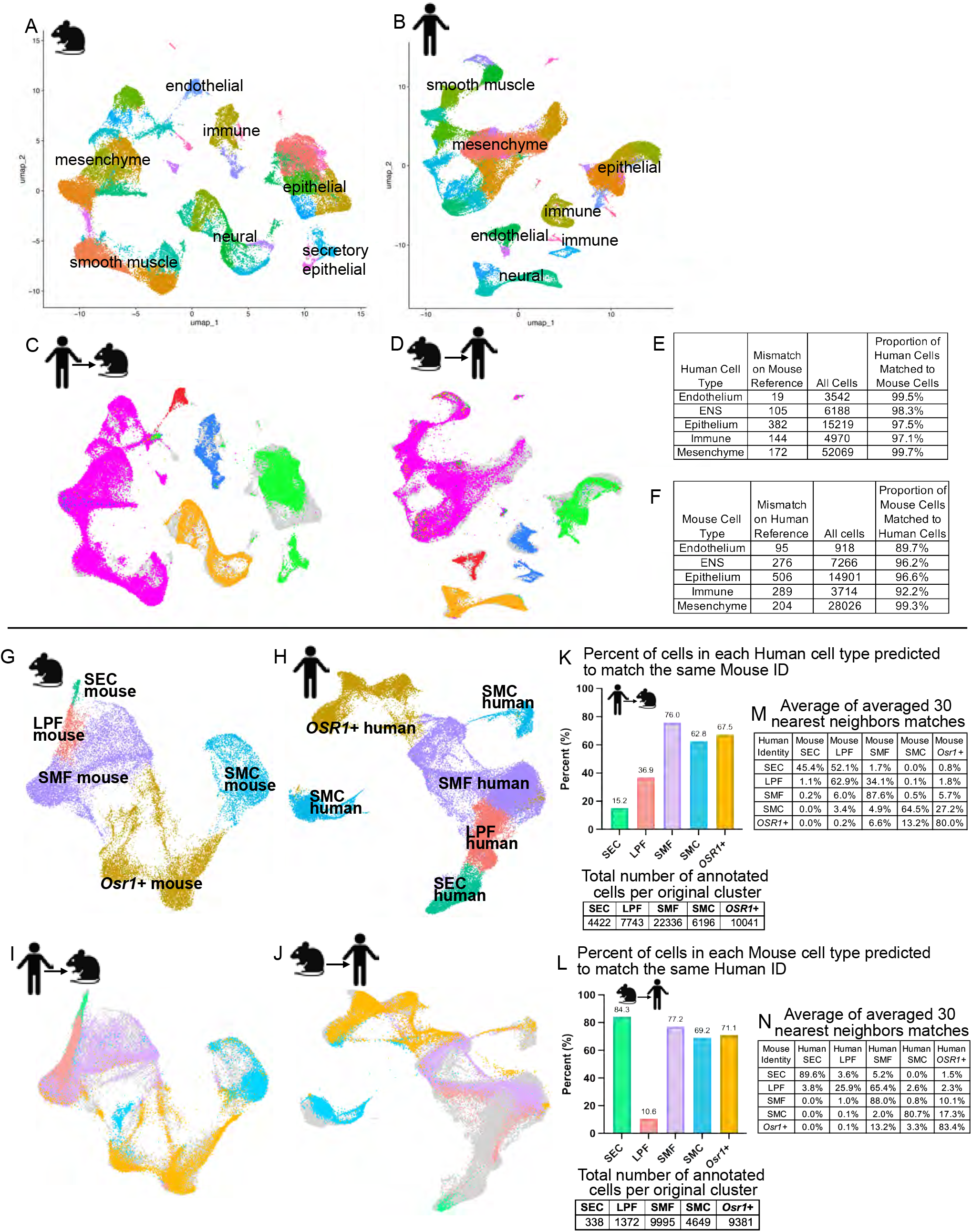
Comparison of the full mouse and human homologous datasets. A,B) UMAP rendering of mouse or human homologs clustered by similarity with cell identities deciphered from the highest expressed transcripts in each cluster. C) Pseudo-colored rendering of human samples projected onto the mouse reference UMAP (gray). D) Pseudo-colored mouse samples projected onto the human reference UMAP (gray). E,F) Tables of human or mouse percent match to their true identity when projected onto the other species. G,H) UMAP rendering of extracted mouse or human mesenchymal homologs clustered by similarity with cell identities deciphered from the highest expressed transcripts in each cluster. I) Pseudo-colored Human mesenchymal homologs projected onto the mouse reference. J) Pseudo-colored mouse homologs projected onto the human reference. K,L) Percentage of human or mouse homologs that matched their true identity when projected onto the other species as determined by nearest neighbor comparison of their transcripts. M,N) The average the averaged percentage match to the 30 nearest neighbors for each mouse or human sample.

Human cell identities were queried against the reference mouse dataset (**Figure 2C**) and mouse cell identities were queried against the human reference dataset (**Figure 2D**) to determine the similarity of each class across species. Similarity was determined using a strict nearest neighbor (NN) calculation that required 93.3% of the closest NNs to match identities. Cells were considered a ‘match’ if the identity of the query cell was the same as >27 of the 30 NNs of the reference dataset. If 3 or more of the 30 NNs were annotated as any other cell type than the query cell, then the prediction was considered a ‘mismatch’.

Based on label transfer of major cell classes (epithelial, mesenchymal, immune, neuronal, endothelial), most cells mapped to their equivalent counterparts (**Figure 2E-F**). Interestingly, the mesenchymal population showed the highest fidelity, with 99.27% of mouse mesenchymal cells mapping to the human mesenchyme and 99.76% of human mesenchymal cells mapping to the mouse mesenchyme. The other classes also mapped mostly to themselves between species: ENS (Mouse-to-Human (*Mm*-to-*Hs*): 96.2% and *Hs*-to-*Mm*: 98.3%), epithelium (*Mm*-to-*Hs*: 96.6% and *Hs*-to-*Mm*: 97.5%), immune (*Mm*-to-*Hs*: 92.2% and *Hs*-to-*Mm*: 97.1%) and finally the endothelium (*Mm*-to-*Hs*: 89.7% and *Hs*-to-*Mm*: 99.5%). Generally, this finding was expected, since the major cell classes are very transcriptionally distinct from one another (i.e. epithelium is highly distinct from immune cells transcriptionally). Nonetheless, this data shows that the label transfer approach is appropriate for comparing cell populations across species.

To test similarity between mesenchymal/fibroblast populations, the mesenchymal populations from both species were extracted (excluding VSMC and pericyte populations discussed in **Figure 1**) and re-clustered (**Figure 2G-H**). The extracted human mesenchymal cells were projected onto the mouse reference dataset (**Figure 2I**) and mouse cells onto the human reference dataset (**Figure 2J**). The percent of each cell type mapping with >93.3% confidence to the 30 NN’s (28 out of 30 cells) from each species comparison was calculated (**Figure 2K-L**). The average composition of the 30 nearest neighbors for each query cell type was also calculated as a way of determining to which reference populations the cell types with low match percentages were most similar (**Figure 2M,N**).

When human mesenchymal cells were queried against the mouse reference dataset (**Figure 2I, K**), the human SMFs had the highest fidelity between species, mapping to mouse SMFs 76.0% of the time. 67.5% of human *OSR1*+ mesenchyme mapped to mouse *OSR1*+ mesenchyme and 62.8% of human SMCs mapped to mouse SMCs. On the other hand, human LPFs only mapped 36.9% of the time to mouse LPFs, while human SECs mapped to mouse telocytes/SECs 15.2% of the time with the most common nearest neighbor defined match being mouse LPFs (52.1%, **Figure 2M**).

Human SECs mismatching to mouse LPFs aligns with our finding that the mouse SECs clustered with mouse LPFs (**Figure 1B**). When human LPF cells were mismatched, they were more likely to match mouse SMFs (**Figure 2M)**.

When mouse cells were queried against the human reference dataset (**Figure 3J, L**), the three most similar populations from the human-to-mouse comparison retained a high level of fidelity. That is, mouse SMFs matched human 77.2% of the time, mouse *Osr1*+ mesenchyme matched human 71.1% of the time, and mouse SMC matched human 69.2% of the time. Interestingly, mapping the mouse SEC data onto human showed a high degree of similarity (84.3%)(something that was not true for human-to-mouse), possibly due to the significantly lower number of telocytes/SECs identified in the mouse data (n=338 mouse, n=4422 human). Mouse LPFs only mapped onto human

**Figure 3.**
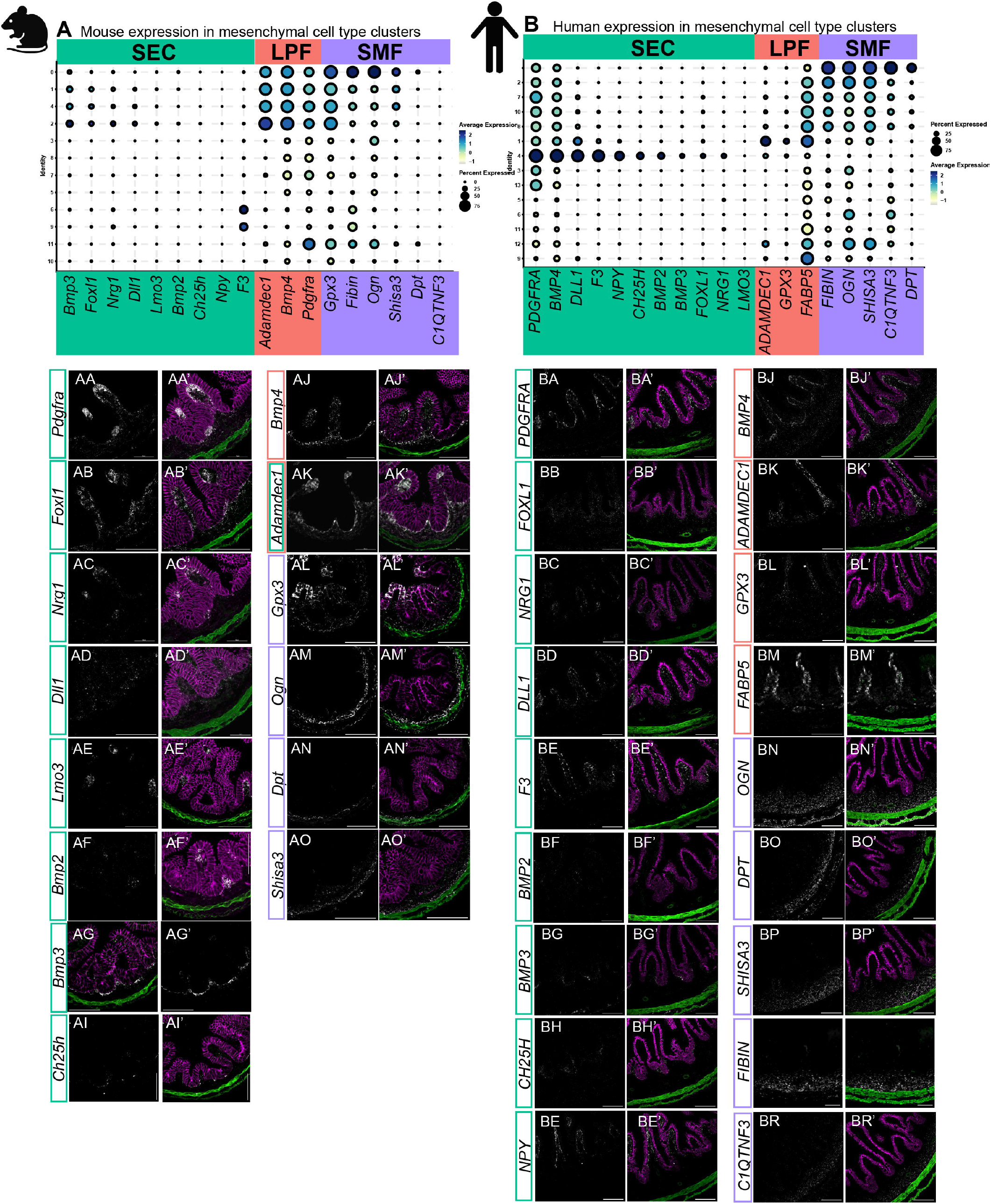
Mesenchymal fibroblast cell type markers and their RNA transcript localization. A,B) Dotplot of the average transcript level in percent of cells per cluster in mouse (A) or human (B). AA-AO, BA-BR) RNA localization of indicated transcripts at E16.5 in mouse (AA-AO) or human at 80d (BA-BR). AA’-AO’, BA’-BR’) same tissue sections as in AA-AO or BA-BR, respectively but overlaid with ECADHERIN (magenta) and SM22/Tagln (green). Background boxes highlight groups of fibroblast cell type markers: green are subepithelial cells, orange are lamina propria fibroblasts and purple are submucosal fibroblasts. Scalebars are 50um.

LPFs 10.6% percent of the time; in fact, based on the average of the nearest neighbors, mouse LPFs are more likely to match to human SMFs (65.4%, **Figure 2N**)

### Spatial localization of mesenchymal and smooth muscle populations within the developing mouse and human small intestine

Using fluorescent *in situ* hybridization (FISH), we visualized the localization of both known and newly identified markers for mesenchymal sub-populations identified in both species (**Figures 3 and 4**). Integrating this localization data, we provide a species-specific map of the cell localizations and their molecular identifiers (**Figure 5**) to characterize the mesenchyme/fibroblast and smooth muscle composition of the human and mouse small intestine. In both species, immediately adjacent to the epithelium are the telocytes/SECs. These cells express high levels of *Pdgfra/PDGFRA*, as previously reported^16,28^ (**Figure 3A, AA, AA’; B, BA, BA’**. The well-defined mouse telocyte/SEC marker, *Foxl1*^50^, shows the expected localization (**Figure 3AB, AB’**); and while it is highly expressed in the human sequencing data, it is observed in only a small subset of cells (**Figure 3B, BB, BB’**). *Nrg1/NRG1* expression matches the previously described localization in telocyte/SECs in both mouse and the human tissue (**Figure 3AC, AC’ and BC, BC’**) but it is also expressed in fewer cells in the human (**Figure 3B**). *Dll1/DLL1* is largely localized to telocytes/SECs but is also expressed in some LPFs in both mouse and human (**Figure 3A, AD, AD’ and B, BD, BD’**).

**Figure 4.**
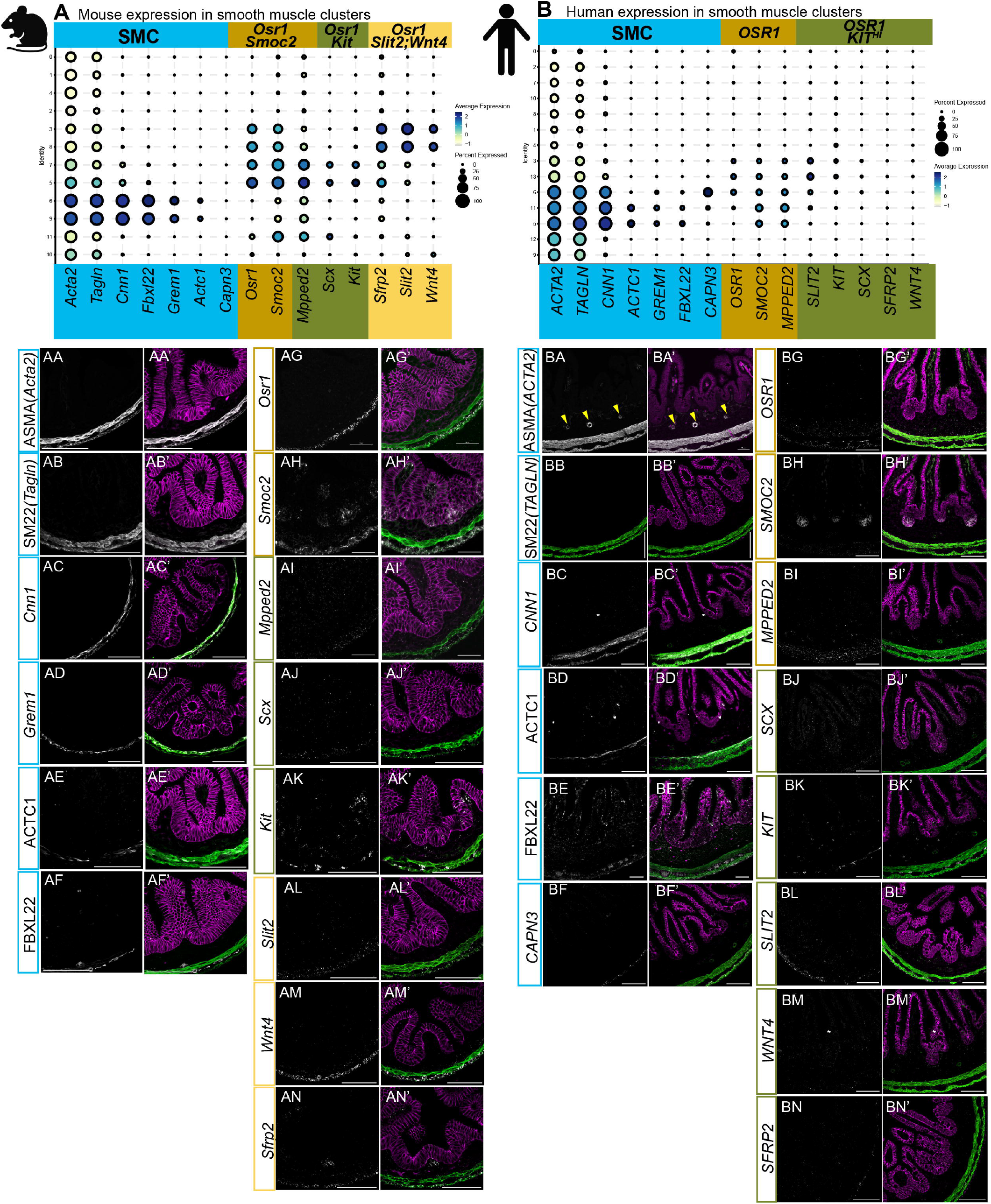
Smooth muscle cell type markers and their RNA transcript or protein localization. A,B) Dotplot of average transcript level in percent of cells per cluster in mouse (A) or human (B). AA-AN, BA-BN) RNA localization of indicated transcripts or proteins (ASMA, SM22, FBXL22 and ACTC1) at E16.5 in mouse (AA-AN) or human at 80d (BA-BN). AA’-AN’, BA’-BN’) same tissue sections as in AA-AN or BA’-BN’, respectively but overlaid with anti-ECADHERIN (magenta) and anti-SM22/TAGLIN (green). Background boxes highlight groups of smooth muscle cell type markers: blue are smooth muscle cells, gold are Osr1+/OSR1+ precursors, olive are Osr1+/OSR1+;Kit+/KIT^HI^; yellow are Osr+;Slit2+;Wnt4+. Scalebars are 50um.

**Figure 5.**
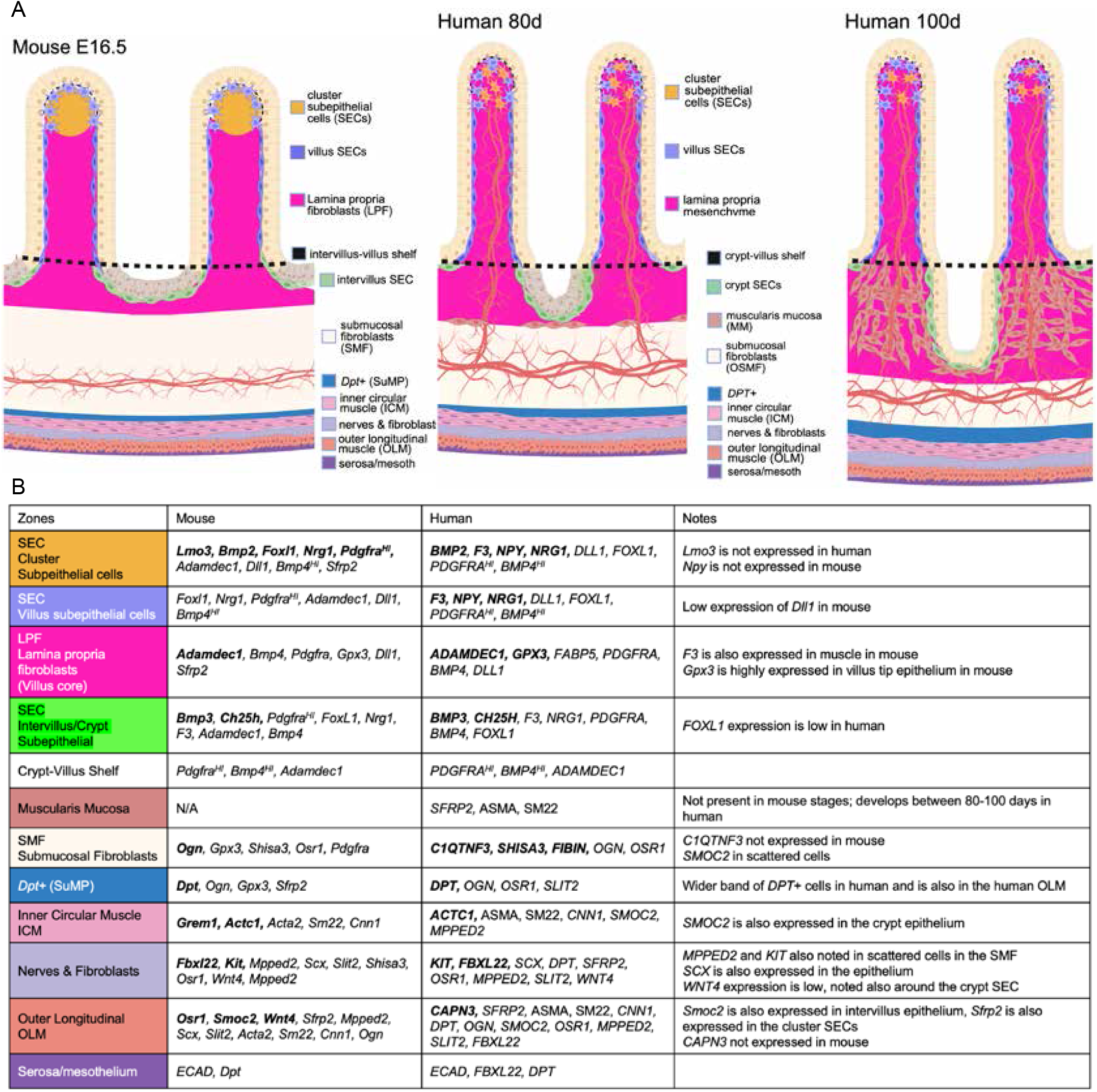
Summary schematic of radial patterning of mesenchymal and smooth muscle cell types and their markers in mouse at E16.5 and in human at 80 and 100dpc. Color coding matches the chart at the bottom summarizing marker localization.

The newly identified SEC markers are *Lmo3* (mouse only), *Bmp2/BMP2, Bmp3/BMP3*, and *Ch25h/CH25H*. Importantly, localization of these transcripts also showed that human SECs could be further divided into either villus localized (*BMP2*, **Figure 3BF,BF’ and Supplemental Figure 4F**,**J**,**J’**) or crypt-localized (*BMP3*, **Figure 3BG, BG’** and magnifications in **Supplemental Figure 4G, K, K’**; *CH25H*, **Figure 3BH, BH’ and magnification in Supplemental Figure 4**,**H**). In the mouse telocytes/SECs these transcripts were expressed all along the epithelium though *Bmp2* showed higher expression at the tips of the villi (**Figure 3AF, AF’ and magnification in Supplemental Figure 4B**) and *Ch25h* showed higher expression in cells along the intervillus epithelium (**Figure 3AH, AH’** and magnification in Supplemental Figure 4D). *Lmo3* is specifically expressed in a subset of mouse SECs at the tips of the villi (**Figure 3AE, AE’ and magnification in Supplemental Figure 4A**) but is not expressed in human SECs (**Figure 3B**). Finally, *F3* is a strong and specific marker of human SECs^28^ (**Figure 4B, BD, BD’**) and we that found it is also highly expressed in mouse SECs, though it is also highly expressed in mouse SMCs (**Figure 3A, c6,9**).

*BMP4* was found to be expressed in human SECs but is also expressed in other cell types at varying levels, similar to *Pdgfra/PDGFRA* expression (**Figure 3A, B**). By imaging, the cells with high expression of *BMP4* are easy to distinguish and identify as SECs in the human tissue (**Figure 3BJ, BJ’**). In the mouse however, *Bmp4* was evenly expressed in not only the telocytes/SECs but other mesenchymal cells in the villus core (**Figure 3AJ, AJ’**). *Adamdec1* can be visualized in both mouse telocytes/SECs and some LPFs (**Figure 3A, AK, AK’**). *ADAMDEC1;* however, is not observed in human SECs but is instead expressed in the LPFs of the villus core mesenchyme (**Figure 3B, BK, BK’**). Finally, *Npy* is expressed in very few cells in the mouse (**Figure 3A**), however in the human, consistent with previous findings^28^, *NPY* is a strong and specific marker of SECs (**Figure 3B, BE, BE’**).

One of the most specific markers in the human LPFs is *ADAMDEC1* (**Figure 4B, BK, BK’ and Supplemental Figure 4L, L’**). However, as noted above, mouse *Adamdec1* was found to be expressed highly in both SECs and LPFs (**Figure 3A,AK,AK’**). As seen previously^28^, *GPX3* expression in the human is confined to the LPFs (**Figure 3BL,BL’**). In the mouse, *Gpx3* expression extends beyond the villus core into the submucosal mesenchyme, marking the intestinal fibroblasts all the way to the smooth muscle layers, and is also highly expressed in the villus tip epithelium (**Figure 3AL, AL’**) making it a poor marker for mouse LPFs. *FABP5*, while highly expressed in human LPFs, it is not specific to the human LPFs and is also observed in the epithelium and pericytes (**Figure 3B, BM, BM’**). We did not examine *Fabp5* localization in the mouse as we had noted wide expression in many cell types with notably low expression in mouse LPF clusters (**Figure 2C**).

The SMFs were found between the muscularis mucosa near the bottom of the epithelium and the longitudinal and circular muscular layers near the outermost edge of the intestine. SMFs of both the mouse and human were found to express *Shisa3/SHISA3* and *Ogn/OGN*. In the mouse, *Shisa3* is also expressed by a few cells of the outer longitudinal muscle (OLM)(**Figure 3, AN, AN’**) while human *SHISA3* is specific to SMFs (**Figure 3BO, BO’**). In both species, *Ogn/OGN* marks the OLM in addition to the SMFs but is notably absent from the ICM (**Figure 3A, AM, AM’ and BN, BN’**). Interestingly, mouse expression of *Shisa3* and *Ogn* are both noticeably higher in the single layer of SMFs immediately adjacent to the inner circular muscle (ICM). This single-cell-thick band was recently described^57^ as the superficial muscularis propria (SuMP) in post-natal mouse intestine and is also marked in the mouse by *Dpt* (**Figure 3A, AO, AO’**) while in the human the *DPT* band matches a broad expression of the other human SMF markers (**Figure 4B, BP, BP’**).

In human tissue, SMFs also express *C1QTNF3* which exhibits a pattern similar to *OGN, DPT*, and *SHISA3* (**Figure 3B,BQ,BQ’**), though it is not expressed in the smooth muscle layers making it a more specific SMF marker than *OGN* and *DPT*. We also found that *C1qtnf3* is not expressed in the mouse (**Figure 3A**). Surprisingly, while the SMF markers were often expressed in smooth muscle, this was always specific to the OLM, and these markers were entirely absent from ICM.

In the smooth muscle populations, we found several well-defined markers^57^ such as alpha smooth muscle actin (ASMA*/ACTA2*; **Figure 5AA, AA’, BA, BA’**), SM22*/TAGLN* (**Figure 5AB, AB’, BB, BB’**) and *Cnn1/CNN1* (**Figure 5AC, AC’, BC, BC’**) are expressed and localized to the ICM and OLM layers in both mouse and human. However, human ASMA also showed expression in the vascular-associated smooth muscle cells (**Figure 3BA, BA’, yellow arrowheads**).

The current literature does not officially define a unique transcriptomic identity for the ICM and OLM (or the MM that develops later) and instead relies primarily on visual localization using imaging. Through our comparative analyses, we have identified a few markers that are unique to one sub-population or the other. *Grem1* in the mouse is expressed in just the ICM (**Figure 4AE, AE’**). ACTC1 protein is localized in the ICM and in cells between the two muscle layers in both mouse and human (**Figure 4AE, AE’, BD, BD’**) while FBXL22 protein is expressed in the OLM and in cells between the muscle layers (**Figure 4AF, AF and BE, BE’**). Finally, *CAPN3* is expressed in just the OLM in human (**Figure 4BF, BF’**) and is not expressed in mouse between E14.5-E17.5 (**Figure 4A**). Overall, we identified markers specific to the mouse ICM, both the mouse and human ICM and the layer of fibroblasts between the ICM and the OLM, and finally the human OLM.

In both human and mouse small intestinal mesenchyme, we also identified an *Osr1/OSR1*+ mesenchyme population. Using immunofluorescence and FISH, we found that these cells are defined by low expression of SMC markers Smooth Muscle Actin (SMA – gene name *ACTA2*) and SM22 (gene name *TAGLN*) in addition to expression of *Osr1+/Smoc2+* (mouse) or *OSR1+/SMOC2+/MPPED2+* (human)(**Figure 4A, B**).

While in single-cell data we observed *Osr1/OSR1* expression in several clusters (Figure 4A (c)3,5,7,8 and 4B (c)9 3,6,13), *Osr1* localization at E16 in the mouse is observed in the OLM (**Figure 4AG, AG’**). In the human, *OSR1* is also localized to the OLM, but it is also expressed in the SMF and LPF domains (**Figure 4BG, BG’**). Mouse *Smoc2* is highly expressed in the OLM, it is also expressed at lower levels in the ICM, intervillus epithelium and in the villus tip SECs (**Figure 4AH, AH’**). *SMOC2* in the human shows a slightly different localization pattern to the mouse with expression in both the ICM and OLM but the strongest expression is in the crypt epithelium and it is not detected in the villus tip (**Figure 4BH, BH’**).

In the mouse, *Mpped2* is expressed in the mouse ICM and OLM with stronger expression in the OLM (**Figure 4AI, AI’, AJ, AJ’**). *MPPED2* is also localized to the human ICM and OLM in human but is also observed in the SMF layer, likely in VSMCs (**Figure 4BI, BI’**). Scx/*SCX* is expressed in the cells between the muscle layers and the OLM but not the ICM in both species while the *SCX* is also expressed throughout the *Hs*-epithelium (**Figure 4BJ, BJ’**). *Kit/KIT* is a known marker for immune cells^58^, interstitial cells of Cajal^59^ and ICC-precursors^60^ which are in close association with the smooth muscle layers and ENS. We observe Kit/KIT localization to cells between the muscle layers as well as cells in the mesenchyme that are likely immune cells (**Figure 5AK, AK’, BK, BK’**).

Another *Osr1*+ mesenchyme population in the mouse expresses *Slit2, Wnt4 and Sfrp2. Slit2* and *Wnt4* are highly expressed in the *Mm*-OLM and cells between the muscle layers in mouse (**Figure 4AL, AL’ and AM, AM’)**. *SLIT2* is expressed in the *Hs*-OLM and some SM22^**+**^ cells between the muscle layers (**Figure 4BL-BL’**). *WNT4* does not show significant staining distinguishable from background in the human tissue at 80dpc. *Sfrp2* is observed in the mouse OLM and SECs of the mouse intestine (**Figure 4AN, AN’**). *SFRP2* is expressed by interspersed cells in the human OLM at very low levels, specifically in some, but not all, of the SM22+ cells that bridge and connect the two muscle layers (**Figure 4BN, BN’**). Unlike in the mouse, *SFRP2* does not appear to be expressed within the human SECs.

## Discussion

In this study, we performed a detailed cross-species comparison of the developing intestinal mesenchyme, leveraging single-cell transcriptomics and spatial localization techniques to build an atlas of fibroblast and smooth muscle populations in mouse and human. Our analysis reveals a high degree of conservation in the major mesenchymal cell types and their spatial organization along the radial axis of the gut tube. However, this broad similarity is nuanced by significant divergence at the level of individual gene expression, where key lineage-defining markers in one species are often expressed differently or not at all in the analogous cell population of the other. These findings underscore both the utility of the mouse as a model for human intestinal development and the critical importance of validating gene expression patterns across species before drawing direct functional comparisons. Based on the regional pattern of these markers, we have defined the radial domains, and their key maker genes, as summarized in **Figure 5A**. We define broad populations of fibroblasts (SECs, LPFs and SMFs) based on their transcriptomes and identify markers for these cell types in the mouse and human that map the localization of these cell types to similar radial domains during early intestinal development. Many of the cell type markers are shared, however, important species-specific differences including some novel markers were noted (**Figure 5B**).

One caveat to this study is that mouse SECs only represented 1.3% of all mesenchymal cells while human SECs were 8.7% of all mesenchymal cells. Since the mouse had fewer SECs, they did not form a distinct cluster separate from the LPFs as they do in human. In fact, the human LPF specific marker, ADAMDEC1, is expressed in mouse in SECs and LPFs. These shared transcriptional similarities and the fact that the average of the 30 nearest neighbors shows that human SECs are more likely to match mouse LPFs, suggest that LPFs may give rise to the SECs. Many SEC markers were shared between species (*Pdgfra/PDGFRA*^*HI*^, *Bmp4/BMP4*^*HI*^ and *Nrg1/NRG1*), but some were species-specific (mouse *Lmo3*, human *NPY*) or were a good marker in one species (human *F3, DLL1, GPX3;* mouse *Foxl1*) but were not cell type specific in the other. We also discovered SEC markers that specifically define sub-domains of localization: villus tip (*Mm*-*Lmo3* and *Bmp2/BMP2*) versus intervillus/crypt (*Bmp3/BMP3* and *Ch25h/CH25H*).

A key contribution of our work is the refinement of molecular markers for these conserved mesenchymal populations and the identification of novel, spatially distinct subpopulations. Our findings also raise questions about mesenchymal lineage relationships. In our label transfer data (**Figure 2**) human LPFs mismatched most commonly with mouse SMFs, and human SMFs (when they did not match mouse SMFs) mismatched about equally with LPFs and *Osr1*+ mesenchyme. On the other hand, mouse LPFs only mapped onto human LPFs 10.6% percent of the time, and most of the mouse LPFs mismatched with human SMFs. These consistent mismatches (LPF to SMF and vice versa) suggest that fibroblast subsets may exist along developmental continua rather than as fixed lineages. The OSR1+ mesenchyme emerges as a potential progenitor pool, capable of contributing to SMFs and SMCs, consistent with its broad transcriptional overlap and proliferative state. Future lineage tracing and functional studies will be necessary to test these hypotheses and to define how mesenchymal progenitors contribute to villus morphogenesis, crypt formation and smooth muscle compartmentalization.

One of the primary challenges in this work was the bioinformatic complexity of cross-species single-cell analysis. The lack of standardized tools for gene homolog conversion required us to develop a custom pipeline to handle one-to-one, one-to-many, and many-to-one homolog relationships, a crucial step for accurate data integration. As cross-species comparisons become more common, the development of robust, maintained bioinformatic tools will be essential.

## Acknowledgements

This work was supported by the NIH National Institute of Diabetes and Digestive and Kidney Diseases (NIDDK) R01KD121166 to K.D.W. This project has also been made possible in part by grant numbers 2019-002440 (Seed Network) and 2021-237566 (Pediatric Network) from the Chan Zuckerberg Initiative DAF, an advised fund of Silicon Valley Community Foundation to J.R.S. J.R.S. was also supported and by the NIDDK (R01DK137806 and RC2DK140862), and by the Intestinal Stem Cell Consortium (U01DK103141), a collaborative research project funded by NIDDK and the National Institute of Allergy and Infectious Diseases (NIAID). K.J. is supported by NIDDK F32DK138694 and I.G. and the University of Washington Laboratory of Developmental Biology were supported by NIH award R24HD000836 from the Eunice Kennedy Shriver National Institute of Child Health and Human Development (NICHD).

The authors are thankful for technical support from Tristan Reed, Sophia Liang, Jehoon Seo and the University of Michigan Unit for Laboratory Animal Medicine.

## Author Contributions

Conceptualization K.D.W., Z.X., J.R.S.; Methodology K.D.W., Z.X., X.D., J.R.S., K.J.; Software Z.X., X.D.; Validation K.D.W., H.I., M.A.; Formal Analysis K.D.W., X.D., Z.X., K.J.; Investigation K.D.W., X.D., Z.X., H.I.; Resources K.D.W., Z.X., X.D., I.G., J.R.S.; Data Curation K.D.W.; Writing K.D.W., K.J., Z.X., X.D.; Visualization K.D.W.; X.D., Z.X.,; Supervision, K.D.W., J.R.S.; Funding Acquisition K.D.W., J.R.S.

## Methods

### Mice

C57BL/6 mice purchased from Jackson Labs were housed and handled humanely according to the Guide for the Care and use of Laboratory Animal standards overseen by the University of Michigan Unit for Laboratory Animal Medicine and the University’s IACUC committee. Timed pregnancies were determined by post-coital plug and fetuses were staged according to the Theiler criteria [https://www.emouseatlas.org/emap/ema/theiler_stages/downloads/theiler2.pdf] at the time of tissue harvest.

### Human tissue

Normal human fetal intestinal tissue was obtained from the University of Washington Laboratory of Developmental Biology. All human tissue was de-identified and all experiments were conducted with approval from the University of Michigan IRB.

### Tissue collection and preparation of RNA libraries for Single-Cell Sequencing and transcript alignment

Whole mouse fetal intestines (from just posterior to the common bile duct through just anterior of the cecum) were collected and pooled for each stage E13.5, E14.5, E15.5, E16 and E17.5 and then dissociated to single cells using the Miltenyi Neural Tissue dissociation kit and single-cell libraries were prepared on the 10x Chromium by the University of Michigan Advanced Genomics core with a target of 5000 cells as previously described^28^. All single-cell RNA-seq sample libraries were prepared with the 10x Chromium Controller using v3 chemistry. Sequencing was performed on an Illumina NovaSeq 6000 with targeted depth of 100,000 reads per cell. Default alignment parameters were used to align reads to the hg19 human reference genome or the mm38 mouse reference genome provided by the 10X CellRanger pipeline. Initial cell demultiplexing and gene quantification were also performed with the default 10x Cell Ranger pipeline. Human fetal intestinal scRNA sequences used for comparisons are from Holloway^28^, Elmentaite^29^, Yu^36^ and Hung^35^.

### Tissue preparation for FISH, immunostaining and imaging

Human and mouse fetal intestines were prepared as described previously^28^. Briefly, 1 cm tissue segments were fixed for 24 hours at room temperature in 10% neutral buffered formalin. After fixation, tissues were washed in RNase free water and then dehydrated through a methanol series prior to paraffin embedding. Five um thick sections were prepared and used for FISH staining according to the ACD manufacturer’s protocol and co-immunostaining was performed as previously described^28^. Probes and antibodies used are listed in **STAR METHODS**. Imaging was performed on a Nikon AXR confocal. Images were assembled in ImageJ. All parameters were kept consistent within an experimental set.

### Bioinformatics/scRNA-seq

#### Overview

To identify and visualize distinct cell populations within the single-cell RNA sequencing datasets, we employed the workflow outlined by the Seurat 4.0 R package^61^. This pipeline includes: filtering cells for quality control; applying the SCTransform technique^62^; reducing dimensionality with principal component analysis (PCA) and uniform manifold approximation and projection (UMAP)^63^; clustering by either the Louvain algorithm^64^ or the Leiden algorithm^65^; log normalization for gene expression visualization and for differential gene expression analysis; label transfer and reference-based UMAP through

Seurat^61^ to compare the heterogeneity of cell types identified in human and mouse intestinal mesenchyme.

#### Quality control of individual samples

The integrated mouse dataset (**Figure 1A-B**) consists of 5 samples (embryonic days E13.5, E14.5, E15.5, E16 and E17.5) and the integrated human dataset (**Figure 1D-E**) consists of 14 samples (42.7dpc, 47dpc Duodenum, Jejunum, Ileum, 58.8dpc, 59dpc Duodenum, Jejunum, Ileum, 64.4dpc, 69.3dpc, 70dpc, 70dpc Duodenum, 72dpc Duodenum, 80dpc Duodenum, 80dpc Ileum, 80dpc Jejunum, 101dpc Duodenum, 101dpc Ileum) where samples 42.7dpc, 58.8dpc, 64.4dpc, 69.3dpc, 70dpc are transformed into Seurat objects from deposited H5AD objects. To ensure quality of the data, all individual samples were first filtered to remove cells expressing too few or too many genes, too low or to high UMI counts according to Supplemental Table 2 or a fraction of mitochondrial genes greater than (0.1 for human samples, 0.05 for mouse samples). The final step was clustering individual samples and removing blood cells clusters before integration using canonical markers such as *HBB*.

#### Sample integration by species, clustering and extraction

After quality control, the standard pre-processing workflow includes normalization, scaling, and selection of highly variable features. After independently pre-processing the data, we then used Seurat’s SCTransform^62,66^. During the SCTransform process, we also chose to regress out a confounding source of variation – mitochondrial mapping percentage. Considering the different time points of our samples, we utilized either SCTransform integration (**Figure 2A-B**) or fastMNN^67^(**Figure 1A-B, D-E; Figure 2D-E, Supplemental Figure 2A-B, D-E;**), an efficient batch correction method that rely on the identification of mutual nearest neighbors (MNNs) in high-dimensional expression space based on their individual performance. After completion batch correction with the default settings, the influence of batch specific technical artifacts on clustering is reduced.

Specifically, for the epithelium extraction of mouse intestine samples (**Supplemental Figure 2B**), we used canonical markers such as *Epcam* and *Cdh1* to identify, extract, and re-cluster candidate clusters 1,10,11,16,18 (**Figure 1B**); for the epithelium extraction of human intestine samples (**Supplemental Figure 2E**), similarly, we identified, extracted, and re-clustered candidate clusters 1,4,13 (**Figure 1E**); for the mesenchymal extraction of mouse intestine samples (**Figure 2B**), we used canonical markers including *Vim* and *Col1a1* to identify, extract, and re-cluster candidate clusters 2,3,5,8,9 from **Figure 1B**, and we subsequently removed prominent proliferative subclusters (i.e., *Top2a*+ and *Mki67*+); for the mesenchyme extraction of human intestine samples (**Figure 2E**), similarly, we identified, extracted, and re-clustered candidate clusters 0,2,3,6,8,11 from (**Figure 1E**).

#### Normalization for visualization and differential gene expression

As recommended by Seurat developers, we employed the method of log normalization on the standard RNA assay to graph dot plots, feature plots, and conduct DGE analysis. Expression matrix read counts per cell were normalized by the total expression, multiplied by a scale factor of 10000, and finally log-transformed. For the differential gene expression testing, we only tested features that are first, detected in a minimum fraction of 0.25 in either of the two cell populations, and second, show at least 0.25-fold difference in log-scale between the two cell clusters on average.

#### Translating features cross-species by gene homology

Before integrating the mouse and human datasets, the gene names of the mouse features were converted to their human homologs, and each dataset was clustered based on the transcriptional profile of each cell. Cell clusters were annotated using the expression of markers identified pre-integration (**Figure 1**).

We used the Mouse Genome Information (MGI) database maintained by The Jackson Laboratory^68^. After downloading the complete homology list (HOM_MouseHumanSequence.rpt) comparing mouse and human, we applied several sub-setting steps to the features for further quality control based on the following criteria: 1) genes with 1-to-1 homologs, 2) genes with one mouse homolog to many human homologs (1-to-many), 3) genes with many mouse homologs to 1 human homolog (many-to-1) and 4) genes present in the mouse data with no defined homologs in the human data. Sub-setting was performed on the basis of the unique homolog ID number (DB.Class.Key) and species. If a feature had DB.Class.Key = 2 with two different species it was subset as a 1-to-1 feature. If DB.Class.Key > 2 with unique species = 2 it was subset as either many-to-1 or 1-to-many depending on which species was more common. Features with DB.Class.Key < 2 and species = mouse were removed from the analysis because our analysis was human-centric, so mouse genes without a human ortholog were removed from downstream analysis but human genes with no mouse homolog were left in the matrix. Genes with 1-to-1 homologs were further subset for features where the mouse and human homologs were exactly the same (ex. *F3/F3*), the features were subset into their own matrix and the homology algorithm was not applied because the feature names were already the same.

For genes that were 1-to-many, in addition to re-naming the feature(s), the mouse feature information was ‘duplicated’ to fill all human homologs while genes that were many-to-1 were combined additively into the single human homolog. After homology, the four matrices were combined to generate the complete, integrated converted mouse dataset with all gene names corresponding to the human homolog(s). The converted mouse dataset and integrated human datasets were merged (50486 genes in the final object), then we utilized the Harmony^69^ integration algorithm embedded in Seurat v5, which is a method for integrating single-cell RNA seq data across multiple datasets by adjusting for batch effects while preserving biological variation using a fast, scalable approach based on the optimization of a latent variable model.

We performed validation of the species integration using well-defined cell class markers known to be shared across both species. The data was then re-split, re-visualized (UMAP), and re-annotated to more closely examine all cells using Louvain clustering algorithm To further examine the mesenchymal cell types post-integration, we extracted the human and mouse mesoderm-derived mesenchyme from the complete datasets post-homology translation and. The mesenchymal data was then re-visualized (UMAP) and re-annotated using the previously defined markers.

#### Reference based mapping and calculation of population fidelity between species

To quantitatively compare the average similarity of mouse cell populations to the human, we projected all mouse cells that underwent homology translation onto the human reference UMAP and vice versa (**Figure 2**). We utilized Seurat’s recommended pipeline to perform single-cell reference mapping. PCAs were first performed on reference and query data. Then a set of anchors were identified based on the default setting of the function FindTransferAnchors, 200 neighbors were used when filtering anchors. With the computed anchors, reference.reduction parameter was set to PCA and reduction.model was set to UMAP, the function MapQuery returned the projected UMAP coordinates of the query cells mapped onto the reference UMAP. We then integrated the projected UMAP (pseudo-color) and the reference UMAP (light gray) to visualize the result of our reference-based mapping in 2D. To quantify the similarity in cell annotations between species, we extracted the 30 nearest neighbors (NN) matrix from the Seurat object of reference cells used to map its projected location by quantifying the distribution of original cell annotations of the 30 NNs of the reference data. That is, of the 30 NNs to cell X, we quantified the number of those 30 that were of each reference cell type. If 28/30 (α = 0.067) of the reference cell IDs of the NNs matched the original, annotated ID of the query cell, it was considered a ‘match’.The number of query cells of each cell type that matched or mismatched was converted to a percent for visualization. The distribution of matched cell types shows the average composition of the 30 NNs for each cell type. This gave us a quantitative measure of how similar or different the individual mouse populations were on average to the corresponding human population and vice versa based on the entire transcriptome of each cell type as well as identifying the composition of the NNs of each population.

**Supplemental Figure 1. Comparison of single-cell sequencing of all mouse cells and all human cells**. A,F) UMAP rendering of mouse or human samples and B,G) by transcriptional similarity. C,H) Dotplot of average expression and percent of cells expressing transcripts in each mouse or human cluster. D,I) Parts of the whole distribution plot of the contribution each mouse or human cluster per stage for all cells. E,J) Heat map of the percent of the sample contributing to a mouse or human cluster at each stage for all cells.

**Supplemental Figure 2. Comparison of single-cell sequencing of mouse and human epithelial cells**. A,F) UMAP rendering of mouse or human epithelial cells by sample and B,G) by transcriptional similarity. C,H) Dotplot of the average expression of transcripts in each mouse or human epithelial cluster. D,I) Parts of the whole distribution plot of the contribution each mouse or human cluster per stage for epithelial cells. E,J) Heat map of the percent of the sample contributing to a mouse or human cluster at each stage for epithelial cells.

**Supplemental Figure 3. Mouse LPF cluster extracted and re-clustered reveals a small, but distinct SEC sub-cluster**. A) UMAP rendering of LPF cells with 5 sub-clusters. B) Dotplot of proliferation markers (gray), LPF markers (orange) and SEC markers (teal).

**Supplemental Figure 4. Magnifications of localization of telocyte/SEC markers (A-H, I-K’) and 100dpc localization of LPF and SMF markers (L-O’)**.

**Supplemental Table 1. Transcripts from genes in the mouse that were not identified in the human dataset**.

**Supplemental Table 2. Filters applied per sample to remove cells expressing too few or too many genes, too low or too high UMI counts**.

## References

1. Yu, Q., Kilik, U., Secchia, S., Adam, L., Tsai, Y.H., Fauci, C., Janssens, J., Childs, C.J., Walton, K.D., Lopez-Sandoval, R., et al. (2025). Recent evolution of the developing human intestine affects metabolic and barrier functions. Science, eadr8628. 10.1126/science.adr8628.

2. Spence, J.R., Lauf, R., and Shroyer, N.F. (2011). Vertebrate intestinal endoderm development. Dev Dyn 240, 501–520. 10.1002/dvdy.22540.

3. Sadler, T.W. (2024). Langman’s medical embryology, Fifteenth edition. Edition (Wolters Kluwer).

4. Lewis, S.L., and Tam, P.P. (2006). Definitive endoderm of the mouse embryo: formation, cell fates, and morphogenetic function. Dev Dyn 235, 2315–2329. 10.1002/dvdy.20846.

5. Thompson, C.A., DeLaForest, A., and Battle, M.A. (2018). Patterning the gastrointestinal epithelium to confer regional-specific functions. Dev Biol 435, 97–108. 10.1016/j.ydbio.2018.01.006.

6. Zwick, R.K., Kasparek, P., Palikuqi, B., Viragova, S., Weichselbaum, L., McGinnis, C.S., McKinley, K.L., Rathnayake, A., Vaka, D., Nguyen, V., et al. (2024). Epithelial zonation along the mouse and human small intestine defines five discrete metabolic domains. Nature Cell Biology. 10.1038/s41556-023-01337-z.

7. Heanue, T.A., and Pachnis, V. (2007). Enteric nervous system development and Hirschsprung’s disease: advances in genetic and stem cell studies. Nat Rev Neurosci 8, 466–479. 10.1038/nrn2137.

8. Lake, J.I., and Heuckeroth, R.O. (2013). Enteric nervous system development: migration, differentiation, and disease. Am J Physiol Gastrointest Liver Physiol 305, G1–24. 10.1152/ajpgi.00452.2012.

9. Li, X., Madison, B.B., Zacharias, W., Kolterud, A., States, D., and Gumucio, D.L. (2007). Deconvoluting the intestine: molecular evidence for a major role of the mesenchyme in the modulation of signaling cross talk. Physiological Genomics 29, 290–301. 10.1152/physiolgenomics.00269.2006.

10. Le Guen, L., Marchal, S., Faure, S., and de Santa Barbara, P. (2015). Mesenchymal-epithelial interactions during digestive tract development and epithelial stem cell regeneration. Cell Mol Life Sci 72, 3883–3896. 10.1007/s00018-015-1975-2.

11. Shoshkes-Carmel, M., Wang, Y.J., Wangensteen, K.J., Toth, B., Kondo, A., Massasa, E.E., Itzkovitz, S., and Kaestner, K.H. (2018). Subepithelial telocytes are an important source of Wnts that supports intestinal crypts. Nature 557, 242–246. 10.1038/s41586-018-0084-4.

12. Walton, K.D., Kolterud, A., Czerwinski, M.J., Bell, M.J., Prakash, A., Kushwaha, J., Grosse, A.S., Schnell, S., and Gumucio, D.L. (2012). Hedgehog-responsive mesenchymal clusters direct patterning and emergence of intestinal villi. Proc Natl Acad Sci U S A 109, 15817–15822. 10.1073/pnas.1205669109.

13. Walton, K.D., Whidden, M., Kolterud, A., Shoffner, S.K., Czerwinski, M.J., Kushwaha, J., Parmar, N., Chandhrasekhar, D., Freddo, A.M., Schnell, S., and Gumucio, D.L. (2016). Villification in the mouse: Bmp signals control intestinal villus patterning. Development 143, 427–436. 10.1242/dev.130112.

14. Cervantes, S., Yamaguchi, T.P., and Hebrok, M. (2009). Wnt5a is essential for intestinal elongation in mice. Dev Biol 326, 285–294. 10.1016/j.ydbio.2008.11.020.

15. Zhang, X., Stappenbeck, T.S., White, A.C., Lavine, K.J., Gordon, J.I., and Ornitz, D.M. (2006). Reciprocal epithelial-mesenchymal FGF signaling is required for cecal development. Development 133, 173–180. 10.1242/dev.02175.

16. Karlsson, L., Lindahl, P., Heath, J.K., and Betsholtz, C. (2000). Abnormal gastrointestinal development in PDGF-A and PDGFR-(alpha) deficient mice implicates a novel mesenchymal structure with putative instructive properties in villus morphogenesis. Development 127, 3457–3466.

17. Kolterud, A., Grosse, A.S., Zacharias, W.J., Walton, K., Kretovich, K.E., Madison, B.B., Waghray, M., Ferris, J.E., Hu, C., Merchant, J.L., et al. (2009). Paracrine Hedgehog signaling in stomach and intestine: new roles for hedgehog in gastrointestinal patterning. Gastroenterology 137, 618–628. 10.1053/j.gastro.2009.05.002.

18. Grun, D., Lyubimova, A., Kester, L., Wiebrands, K., Basak, O., Sasaki, N., Clevers, H., and van Oudenaarden, A. (2015). Single-cell messenger RNA sequencing reveals rare intestinal cell types. Nature 525, 251–255. 10.1038/nature14966.

19. McCarthy, N., Manieri, E., Storm, E.E., Saadatpour, A., Luoma, A.M., Kapoor, V.N., Madha, S., Gaynor, L.T., Cox, C., Keerthivasan, S., et al. (2020). Distinct Mesenchymal Cell Populations Generate the Essential Intestinal BMP Signaling Gradient. Cell Stem Cell 26, 391–402 e395. 10.1016/j.stem.2020.01.008.

20. Zhao, L., Song, W., and Chen, Y.G. (2022). Mesenchymal-epithelial interaction regulates gastrointestinal tract development in mouse embryos. Cell Rep 40, 111053. 10.1016/j.celrep.2022.111053.

21. Paerregaard, S.I., Wulff, L., Schussek, S., Niss, K., Morbe, U., Jendholm, J., Wendland, K., Andrusaite, A.T., Brulois, K.F., Nibbs, R.J.B., et al. (2023). The small and large intestine contain related mesenchymal subsets that derive from embryonic Gli1(+) precursors. Nat Commun 14, 2307. 10.1038/s41467-023-37952-5.

22. Fazilaty, H., Brugger, M.D., Valenta, T., Szczerba, B.M., Berkova, L., Doumpas, N., Hausmann, G., Scharl, M., and Basler, K. (2021). Tracing colonic embryonic transcriptional profiles and their reactivation upon intestinal damage. Cell Rep 36, 109484. 10.1016/j.celrep.2021.109484.

23. Bahar Halpern, K., Massalha, H., Zwick, R.K., Moor, A.E., Castillo-Azofeifa, D., Rozenberg, M., Farack, L., Egozi, A., Miller, D.R., Averbukh, I., et al. (2020). Lgr5+ telocytes are a signaling source at the intestinal villus tip. Nat Commun 11, 1936. 10.1038/s41467-020-15714-x.

24. Hong, S.P., Yang, M.J., Cho, H., Park, I., Bae, H., Choe, K., Suh, S.H., Adams, R.H., Alitalo, K., Lim, D., and Koh, G.Y. (2020). Distinct fibroblast subsets regulate lacteal integrity through YAP/TAZ-induced VEGF-C in intestinal villi. Nat Commun 11, 4102. 10.1038/s41467-020-17886-y.

25. Burclaff, J., Bliton, R.J., Breau, K.A., Ok, M.T., Gomez-Martinez, I., Ranek, J.S., Bhatt, A.P., Purvis, J.E., Woosley, J.T., and Magness, S.T. (2022). A Proximal-to-Distal Survey of Healthy Adult Human Small Intestine and Colon Epithelium by Single-Cell Transcriptomics. Cell Mol Gastroenterol Hepatol 13, 1554–1589. 10.1016/j.jcmgh.2022.02.007.

26. Gao, S., Yan, L., Wang, R., Li, J., Yong, J., Zhou, X., Wei, Y., Wu, X., Wang, X., Fan, X., et al. (2018). Tracing the temporal-spatial transcriptome landscapes of the human fetal digestive tract using single-cell RNA-sequencing. Nat Cell Biol 20, 721–734. 10.1038/s41556-018-0105-4.

27. Kinchen, J., Chen, H.H., Parikh, K., Antanaviciute, A., Jagielowicz, M., Fawkner-Corbett, D., Ashley, N., Cubitt, L., Mellado-Gomez, E., Attar, M., et al. (2018). Structural Remodeling of the Human Colonic Mesenchyme in Inflammatory Bowel Disease. Cell 175, 372–386 e317. 10.1016/j.cell.2018.08.067.

28. Holloway, E.M., Czerwinski, M., Tsai, Y.H., Wu, J.H., Wu, A., Childs, C.J., Walton, K.D., Sweet, C.W., Yu, Q., Glass, I., et al. (2021). Mapping Development of the Human Intestinal Niche at Single-Cell Resolution. Cell Stem Cell 28, 568–580 e564. 10.1016/j.stem.2020.11.008.

29. Elmentaite, R., Ross, A.D.B., Roberts, K., James, K.R., Ortmann, D., Gomes, T., Nayak, K., Tuck, L., Pritchard, S., Bayraktar, O.A., et al. (2020). Single-Cell Sequencing of Developing Human Gut Reveals Transcriptional Links to Childhood Crohn’s Disease. Developmental cell 55, 771–783 e775. 10.1016/j.devcel.2020.11.010.

30. Hickey, J.W., Becker, W.R., Nevins, S.A., Horning, A., Perez, A.E., Zhu, C., Zhu, B., Wei, B., Chiu, R., Chen, D.C., et al. (2023). Organization of the human intestine at single-cell resolution. Nature 619, 572–584. 10.1038/s41586-023-05915-x.

31. Elmentaite, R., Kumasaka, N., Roberts, K., Fleming, A., Dann, E., King, H.W., Kleshchevnikov, V., Dabrowska, M., Pritchard, S., Bolt, L., et al. (2021). Cells of the human intestinal tract mapped across space and time. Nature 597, 250–255. 10.1038/s41586-021-03852-1.

32. Fawkner-Corbett, D., Antanaviciute, A., Parikh, K., Jagielowicz, M., Gerós, A.S., Gupta, T., Ashley, N., Khamis, D., Fowler, D., Morrissey, E., et al. (2021). Spatiotemporal analysis of human intestinal development at single-cell resolution. Cell 184, 810-+. 10.1016/j.cell.2020.12.016.

33. Cao, J., O’Day, D.R., Pliner, H.A., Kingsley, P.D., Deng, M., Daza, R.M., Zager, M.A., Aldinger, K.A., Blecher-Gonen, R., Zhang, F., et al. (2020). A human cell atlas of fetal gene expression. Science 370. 10.1126/science.aba7721.

34. Zwick, R.K., Kasparek, P., Palikuqi, B., Viragova, S., Weichselbaum, L., McGinnis, C.S., McKinley, K.L., Rathnayake, A., Vaka, D., Nguyen, V., et al. (2024). Epithelial zonation along the mouse and human small intestine defines five discrete metabolic domains. Nat Cell Biol 26, 250–262. 10.1038/s41556-023-01337-z.

35. Hung, Y.H., Huang, S., Dame, M.K., Yu, Q.H., Yu, Q.C., Zeng, Y.A., Camp, J.G., Spence, J.R., and Sethupathy, P. (2021). Chromatin regulatory dynamics of early human small intestinal development using a directed differentiation model. Nucleic Acids Research 49. 10.1093/nar/gkaa1204.

36. Yu, Q., Kilik, U., Holloway, E.M., Tsai, Y.H., Harmel, C., Wu, A., Wu, J.H., Czerwinski, M., Childs, C.J., He, Z., et al. (2021). Charting human development using a multi-endodermal organ atlas and organoid models. Cell 184, 3281–3298 e3222. 10.1016/j.cell.2021.04.028.

37. Powell, D.W., Pinchuk, I.V., Saada, J.I., Chen, X., and Mifflin, R.C. (2011). Mesenchymal cells of the intestinal lamina propria. Annu Rev Physiol 73, 213–237. 10.1146/annurev.physiol.70.113006.100646.

38. Paerregaard, S.I., Wulff, L., Schussek, S., Niss, K., Morbe, U., Jendholm, J., Wendland, K., Andrusaite, A.T., Brulois, K.F., Nibbs, R.J.B., et al. (2023). The small and large intestine contain related mesenchymal subsets that derive from embryonic precursors. Nature Communications 14. ARTN 230710.1038/s41467-023-37952-5.

39. Han, L., Chaturvedi, P., Kishimoto, K., Koike, H., Nasr, T., Iwasawa, K., Giesbrecht, K., Witcher, P.C., Eicher, A., Haines, L., et al. (2020). Single cell transcriptomics identifies a signaling network coordinating endoderm and mesoderm diversification during foregut organogenesis. Nat Commun 11, 4158. 10.1038/s41467-020-17968-x.

40. Bahar Halpern, K., Massalha, H., Zwick, R.K., Moor, A.E., Castillo-Azofeifa, D., Rozenberg, M., Farack, L., Egozi, A., Miller, D.R., Averbukh, I., et al. (2020). Lgr5+telocytes are a signaling source at the intestinal villus tip. Nature Communications 11. ARTN 193610.1038/s41467-020-15714-x.

41. Cheng, H., and Leblond, C.P. (1974). Origin, differentiation and renewal of the four main epithelial cell types in the mouse small intestine. I. Columnar cell. Am J Anat 141, 461–479. 10.1002/aja.1001410403.

42. Cheng, H. (1974). Origin, differentiation and renewal of the four main epithelial cell types in the mouse small intestine. II. Mucous cells. Am J Anat 141, 481–501. 10.1002/aja.1001410404.

43. Cheng, H., and Leblond, C.P. (1974). Origin, differentiation and renewal of the four main epithelial cell types in the mouse small intestine. III. Entero-endocrine cells. Am J Anat 141, 503–519. 10.1002/aja.1001410405.

44. Cheng, H. (1974). Origin, differentiation and renewal of the four main epithelial cell types in the mouse small intestine. IV. Paneth cells. Am J Anat 141, 521–535. 10.1002/aja.1001410406.

45. Cheng, H., and Leblond, C.P. (1974). Origin, differentiation and renewal of the four main epithelial cell types in the mouse small intestine. V. Unitarian Theory of the origin of the four epithelial cell types. Am J Anat 141, 537–561. 10.1002/aja.1001410407.

46. Barker, N., van Es, J.H., Kuipers, J., Kujala, P., van den Born, M., Cozijnsen, M., Haegebarth, A., Korving, J., Begthel, H., Peters, P.J., and Clevers, H. (2007). Identification of stem cells in small intestine and colon by marker gene Lgr5. Nature 449, 1003–1007. 10.1038/nature06196.

47. Furuya, S., and Furuya, K. (2007). Subepithelial fibroblasts in intestinal villi: roles in intercellular communication. International review of cytology 264, 165–223. 10.1016/S0074-7696(07)64004-2.

48. Kaestner, K.H. (2019). The Intestinal Stem Cell Niche: A Central Role for Foxl1-Expressing Subepithelial Telocytes. Cell Mol Gastroenterol Hepatol 8, 111–117. 10.1016/j.jcmgh.2019.04.001.

49. Johnson, K.F., Dong, X., Tsai, Y.H., Wu, A., Clark, S.G., Huang, S., Zwick, R.K., Glass, I., Walton, K.D., Klein, O.D., and Spence, J.R. (2025). Mapping mesenchymal diversity in the developing human intestine and organoids. bioRxiv. 10.1101/2025.07.22.665939.

50. Aoki, R., Shoshkes-Carmel, M., Gao, N., Shin, S., May, C.L., Golson, M.L., Zahm, A.M., Ray, M., Wiser, C.L., Wright, C.V., and Kaestner, K.H. (2016). Foxl1-expressing mesenchymal cells constitute the intestinal stem cell niche. Cell Mol Gastroenterol Hepatol 2, 175–188. 10.1016/j.jcmgh.2015.12.004.

51. Tallini, Y.N., Greene, K.S., Craven, M., Spealman, A., Breitbach, M., Smith, J., Fisher, P.J., Steffey, M., Hesse, M., Doran, R.M., et al. (2009). c-kit expression identifies cardiovascular precursors in the neonatal heart. Proc Natl Acad Sci U S A 106, 1808–1813. 10.1073/pnas.0808920106.

52. Gannoun, L., De Schrevel, C., Belle, M., Dauguet, N., Achouri, Y., Loriot, A., Vanderaa, C., Cordi, S., Dili, A., Heremans, Y., et al. (2023). Axon guidance genes control hepatic artery development. Development 150. 10.1242/dev.201642.

53. Zhang, J., Song, Y., Wang, X., Wang, X., Li, S., Song, X., Zhao, C., Qi, J., Tian, Y., Zhao, B., et al. (2025). The transcription factor PITX1 cooperates with super-enhancers to regulate the expression of DUSP4 and inhibit pyroptosis in pulmonary artery smooth muscle cells. Respir Res 26, 149. 10.1186/s12931-025-03222-9.

54. Andras Czigler, L.T., Nikolett Szarka, Krisztina Szilágyi, Zoltan Kellermayer, Alexandra Harci, Monika Vecsernyes, Zoltan Ungvari, Alex Szolics, Akos Koller, Andras Buki, Peter Toth (2020). Prostaglandin E2, a postulated mediator of neurovascular coupling, at low concentrations dilates whereas at higher concentrations constricts human cerebral parenchymal arterioles. Prostaglandins & Other Lipid Mediators 146. 10.1016/j.prostaglandins.2019.106389.

55. Martin-Ventura, J.L., Madrigal-Matute, J., Munoz-Garcia, B., Blanco-Colio, L.M., Van Oostrom, M., Zalba, G., Fortuno, A., Gomez-Guerrero, C., Ortega, L., Ortiz, A., et al. (2009). Increased CD74 expression in human atherosclerotic plaques: contribution to inflammatory responses in vascular cells. Cardiovascular research 83, 586–594. 10.1093/cvr/cvp141.

56. Song, Y., Miao, Z., Brazma, A., and Papatheodorou, I. (2023). Benchmarking strategies for cross-species integration of single-cell RNA sequencing data. Nat Commun 14, 6495. 10.1038/s41467-023-41855-w.

57. McCarthy, N., Tie, G., Madha, S., He, R., Kraiczy, J., Maglieri, A., and Shivdasani, R.A. (2023). Smooth muscle contributes to the development and function of a layered intestinal stem cell niche. Developmental cell 58, 550–564 e556. 10.1016/j.devcel.2023.02.012.

58. Chinen, H., Matsuoka, K., Sato, T., Kamada, N., Okamoto, S., Hisamatsu, T., Kobayashi, T., Hasegawa, H., Sugita, A., Kinjo, F., et al. (2007). Lamina propria c-kit+ immune precursors reside in human adult intestine and differentiate into natural killer cells. Gastroenterology 133, 559–573. 10.1053/j.gastro.2007.05.017.

59. Romert, P., and Mikkelsen, H.B. (1998). c-kit immunoreactive interstitial cells of Cajal in the human small and large intestine. Histochemistry and cell biology 109, 195–202. 10.1007/s004180050218.

60. Torihashi, S., Ward, S.M., and Sanders, K.M. (1997). Development of c-Kit-positive cells and the onset of electrical rhythmicity in murine small intestine. Gastroenterology 112, 144–155. 10.1016/s0016-5085(97)70229-4.

61. Hao, Y., Hao, S., Andersen-Nissen, E., Mauck, W.M., 3rd, Zheng, S., Butler, A., Lee, M.J., Wilk, A.J., Darby, C., Zager, M., et al. (2021). Integrated analysis of multimodal single-cell data. Cell 184, 3573–3587 e3529. 10.1016/j.cell.2021.04.048.

62. Hafemeister, C., and Satija, R. (2019). Normalization and variance stabilization of single-cell RNA-seq data using regularized negative binomial regression. Genome Biol 20, 296. 10.1186/s13059-019-1874-1.

63. Becht, E., McInnes, L., Healy, J., Dutertre, C.A., Kwok, I.W.H., Ng, L.G., Ginhoux, F., and Newell, E.W. (2018). Dimensionality reduction for visualizing single-cell data using UMAP. Nat Biotechnol. 10.1038/nbt.4314.

64. Blondel, V.D., Guillaume, J.L., Hendrickx, J.M., de Kerchove, C., and Lambiotte, R. (2008). Local leaders in random networks. Phys Rev E Stat Nonlin Soft Matter Phys 77, 036114. 10.1103/PhysRevE.77.036114.

65. Traag, V.A., Waltman, L., and van Eck, N.J. (2019). From Louvain to Leiden: guaranteeing well-connected communities. Sci Rep 9, 5233. 10.1038/s41598-019-41695-z.

66. Choudhary, S., and Satija, R. (2022). Comparison and evaluation of statistical error models for scRNA-seq. Genome Biol 23, 27. 10.1186/s13059-021-02584-9.

67. Haghverdi, L., Lun, A.T.L., Morgan, M.D., and Marioni, J.C. (2018). Batch effects in single-cell RNA-sequencing data are corrected by matching mutual nearest neighbors. Nat Biotechnol 36, 421–427. 10.1038/nbt.4091.

68. Baldarelli, R.M., Smith, C.L., Ringwald, M., Richardson, J.E., Bult, C.J., and Mouse Genome Informatics, G. (2024). Mouse Genome Informatics: an integrated knowledgebase system for the laboratory mouse. Genetics 227. 10.1093/genetics/iyae031.

69. Korsunsky, I., Millard, N., Fan, J., Slowikowski, K., Zhang, F., Wei, K., Baglaenko, Y., Brenner, M., Loh, P.R., and Raychaudhuri, S. (2019). Fast, sensitive and accurate integration of single-cell data with Harmony. Nat Methods 16, 1289–1296. 10.1038/s41592-019-0619-0.

