## Supplementary figures and images for "Human-mouse cross-species comparison identifies common and unique aspects of intestinal mesenchyme development"

### Supplemental Figure 1

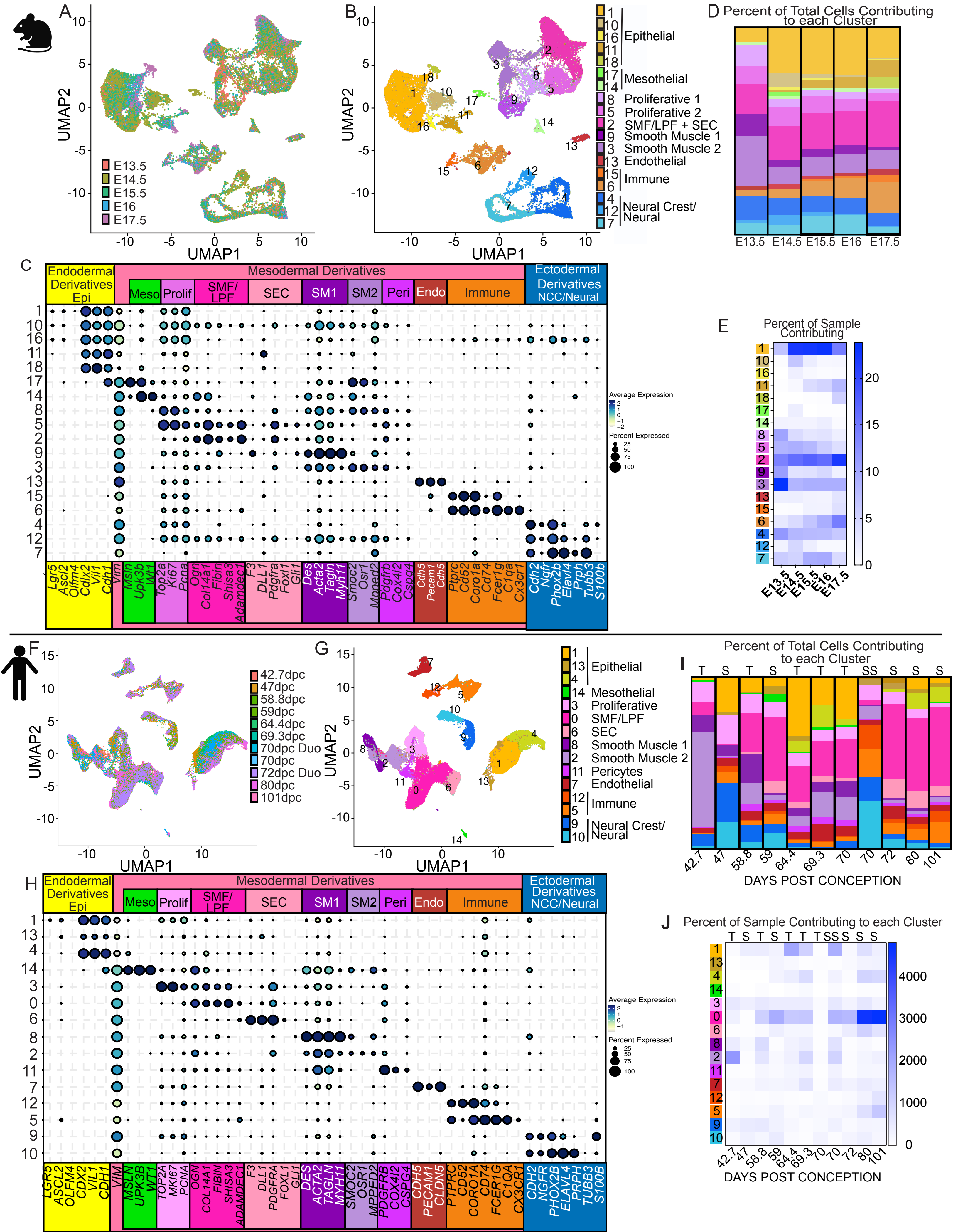

### Supplemental Figure 2

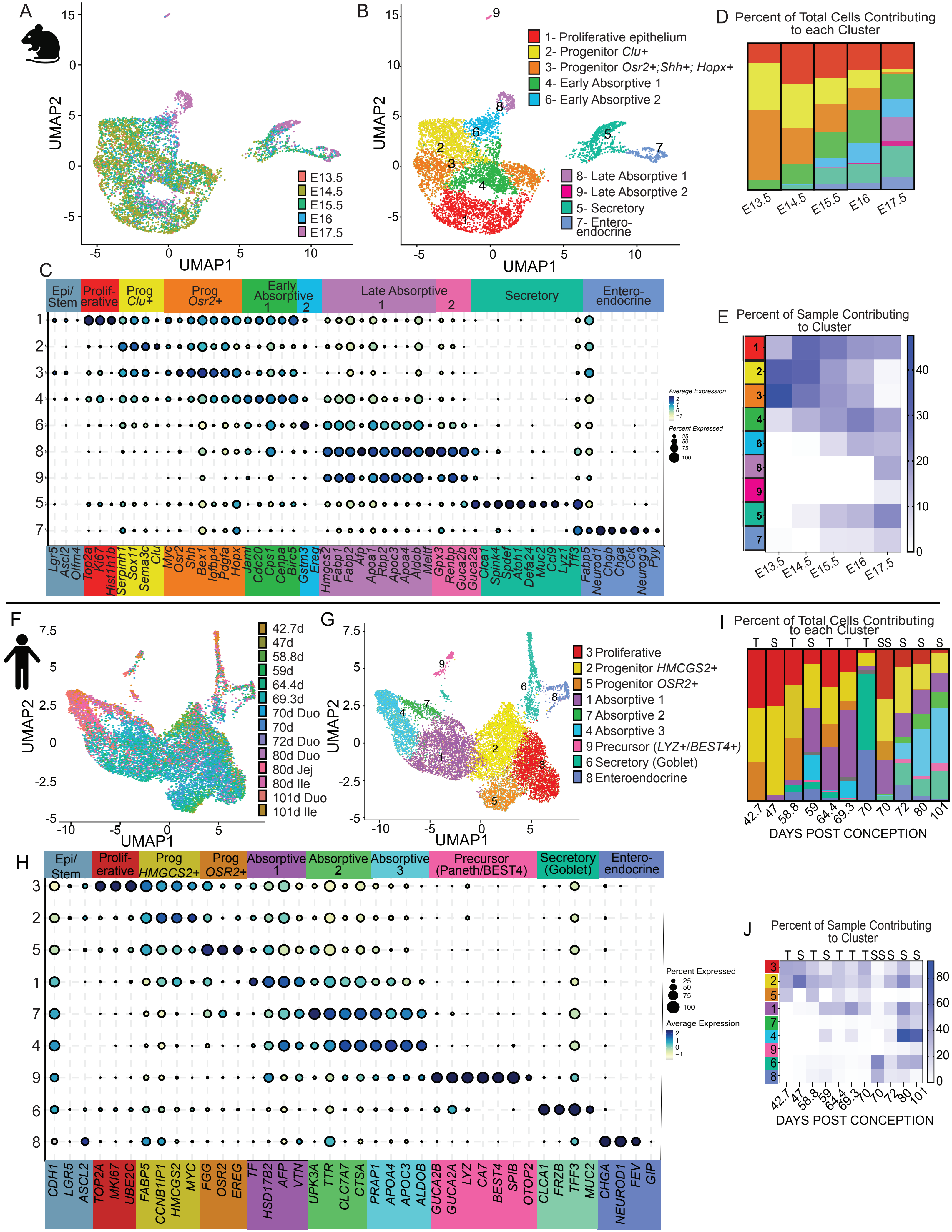

### Supplemental Figure 3

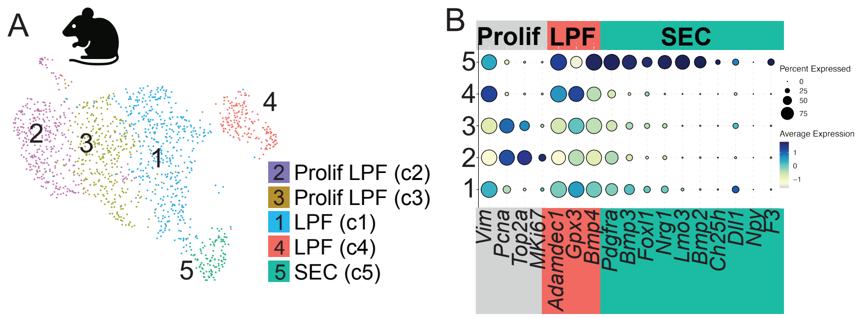

### Supplemental Figure 4

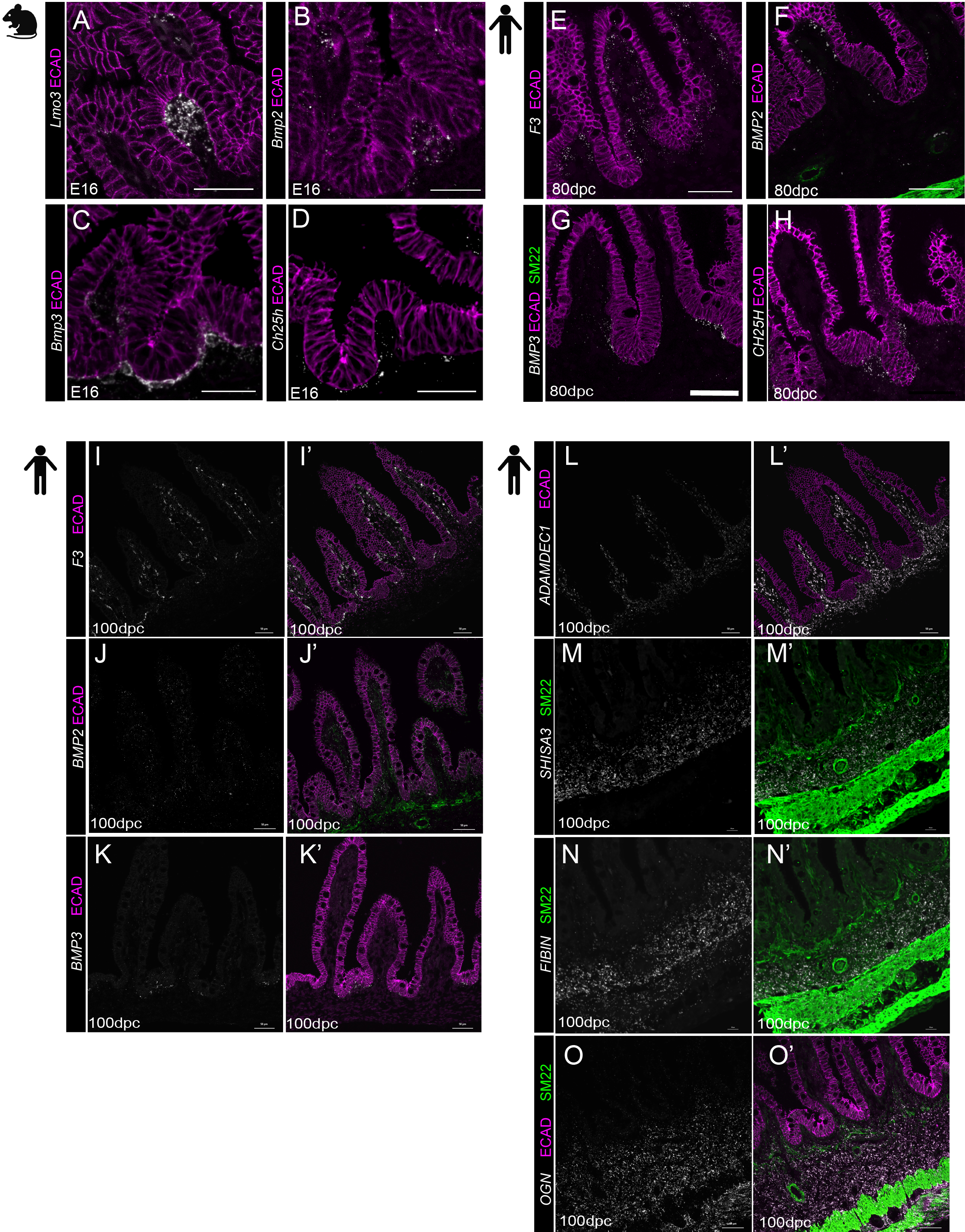
