## Supplemental Table 1 for "Human-mouse cross-species comparison identifies common and unique aspects of intestinal mesenchyme development"

| | | | | |
|---|---|---|---|---|
| Gm1992 | Gm28644 | Gm28634 | Aox4 | Apol7d |
| Gm37381 | 4931428L18Rik | Gm29260 | Gm15759 | Gm8883 |
| Gm37323 | Gm29128 | Gm9915 | Gm15834 | Gm28818 |
| Gm37988 | Pih1d3 | Gm29664 | Fam126b | 4933417E11Rik |
| Gm16041 | Gm5415 | Gm28175 | Als2cr12 | 4930556G22Rik |
| 4732440D04Rik | Gm37724 | Gm28140 | C2cd6b | Gm4319 |
| Gm26901 | Gm37591 | Gm28782 | Gm29018 | D230017M19Rik |
| Gm30414 | Gm37958 | 8430432A02Rik | G730003C15Rik | 4-Mar |
| 3110035E14Rik | Gm37233 | Gm29040 | Gm29017 | Gm39662 |
| Gm29520 | Gm28306 | 1500015O10Rik | Gm29016 | 1700027A15Rik |
| 1700034P13Rik | 4930568A12Rik | Gm29155 | Gm26813 | Gm29185 |
| Gm15818 | Prss39 | Gm29157 | Gm28411 | Pinc |
| Gm17644 | 1700101I19Rik | Gm29156 | 1700122D07Rik | D530049I02Rik |
| Gm29663 | Gm38336 | Gm8251 | Gm11579 | C530043A13Rik |
| Gm29283 | Gm37068 | Kdelc1 | 2310016D23Rik | Gm29186 |
| Gm29570 | Gm28415 | 4930521E06Rik | Gm28083 | 6030407O03Rik |
| Gm9947 | 1110002O04Rik | Gm5269 | Gm11587 | Gm29183 |
| Gm28095 | Gm37146 | Gm28151 | 9530026F06Rik | Gm28364 |
| Gm7568 | Gm33222 | Gm28826 | Gm11588 | Gm29539 |
| D030040B21Rik | 4930535G08Rik | C230029F24Rik | Gm28449 | Gm15841 |
| Gm28376 | Gm37335 | 4933411E06Rik | 4930587A21Rik | A630095N17Rik |
| Gm28783 | Gm33280 | Gm29665 | Pard3bos1 | Gm28294 |
| Gm28784 | 4930403P22Rik | Gm28321 | Pard3bos2 | Gm28902 |
| Gm28154 | Gm38033 | Gm28322 | Pard3bos3 | Gm15179 |
| Gm16070 | Gm42417 | Gm28319 | Gm29084 | Gm15178 |
| Gm28153 | Gm33533 | Gm29325 | Gm29083 | Gm29065 |
| Gm15825 | Gm37020 | 9330175M20Rik | Gm4208 | Gm29069 |
| Gm28340 | 4930439A04Rik | 1700072G22Rik | Gm20342 | Gm17751 |
| 4930486I03Rik | 2010300C02Rik | Gm17767 | Gpr1 | Gm816 |
| Gm28653 | 4930556I23Rik | Gm28055 | Gm11608 | Gm28386 |
| Gm28065 | Gm26805 | Gm31812 | Gm39653 | Gm28387 |
| 6720483E21Rik | Gm15457 | Gm553 | 4933402D24Rik | Gm28410 |
| Gm28287 | Gm5099 | Gm28178 | Gm13749 | BC035947 |
| Gm28836 | Gm16150 | 1700019A02Rik | Gm26649 | 5730419F03Rik |
| Gm26580 | Gm16151 | Gm28777 | Gm13748 | Gm29611 |
| 4933415F23Rik | Gm16152 | Gm28778 | Gm28981 | Gm29536 |
| Gm27028 | Gm37707 | Gm28551 | Gm28982 | Gm29187 |
| Gm29107 | Gm37821 | 4930444A19Rik | Gm29152 | 9830004L10Rik |
| Gm28822 | D930019O06Rik | Gm10561 | Gm28845 | 2310015K22Rik |
| Gm28237 | Gm3646 | 9130227L01Rik | Gm10558 | Gm29125 |
| Gm29414 | 1700066B17Rik | 1700003I22Rik | Gm15668 | 1700016L21Rik |
| 4931408C20Rik | Gm16894 | Gm28240 | Gm15669 | Gm45261 |
| Gm5524 | Gm35801 | 1700126A01Rik | Gm15671 | Gm9747 |
| Gm597 | Gm37623 | BC055402 | Gm28497 | Gm28940 |
| Gm9898 | Gm37915 | 4930558J18Rik | Gm29113 | Gm47791 |
| Gm29669 | 4930448I06Rik | Gm17234 | Gm29114 | Gm28942 |

| | | | | |
|---|---|---|---|---|
| Gm7544 | Gm28722 | Gm15699 | 2900009J06Rik | Gm37552 |
| Gm47955 | Gm9991 | Gm29012 | Gm28800 | Gm26781 |
| Gm47959 | Gm28199 | Gm17634 | Dars | Gm37799 |
| Gm47969 | Gm26683 | 2310035C23Rik | Gm26686 | Gm33994 |
| 4933436I20Rik | Gm17090 | Gm7160 | Gm16081 | 4933409D19Rik |
| C130026I21Rik | 1700020N18Rik | Gm20753 | Gm15674 | Platr23 |
| A530032D15Rik | Gm28382 | A530053M12Rik | Gm15675 | Gm19705 |
| Gm10553 | Gm28380 | D630008O14Rik | Gm16083 | Platr22 |
| A530040E14Rik | Gm29100 | Gm15391 | Gm29427 | Gm26979 |
| Gm16028 | Gm26720 | Gm15389 | Gm28857 | Gm26936 |
| Gm16025 | Gm29099 | D830032E09Rik | Gm28856 | A430106G13Rik |
| Gm16092 | Olfr1416 | Gm29088 | Gm15848 | Gm28556 |
| G530012D18Rik | Olfr1415 | Gm28189 | Gm28914 | Gm28501 |
| Gm16094 | Olfr1414 | 9330185C12Rik | Gm28913 | Gm5833 |
| Gm10552 | Olfr1413 | Gm19965 | Gm26892 | 1700019P21Rik |
| Gm17017 | 1700054K02Rik | Zfp813-ps | Gm29629 | 4933436E23Rik |
| A630001G21Rik | Olfr1412 | Gm28363 | 1700037F24Rik | Gm34816 |
| 9930111H07Rik | Olfr1411 | B020011L13Rik | Gm10188 | Gm4788 |
| 4933407L21Rik | Olfr1410 | Gm28360 | Gm29630 | 4930590L20Rik |
| Gm28884 | Olfr12 | Gm28168 | F730311O21Rik | Trove2 |
| Gm28100 | Gm29483 | Gm7145 | Gm15849 | Gm29514 |
| Gm21972 | Gm29482 | Gm29106 | Gm10538 | Gm15584 |
| Gm16341 | Gm29481 | Gm28867 | Gm28609 | Gm29515 |
| Gm28626 | Gm29480 | 2900060B14Rik | Gm19461 | Gm29020 |
| Gm28375 | 5033417F24Rik | Gm26831 | Gm28040 | Gm29398 |
| Gm29371 | 9430060I03Rik | Gm29456 | Kiss1.1 | Ptgs2os2 |
| Gm29374 | Gm28086 | Gm29455 | Gm26706 | 2310030A07Rik |
| Efhd1os | 2310007B03Rik | Gm27184 | Zc3h11a.1 | BC003331 |
| 3110079O15Rik | Gm28535 | 3830432H09Rik | Gm38394 | Gm20631 |
| Gm19582 | 2-Sep | Gm28209 | Gm28441 | Gm29188 |
| Ugt1a10 | Sept2.1 | 1700012E03Rik | Gm15851 | C730036E19Rik |
| 4930453O03Rik | Gm28536 | 2610027F03Rik | Gm1627 | Gm47985 |
| Gm29538 | Gm10550 | Gm29346 | Chil1 | Gm8947 |
| Gm28888 | 4930440C22Rik | Gm29345 | 4933406M09Rik | Gm47995 |
| Glrp1 | 4930598F16Rik | Gm29348 | Platr1 | 3110040M04Rik |
| Gm19589 | 1700063A18Rik | Gm29347 | Gm26783 | Gm47996 |
| Gm29336 | Gm29601 | Celrr | Gm28892 | Gm10138 |
| Platr5 | 4930533P14Rik | Gm29359 | Gm10535 | Gm28181 |
| Gm29337 | Panct2 | 2210011K15Rik | Gm15445 | Fam129a |
| 1700067G17Rik | Gm28901 | Gm15392 | Gm26642 | 2810414N06Rik |
| Gm28342 | D1Ertd622e | Htr5b | Gm4793 | Gm28610 |
| C030007H22Rik | 1810006J02Rik | Gm28590 | Gm38399 | Gm28960 |
| Iqca | Gm29461 | Gm28928 | Gm37759 | Gm29529 |
| 4930434B07Rik | Gm7967 | 1700121L03Rik | 5730559C18Rik | C230024C17Rik |
| Gm28721 | Gm20268 | Gm28706 | Gm26568 | Gm28513 |
| Gm28723 | Gm28187 | 4930599A14Rik | A430034D21Rik | Gm29290 |

| | | | | |
|---|---|---|---|---|
| Gm29291 | 1700029M03Rik | Ifi214 | 1700047M11Rik | Gm29678 |
| Gm28286 | 4930568G15Rik | Ifi213 | Gm37885 | Gm30725 |
| A830008E24Rik | Gm38325 | Ifi209 | Gm37218 | Gm30932 |
| 4930452N14Rik | Gm37073 | Ifi208 | Gm38079 | Gm31373 |
| 4930532M18Rik | Gm36972 | Ifi203 | Gm37986 | A230020J21Rik |
| Gm29282 | Gm37469 | Olfr433 | Gm33973 | Gm38188 |
| Gm9530 | Gm26665 | Olfr432 | Gm34068 | Gm38161 |
| Gm37622 | Gm37923 | Olfr430 | Gm38108 | Fam71a |
| Gm5532 | Dusp27 | Olfr429 | 1700056E22Rik | D730003I15Rik |
| Gm37571 | 5330438I03Rik | Olfr427 | Gm37223 | Gm37168 |
| 4930518J20Rik | Gm15853 | Olfr231 | Gm37712 | Tmem206 |
| Gm2000 | Gm4846 | Olfr424 | 1700112H15Rik | Gm32200 |
| Gm10031 | Gm4847 | Olfr420 | Gm37800 | Gm37432 |
| 4930439D14Rik | Fmo9 | Olfr419 | 4930488B22Rik | Gm38037 |
| Gm28694 | Gm16701 | Olfr417 | Gm34342 | 1700034H15Rik |
| Gm15486 | Gm16701.1 | Olfr248 | 1-Mar | Gm26670 |
| Gm37679 | Gm37982 | Olfr414 | 2-Mar | Gm26879 |
| 1600012P17Rik | Gm2453 | Olfr220 | Gm38251 | Gm10516 |
| 4930562F17Rik | Gm20711 | Gm36307 | Gm34732 | 1700065J18Rik |
| Gm38039 | Gm37839 | Gm38100 | 4930433J02Rik | Gm15867 |
| Gm37294 | 3110045C21Rik | Rbm8a2 | Gm37662 | Gm31406 |
| 4930562F07Rik | Gm37748 | Gm38101 | Gm34882 | Diexf |
| Gm36975 | Gm37502 | Gm36536 | 2010103J01Rik | Gm15872 |
| Gm37083 | 4930500M09Rik | Gm36992 | 4930532G15Rik | 4930570N18Rik |
| Gm38060 | Gm26620 | Gm15423 | Eprs | Gm37650 |
| Gm37362 | Gm38204 | Gm26801 | 9630028B13Rik | 4631405K08Rik |
| Gm37634 | Gm10522 | Gm37306 | Gm2061 | 4930503O07Rik |
| Gm31925 | Gm20045 | Gm37280 | Gm38093 | Gm16897 |
| Gm38304 | Gm26641 | 2310043L19Rik | Gm37134 | Gm32250 |
| 2810442N19Rik | B930036N10Rik | 1700016C15Rik | A730004F24Rik | Gm37027 |
| Dnm3os | Cd244 | Adss | Gm29491 | Gm13184 |
| Gm37273 | Gm37787 | B230369F24Rik | A430105J06Rik | Gm45902 |
| Gm20471 | Gm10521 | Gm16564 | 1700023G09Rik | Gm13185 |
| Gm26523 | Gm37065 | Gm10518 | Gm37912 | Gm13180 |
| Gm29500 | Gm37756 | Gm37336 | D1Pas1 | Gm13191 |
| Mettl11b | Gm17224 | Gm31728 | 1700007P06Rik | 1700080N15Rik |
| BC055324 | Gm37950 | Gm37267 | Gm38155 | Gm13189 |
| Gm16587 | Gm36937 | Gm37768 | Gm37896 | Gm13175 |
| Gm16548 | Olfr16 | Gm38293 | Gm36388 | Gm13179 |
| Gm32391 | Olfr1408 | Gm17275 | 9330162B11Rik | 4930551O13Rik |
| Gm26685 | Olfr1406 | H3f3aos | Gm38064 | Gm26776 |
| Gm37754 | Olfr218 | 2210411M09Rik | A430027H14Rik | Gm13199 |
| Gm37853 | Olfr1404 | 9130409I23Rik | Gm26574 | Gm13267 |
| Gm37184 | Olfr418 | Gm37406 | Gm37422 | Gm10857 |
| Gm36945 | Mptx2 | A430110L20Rik | Gm28172 | A230108P19Rik |
| Gm32569 | Ifi206 | Gm5533 | Prox1os | Gm13383 |

| | | | | |
|---|---|---|---|---|
| Gm13388 | 4931423N10Rik | Gm13547 | Olfr339 | Gm28639 |
| Gm13391 | Il1f8 | 2600006K01Rik | Olfr340 | Gm13497 |
| Gm13389 | Il1f9 | Wdr34 | Olfr341 | Gm13499 |
| Gm26565 | Il1f6 | 1700084E18Rik | Olfr342 | 1700057H21Rik |
| Gm28641 | Il1f5 | Gm28035 | Olfr344 | Tas2r134 |
| Gm13211 | Gm13411 | Gm28038 | Olfr345 | Gm13490 |
| 1700061F12Rik | Gm13417 | Gm16323 | Olfr346 | Gm13522 |
| Gm13219 | Gm13387 | Gm14487 | Olfr347 | Gm26603 |
| Gm13218 | Gm38287 | Cstad | Olfr348 | Gm14033 |
| Gm13256 | Fam166a | 9330198N18Rik | Olfr50 | Gm26794 |
| 4930412O13Rik | AL732309.1 | Gm14486 | Olfr3 | Gm33594 |
| 9230102O04Rik | AL732309.2 | 4930527E20Rik | Olfr350 | Gm13546 |
| Gm13262 | Grin1os | Gm13405 | Olfr351 | Gm13544 |
| Atp5c1 | AA543186 | Gm13425 | Olfr352 | Gm17409 |
| Gm13264 | Gm996 | Gm13427 | Olfr353 | Gm11099 |
| Gm38171 | Gm26702 | Gm13441 | Olfr354 | Gm13572 |
| 8030442B05Rik | Fcnaos | Gm13442 | Olfr355 | Gm13620 |
| Gm13293 | Lcn5 | Gm13609 | Olfr356 | 7-Mar |
| Gm13291 | A230005M16Rik | Gm13610 | Olfr357 | Gm13571 |
| Gm10851 | Bmyc | 1110008P14Rik | Olfr358 | Gm13580 |
| Gm17490 | Lcn3 | Gm13412 | Olfr360 | Gm13583 |
| Fbxo18 | Lcn11 | Fam102a | Olfr361 | Gm13582 |
| E030013I19Rik | Gm13539 | 9430097D07Rik | Olfr362 | Gm13561 |
| Gm37565 | Gm13554 | 6330409D20Rik | Olfr364-ps1 | Gm13565 |
| Gm37126 | Tmem250-ps | Gm13524 | Olfr365 | Gm13564 |
| Gm10848 | 1810012K08Rik | Gm13523 | Olfr366 | Gm13569 |
| Gm13269 | Gm13553 | Fam129b | Olfr368 | Gm13575 |
| Gm13322 | Gm27196 | Gm13528 | Gm13556 | Gm13594 |
| Gm13324 | Gm13563 | Gm13536 | Gm27197 | Gm13595 |
| A930004D18Rik | Sdccag3 | Gm13530 | Gm13584 | Gm13619 |
| Gm13330 | Gm20532 | C130021I20Rik | Gm13589 | Gm13617 |
| Carlr | Fam69b | 9430024E24Rik | Nr5a1os | Gm28638 |
| 4930426L09Rik | Snhg7os | Gm13403 | Nr6a1os | Gm13618 |
| 4930447M23Rik | Gm13398 | Gm13404 | Gm13496 | Gm13629 |
| Gm33940 | Dbhos | C230014O12Rik | Arhgap15os | Gm13598 |
| Gm13344 | Sardhos | Gm38389 | Gm28643 | Gm1322 |
| Ptf1aos | AA645442 | C79798 | Gm13479 | Gm13599 |
| Gm13348 | Gm13371 | Gm34653 | 1700019E08Rik | Gm21830 |
| Gm13346 | F730016J06Rik | Gm13449 | Gm13465 | Gm26727 |
| Gm17171 | Gm13373 | Gm13605 | Gm13467 | Gm13601 |
| Gm13335 | 1700007K13Rik | 4930568D16Rik | Gm13470 | Gm26743 |
| Gm13362 | Gm10134 | 4930402F06Rik | Gm13481 | Gm13596 |
| Gm13375 | Gm13381 | Gm35202 | Gm13480 | AL844841.2 |
| Gm13376 | Gm13393 | 9030204H09Rik | Gm13483 | 4933409G03Rik |
| Gm13327 | Gm13402 | Gm13431 | Gm48908 | 4932414N04Rik |
| 1700092C17Rik | Gm13420 | Olfr338 | Gm13485 | Gm36231 |

| | | | | |
|---|---|---|---|---|
| Gm13613 | Ttc30b | Olfr1015 | Olfr1082 | Olfr1134 |
| Ccdc173 | Ttc30a2 | Olfr1016 | Olfr1084 | Olfr1135 |
| Mettl5os | Ttc30a1 | Olfr1018 | Olfr1085 | Olfr1136 |
| Gad1os | Gm13938 | Olfr1019 | Olfr1086 | Olfr1137 |
| Erich2os | Gm13943 | Olfr1020 | Olfr1087 | Olfr1138 |
| Gm26558 | Gm13944 | Olfr1022 | Olfr1089 | Olfr1140 |
| Gm13632 | Gm13661 | Olfr1023 | Olfr1090 | Olfr1141 |
| Dlx1as | 4930440I19Rik | Olfr1024 | Olfr1093 | Olfr152 |
| Platr26 | Gm14461 | Olfr1025-ps1 | Olfr141 | Olfr1143 |
| Gm13647 | Gm26755 | Olfr1026 | Olfr1094 | Olfr1145 |
| Gm17250 | Ssfa2 | Olfr1028 | Olfr1095 | Olfr1148 |
| Gm13746 | Gm13752 | Olfr1029 | Gm13723 | Olfr1150-ps1 |
| Rapgef4os1 | A130030D18Rik | Olfr1030 | Olfr1097 | Olfr1151 |
| Rapgef4os2 | Gm28675 | Olfr1031 | Olfr1098 | Olfr1152 |
| Rapgef4os3 | Gm13686 | Olfr1032 | Olfr1099 | Olfr1153 |
| 8430437L04Rik | Gm13698 | Olfr1033 | Olfr1100 | Olfr1154 |
| 6430710C18Rik | Gm13693 | Olfr1034 | Olfr1101 | Olfr1155 |
| Gm11084 | Gm13695 | Olfr1036 | Olfr1102 | Olfr1156 |
| Sp3os | Gm13696 | Olfr1037 | Olfr1104 | Olfr1157 |
| Gm13703 | Gm13694 | Olfr1039 | Olfr1105 | Olfr74 |
| Gm13709 | Gm13697 | Olfr1040 | Olfr1106 | Olfr1158 |
| Gm13707 | Gm13691 | Olfr1042 | Olfr1107 | Olfr1160 |
| Gm13708 | Gm13710 | Olfr1043 | Olfr1109 | Olfr1161 |
| Chrna1os | Gm28635 | Olfr1044 | Olfr259 | Olfr73 |
| Chn1os3 | Gm19426 | Olfr52 | Olfr1110 | Olfr1162 |
| Chn1os1 | Gm13713 | Olfr1045 | Olfr1111 | Olfr1163 |
| Gm10822 | Gm13716 | Olfr1046 | Olfr1112 | Olfr1164 |
| Atp5g3 | Gm13715 | Olfr1047 | Olfr1113 | Olfr1166 |
| A630050E04Rik | Olfr987 | Olfr1048 | Olfr1115 | Olfr1167 |
| Gm28640 | Olfr988 | Olfr1049 | Olfr1116 | Olfr1168 |
| Gm13667 | Olfr992 | Olfr1051 | Olfr1118 | Olfr1170 |
| 4930441J16Rik | Olfr993 | Olfr1052 | Olfr1120 | Olfr1173 |
| 4930445N08Rik | Olfr994 | Olfr1053 | Olfr1121 | Olfr1174-ps |
| Gm14396 | Olfr995 | Olfr1054 | Olfr1122 | Olfr1175-ps |
| Gm28793 | 4833423E24Rik | Olfr1055 | Olfr1123 | Olfr1176 |
| Hoxd3os1 | Olfr996 | Olfr1056 | Olfr1124 | Olfr1177-ps |
| Hoxd4.1 | Olfr998 | Olfr1057 | Olfr1126 | Olfr1178 |
| Gm28230 | Olfr1000 | Olfr1058 | Pramel7 | Olfr1179 |
| Gm14424 | Olfr1002 | Olfr1061 | Pramel6 | Olfr1180 |
| Gm14426 | Olfr154 | Olfr1062 | Olfr153 | Olfr1181 |
| Gm13652 | Olfr1006 | Olfr1065 | Olfr1128 | Gm13757 |
| 2600014E21Rik | Olfr1008 | Olfr1066 | Olfr1129 | Olfr1183 |
| A330043C09Rik | Olfr1009 | Olfr228 | Olfr1130 | Olfr1184 |
| Gm13658 | Olfr1012 | Olfr1076 | Olfr1131 | Olfr1185-ps1 |
| Gm13660 | Olfr1013 | Olfr1079 | Olfr1132 | Olfr1186 |
| Gm13656 | Olfr1014 | Olfr1080 | Olfr1133 | Olfr1188 |

| | | | | |
|---|---|---|---|---|
| Olfr1189 | Olfr1245 | Gm10804 | Olfr1279 | Gm13976 |
| Olfr1191-ps1 | Olfr1246 | Gm10803 | Olfr1280 | 3110099E03Rik |
| Olfr1192-ps1 | Olfr1247 | Gm27027 | Olfr1281 | Gm13977 |
| Olfr1193 | Olfr1248 | Alkbh3os1 | Olfr1282 | BC052040 |
| Olfr1195 | Olfr1249 | Gm13793 | Olfr1283 | 2700033N17Rik |
| Olfr1196 | Olfr1250 | Gm13794 | Olfr1284 | G630016G05Rik |
| Olfr1197 | Olfr1251 | Gm13799 | Olfr1286 | Gm13990 |
| Olfr1198 | Olfr1252 | 4930445B16Rik | Olfr1287 | D330050G23Rik |
| Olfr1199 | Olfr1253 | Gm10801 | Olfr1288 | Gm29340 |
| Olfr1200 | Olfr1254 | Gm10800 | Olfr1289 | Gm13982 |
| Olfr1201 | Olfr1255 | Gm29235 | Olfr1290 | Gm28183 |
| Olfr1202 | Olfr1256 | B230118H07Rik | Olfr1291-ps1 | Gm13985 |
| Olfr1205 | Olfr48 | Gm13905 | Olfr1293-ps | 4930412B13Rik |
| Olfr1204 | Olfr1257 | Gm13919 | Olfr1294 | Gm29233 |
| Olfr1206 | Olfr1258 | Gm13920 | Olfr1295 | A930104D05Rik |
| Olfr1208 | Olfr1259 | Gm13872 | Olfr1297 | 1700054M17Rik |
| Olfr1209 | Olfr1260 | Gm13874 | Olfr1298 | A430105I19Rik |
| Olfr1211 | Olfr1261 | Gm13869 | Olfr1299 | Gm14091 |
| Olfr1212 | Olfr1262 | A930006I01Rik | Olfr1301 | Gm14089 |
| Gm13762 | Olfr1263 | BC016548 | Olfr1302 | 1700020I14Rik |
| Olfr1214 | Olfr1264 | Gm13881 | Olfr1303 | Gm13999 |
| Olfr1215 | Olfr1265 | 4931422A03Rik | Olfr1305 | Gm28042 |
| Olfr1216 | Olfr140 | Gm10912 | Olfr1306 | Gm26899 |
| Olfr1217 | Olfr1269 | Gm10799 | Olfr1307 | Gm14978 |
| Olfr1218 | Olfr32 | Gm13883 | Olfr1308 | Gm14019 |
| Olfr1219 | Olfr1270 | Gm11060 | Olfr1309 | Casc4 |
| Olfr1220 | Olfr1506 | 4930527A07Rik | Olfr1310 | Gm14085 |
| Olfr1221 | Olfr142 | Pax6os1 | Olfr1311 | Bloc1s6os |
| Olfr1222 | Olfr1271 | Gm13954 | Olfr1312 | 4930517E11Rik |
| Olfr1223 | Olfr1272 | Gm29053 | Olfr1313 | 4930583P06Rik |
| Olfr1224-ps1 | Gm13769 | Gm14015 | Olfr1314 | Gm13994 |
| Olfr1225 | Olfr1274-ps | Gm14014 | Olfr1315-ps1 | Gm14002 |
| Olfr1226 | 4933423P22Rik | Gm13912 | Olfr1316 | A530010F05Rik |
| Olfr1228 | A330069E16Rik | Gm13923 | Olfr1317 | Gm14003 |
| Olfr1229 | Gm17281 | Gm45346 | Olfr1318 | Gm9913 |
| Olfr1230 | 1110051M20Rik | Ccdc34os | Gm21985 | Gm17555 |
| Olfr1231 | Gm9821 | Gm13936 | Gm13963 | Gm26697 |
| Olfr1232 | Gm13780 | Gm13942 | Gm13966 | Gm27003 |
| Olfr1233 | 2900072N19Rik | Gm13941 | Gm13964 | 1810024B03Rik |
| Olfr1234 | 1700029I15Rik | Gm15130 | 4930533B01Rik | Gm14244 |
| Olfr1238 | Gm13814 | 4930430A15Rik | C130080G10Rik | 1500011K16Rik |
| Olfr1239 | Gm13817 | Olfr1275 | Gm29234 | Gm14005 |
| Olfr1240 | Gm13816 | Gm13962 | Gm13974 | Gm14009 |
| Olfr1241 | 4631405J19Rik | Olfr1276 | Gm28494 | Gm14012 |
| Olfr1242 | Gm13802 | Olfr1277 | Gm28493 | Gm14010 |
| Olfr1243 | Gm13807 | Olfr1278 | 4930528P14Rik | Gm29010 |

| | | | | |
|---|---|---|---|---|
| Spdye4c | Gm28214 | Gm14144 | 4930518I15Rik | Wfdc16 |
| Gm39929 | Gm14055 | 4930442J19Rik | 1110008F13Rik | Gm11457 |
| Gm14011 | Gm14064 | Gm14152 | Soga1 | Zfp335os |
| 4933427J07Rik | Gm14062 | Gm14146 | Gm14286 | 1700025C18Rik |
| Gm10762 | Gm14061 | 4921509C19Rik | Platr27 | Gm14397 |
| Gm14025 | Gm17374 | Gm14167 | 2010009K17Rik | Gm14437 |
| AI847159 | Macrod2os2 | 4930556L07Rik | 1700060C20Rik | Gm28163 |
| Gm14029 | Macrod2os1 | Gm14155 | Gm23925 | Gm11465 |
| A730036I17Rik | Kif16bos | Rspo4os | Gm14205 | Platr29 |
| Gm14024 | Pcsk2os1 | Gm14154 | Gm14223 | Gm11464 |
| Il1bos | Pcsk2os2 | AA387200 | 9130015L21Rik | Gm11466 |
| F830045P16Rik | Banf2os | Gm14161 | Gm826 | Gm11467 |
| Gm14041 | Gm5535 | Gm17416 | Gm14221 | Gm11468 |
| Gm28196 | 4930444E06Rik | Defb45 | Gm27206 | Gm14268 |
| Gm28372 | 1700108N11Rik | 1700030C14Rik | Gm16751 | 5031425F14Rik |
| 4930473A02Rik | Gm14092 | Gm26841 | Gm14228 | Gm17096 |
| 4930402H24Rik | A930019D19Rik | Gm14199 | Gm45447 | Gm14291 |
| Gm11037 | 4933406D12Rik | 4930404H24Rik | Gm14243 | Gm20431 |
| 1700037H04Rik | Gm14114 | Dnmt3bos | Gm14242 | Tmem189 |
| Gm14232 | 6430503K07Rik | Dnmt3c | Gm14241 | Gm14320 |
| Gm14285 | Gm14110 | Bpifa6 | Ptprtos | A530013C23Rik |
| Gm14280 | AI646519 | Bpifb9b | Gm14246 | Gm14321 |
| 5330413P13Rik | A530006G24Rik | 4930519P11Rik | Gm14255 | 9230111E07Rik |
| Erv3 | Gm14120 | Gm14198 | Gm11454 | 1200007C13Rik |
| Prn | 8030411F24Rik | Gm14216 | Gm14254 | Gm14319 |
| Gm14051 | 9230104L09Rik | Gm14214 | 2900093K20Rik | Gm14236 |
| AV099323 | Gm1330 | Gm14226 | 2310001K24Rik | Gm14235 |
| 4921508D12Rik | Gm10750 | Gm45609 | Hnf4aos | E130018N17Rik |
| Gm14095 | Gm14135 | 2310005A03Rik | 0610039K10Rik | Gm20716 |
| 1700026D11Rik | Cst10 | Acss2os | Gm16316 | Gm14262 |
| AU019990 | Gm14133 | Gssos2 | Wisp2 | 1700017J07Rik |
| 1110034G24Rik | C530025M09Rik | Gssos1 | Gm11455 | Gm14260 |
| Gm14097 | Zfp120 | Gm14257 | A730032A03Rik | Gm9873 |
| Gm14104 | Gm10770 | Gm17581 | Wfdc15a | Gm14258 |
| Gm14102 | Gm14139 | BC029722 | Wfdc15b | 9430093N23Rik |
| A430048G15Rik | Gm21994 | Gm16098 | Svs2 | Gm14250 |
| Gm14100 | Gm14124 | Gm15557 | Svs3b | 4930529I22Rik |
| 4930545L23Rik | 3300002I08Rik | Gm28036 | Svs4 | Gm14249 |
| 9630028H03Rik | Gm10130 | Gm14252 | Svs3a | Gm11011 |
| Platr3 | Platr15 | Gm14170 | Svs6 | Gm26883 |
| Gm14211 | Gm28450 | 2900097C17Rik | Svs5 | A630075F10Rik |
| Gm14210 | Gm26681 | Gm14225 | Gm14302 | Gm17619 |
| Pak7 | Gm14151 | Gm14224 | Gm20458 | Bcas1os2 |
| Gm14218 | Gm14147 | Gm14168 | Gm14317 | Bcas1os1 |
| AL731706.1 | Gm14145 | Gm14174 | Wfdc6a | Gm16796 |
| AL731706.2 | Gm14149 | Gm14230 | Wfdc6b | Gm14637 |

| | | | | |
|---|---|---|---|---|
| 4930470P17Rik | Gm26869 | Gm14505 | Btbd35f6 | 4930515L19Rik |
| Gm14264 | 4930591A17Rik | Drr1 | Gm21870 | Gm29242 |
| Gm14266 | Gm17180 | Cypt1 | Btbd35f9 | 9530027J09Rik |
| Gm14640 | Gata5os | 4930578C19Rik | Btbd35f8 | Gm595 |
| 1700007M16Rik | B230312C02Rik | Gm26652 | Gm14552 | Gm14696 |
| Gm14272 | Gm14340 | Tex13c3 | Gm2309 | Gm14697 |
| Gm14641 | Gm14342 | Zfp300 | Gm14553 | Olfr1320 |
| Gm14271 | Gm14344 | Gm21876 | Gm6268 | Olfr1321 |
| Gm14275 | Gm14339 | 4930453H23Rik | Mdrl | Olfr1322 |
| Gm14273 | Gm27032 | Gm6938 | Gm14549 | Olfr1323 |
| Gm14455 | Gm14341 | Gm26593 | C330007P06Rik | Olfr1324 |
| Gm14453 | Uckl1os | Gm28269 | Gm15008 | 1700080O16Rik |
| Ctcflos | Gm14496 | Gm28268 | 6-Sep | Gm14582 |
| Pmepa1os | Btbd35f23 | Gm4907 | Gm9 | A630012P03Rik |
| Gm14642 | Btbd35f24 | Gm4985 | Rhox1 | Hprt |
| 1700021F07Rik | Btbd35f11 | Gm27192 | Rhox2a | Gm28730 |
| 1700010B08Rik | Btbd35f18 | E330010L02Rik | Rhox4a | Fam122b |
| Gm10714 | Btbd35f10 | Gm2101 | Rhox4a2 | Fam122c |
| Nespas | Btbd35f16 | Gm2117 | Rhox2b | Gm14597 |
| Gm20721 | Btbd35f3 | Gm2165 | Rhox4b | 4930502E18Rik |
| Atp5e | Btbd35f28 | Gm2200 | Rhox2c | Zfp36l3 |
| Gm14618 | Btbd35f20 | Gm26818 | Rhox4c | Gm16405 |
| Gm14616 | Btbd35f17 | Gm3669 | Rhox2d | Gm16430 |
| Gm14393 | Btbd35f4 | E330016L19Rik | Rhox4d | Slxl1 |
| Gm14440 | Btbd35f27 | Gm7437 | Rhox2e | 1600025M17Rik |
| Zfp968 | Btbd35f7 | Gm14974 | Rhox4e | Gm2155 |
| Gm11007 | AU022751 | Btbd35f29 | Rhox2f | Gm2174 |
| Zfp969 | Flicr | Btbd35f26 | Rhox3f | Gm10477 |
| Gm2007 | Gm14703 | Btbd35f19 | Rhox4f | Gm648 |
| Gm14288 | Gm45208 | Gm21883 | Rhox3g | Gm14718 |
| Gm11008 | Gm10491 | Spin2e | Rhox3h | Gm364 |
| Gm2004 | Gm10490 | Btbd35f25 | Rhox2h | 4930550L24Rik |
| 2210418O10Rik | Gm14820 | Gm21637 | Rhox5 | Gm14661 |
| Gm11009 | Rbm3os | Btbd35f22 | Rhox6 | Gm14662 |
| Gm14435 | Gm14459 | Btbd35f2 | Rhox7a | Gm14664 |
| Gm14305 | Ssx9 | Btbd35f1 | Rhox8 | C230004F18Rik |
| Gm14295 | Gm6592 | Gm5926 | Rhox7b | Cdr1 |
| Gm14408 | Gm5751 | Btbd35f21 | Rhox9 | 4933402E13Rik |
| Gm14296 | B630019K06Rik | Btbd35f30 | Btg1-ps1 | 4931400O07Rik |
| Gm14419 | 4930402K13Rik | Gm21789 | Btg1-ps2 | Tslrn1 |
| Gm14410 | Gm14862 | Btbd35f5 | Gm7598 | Gm6760 |
| Gm14409 | Hypm | Spin2-ps6 | Gm14565 | 3830417A13Rik |
| Gm14406 | Gm10489 | Btbd35f12 | 6030498E09Rik | Ctag2 |
| Gm14327 | Gm14493 | Btbd35f15 | Cypt15 | 4930447F04Rik |
| Zfp972 | Gm14635 | Btbd35f13 | Cypt14 | 1700036O09Rik |
| Gm14326 | 2010308F09Rik | Btbd35f14 | Tex13c2 | 4933436I01Rik |

| | | | | |
|---|---|---|---|---|
| Fmr1os | Pet2 | 2610002M06Rik | 4932411N23Rik | Gm15104 |
| Gm14698 | 4932429P05Rik | Fam46d | Gm382 | Tmem29 |
| Gm6812 | 4930415L06Rik | Gm732 | 4921511C20Rik | Gm27191 |
| Gm14705 | Gm44 | Gm379 | 4930558G05Rik | Gm15138 |
| 1700111N16Rik | Gm14773 | Gm6377 | Gm26851 | A230072E10Rik |
| 1700020N15Rik | Gm5072 | Gm44593 | Gm15023 | Cypt3 |
| 1110012L19Rik | Gm8914 | Gm10112 | Pramel3 | Kctd12b |
| 4930567H17Rik | 1700084M14Rik | Gm45194 | Gm5128 | Gm45022 |
| Gm16189 | Gm14781 | 4933403O08Rik | Gm7903 | Samt1 |
| Gm1141 | Mageb1 | 2010106E10Rik | AV320801 | 4921511M17Rik |
| Gm14684 | Gm5941 | Gm14936 | Prame | Gm10057 |
| Gm14685 | 1700003E24Rik | Ube2dnl1 | Arxes2 | Gm15140 |
| DXBay18 | BC061195 | Ube2dnl2 | Arxes1 | 4930524N10Rik |
| Gm45015 | AU015836 | 4930555B12Rik | H2bfm | Samt4 |
| Gm18336 | Gm14798 | H2afb2 | Tmsb15l | Samt2 |
| Gm26726 | Gm14827 | H2afb1 | 4930513O06Rik | Gm15156 |
| Taz | Gm39526 | Gm28579 | 4933428M09Rik | Gm15155 |
| Gm6880 | Gm26617 | Gm14929 | Mum1l1 | Gm15169 |
| Olfr1326-ps1 | Pfn5 | H2afb3 | Trap1a | Gm15241 |
| Olfr1325 | 1700010D01Rik | Gm17521 | D330045A20Rik | Gm15243 |
| Gm5640 | F630028O10Rik | Astx6 | Gm15013 | Gm15262 |
| Gm6890 | Hsf3 | Srsx | Pih1h3b | Scml1 |
| Gm5936 | Pgr15l | Gm17577 | Gm15046 | Gm15205 |
| Gm15384 | Gm14812 | Gm14951 | Eif2c5 | Gm15202 |
| Gm15063 | Gm14809 | Astx2 | Gm15295 | Rnf138rt1 |
| Gm14715 | Gm14808 | Gm17412 | Gm15294 | Tmem27 |
| Gm14707 | Tmem28 | Gm14950 | Gm15298 | Gm17604 |
| Gm14717 | Gm14902 | Gm17467 | Kcne1l | Gm15226 |
| Gm14742 | Gm20489 | Cldn34c3 | A730046J19Rik | Gm1720 |
| Pbsn | Gm3880 | Astx5 | Gm15128 | Gm15230 |
| Gm14744 | Gm9112 | Vmn2r121 | Gm15080 | Gm8817 |
| 5430402E10Rik | Dmrtc1c1 | Astx1a | Gm15107 | Gm15232 |
| Obp1a | Dmrtc1c2 | Gm17584 | Gm15114 | Gm15228 |
| Gm5938 | 1700031F05Rik | Astx4a | Gm8334 | Gm15239 |
| Obp1b | 1700011M02Rik | Gm17469 | Gm15127 | Gm15261 |
| Gm14743 | 1700018G05Rik | Astx4b | Luzp4 | Gm15245 |
| 4930480E11Rik | Gm26952 | Astx1b | Gm15099 | 4933400A11Rik |
| Gm7173 | Tsx | Gm17361 | Ott | Gm15726 |
| Gm26775 | Gm26992 | Gm21616 | Gm15092 | Gm15247 |
| 4930595M18Rik | 1700121L16Rik | Astx4c | Gm15093 | Gm21887 |
| Tsga8 | 5330434G04Rik | Gm17693 | Gm15100 | 1110015O18Rik |
| Gm14764 | Cypt2 | Astx1c | Gm15085 | Gm17308 |
| Gm14762 | Tlr13 | Gm17522 | Gm15086 | 2700069I18Rik |
| 5430427O19Rik | Fnd3c2 | Astx4d | Gm10439 | 4930555M17Rik |
| Samt3 | Fndc3c1 | Gm17267 | Gm15097 | Gm8797 |
| Gm27000 | Gm5127 | Astx3 | Gm15091 | Gm37350 |

| | | | | |
|---|---|---|---|---|
| 1700008P02Rik | Gm1527 | Gm42921 | Gm21958 | Gm38353 |
| Gm16685 | Gm33206 | Gm20755 | 4931419H13Rik | 4930509J09Rik |
| 1700010I02Rik | Mannr | Gm43821 | Spg20 | Gm20356 |
| Gm32496 | Mecomos | 5430434I15Rik | Gm42609 | Platr10 |
| C230057A21Rik | Gm38258 | Gm42920 | Gm21954 | Gm29133 |
| 4930539M17Rik | Gm15462 | Gm36823 | Gm43549 | Gm37256 |
| C030034L19Rik | 4933429H19Rik | Gm43538 | Gm43376 | 4921511C10Rik |
| Gm16337 | Gm15496 | C230034O21Rik | Gm26671 | 6430573P05Rik |
| Gm38335 | Gm38025 | Gm43539 | Gm40055 | Gm17359 |
| Gm9833 | Gm37592 | 1700017G19Rik | 4921539H07Rik | 4930589L23Rik |
| Gm37389 | Gm37640 | 1700034I23Rik | 1700007F19Rik | Gm36569 |
| Gm38303 | Gm37459 | Gm40038 | 4930593A02Rik | Fam198b |
| Gm10745 | 4930419G24Rik | Gm16508 | Gm43570 | A330069K06Rik |
| Gm2464 | Gm43666 | 1700052H01Rik | Gm38186 | Gm37892 |
| Gm2464.1 | Gm43140 | Gm40040 | Gm5538 | Gm37971 |
| Slc7a12 | 4930431L21Rik | Platr4 | Gm8298 | Gm42468 |
| 1810022K09Rik | Gm43077 | 2400006E01Rik | C130079G13Rik | Gm42476 |
| A930001A20Rik | Gm38505 | Gm31266 | Gm38326 | Gm16000 |
| Gm33819 | Gm34599 | 1700027H10Rik | Gm37035 | Gm43346 |
| Gm9733 | Gm20515 | 1700018B24Rik | Gm37696 | A830029E22Rik |
| Gm37468 | Gm29135 | Gm37190 | B430305J03Rik | Gm30043 |
| Gm30340 | Gm42695 | Gm31415 | 9330121J05Rik | Gm43348 |
| Mir124-2hg | Gm42205 | Gm20557 | Gm34240 | Rbm46os |
| Cypt12 | Gm3143 | Gm37854 | Gm34302 | Gm30097 |
| Gm30667 | Gm38509 | Gm29230 | Gm26850 | Gm10710 |
| 4930433B08Rik | Gm21388 | Gm38246 | Vmn2r1 | Gm38096 |
| Gm16093 | 1700017M07Rik | Gm5103 | Vmn2r2 | 1700028M03Rik |
| 1700064H15Rik | Gm6639 | Gm37229 | Vmn2r3 | Gm26771 |
| 4632415L05Rik | Gm15952 | Gm20089 | Vmn2r4 | 4930565D16Rik |
| Gyg | Mccc1os | Gm37202 | Vmn2r5 | Gm37240 |
| Gm43525 | Ccdc144b | Gm30074 | Vmn2r6 | 1700036G14Rik |
| Gm31466 | Gm43079 | Gm37170 | Vmn2r7 | Gm42813 |
| Gm29137 | D3Ertd254e | Gm20402 | Vmn2r7.1 | Gm45790 |
| Gm31693 | Gm43255 | Gm30173 | A330015K06Rik | Gm42812 |
| Gm43674 | Gm42208 | Gm2447 | A730090N16Rik | Fam160a1 |
| Gm43672 | 1810062G17Rik | Gm30292 | Gm37359 | Glt28d2 |
| Gm38324 | Gm11548 | Gm20750 | Gm37305 | Gm37933 |
| Gm37021 | Gm11549 | Gm42901 | Gm17402 | Gm37876 |
| 4930412E21Rik | 4932438A13Rik | Lhfp | Gm35299 | Kirrel |
| A830092H15Rik | Gm12531 | Gm43803 | 4930402C01Rik | Gm37855 |
| 1700125G22Rik | Gm12532 | Gm16206 | Gm21949 | Fcrls |
| Gm42196 | Gm12534 | Gm30735 | 4930535E02Rik | Rrnad1 |
| 1700112D23Rik | Fgf2os | C820005J03Rik | Gm35584 | 1700113A16Rik |
| Gm32950 | Spata5 | Gm10985 | Gm1647 | AI849053 |
| Gm33051 | 4930594O21Rik | Gm31026 | Gm6634 | Gm38392 |
| Gm37867 | Gm5148 | Gm26678 | Gm20754 | Gm37584 |

| | | | | |
|---|---|---|---|---|
| Gm20652 | Gm5286 | 4930573H18Rik | Gm35507 | Gm43608 |
| Gm43714 | Gm4778 | Olfr1402 | Gm27008 | Gm43607 |
| 2810403A07Rik | Gm38411 | Gm17651 | AI504432 | Gm40190 |
| Gm17146 | Gm43401 | Gm31305 | A930002I21Rik | Gm4610 |
| Gm43713 | 1700040D17Rik | Gm15999 | A630076J17Rik | Gm42836 |
| Gm43738 | Gm10972 | Gm5544 | Gm10961 | Gm43569 |
| Gm16069 | Gm15234 | Gm4450 | Gm36211 | Gm43568 |
| Fam189b | B230398E01Rik | Gm10681 | 4933431E20Rik | 4933405D12Rik |
| Gba | A730011C13Rik | 5730437C11Rik | Gm12498 | Gm10959 |
| Gm15998 | Gm15265 | Gm12440 | Gm12500 | 1810037I17Rik |
| 4731419I09Rik | Gm16740 | Gm43120 | Gm12524 | Gm35065 |
| Gm15417 | 6330562C20Rik | Spag17os | Gm40123 | Gm43731 |
| 4632404H12Rik | 4930558C23Rik | Gm43121 | 1700010K24Rik | 1700006A11Rik |
| Gm42809 | Gm4349 | Fam46c | Gm12523 | Gm42823 |
| Gm19710 | Gm26594 | Gm12474 | Gm12525 | 1700003H04Rik |
| 4930537H20Rik | C920021L13Rik | Gm43464 | Sars | Gm42825 |
| Gm16048 | Gm17690 | Gm42538 | 5330417C22Rik | Gm42827 |
| Gm42674 | Gm15444 | Gm43135 | 1700013F07Rik | Gm17225 |
| 9130204L05Rik | Hist2h2ab | Gm43134 | Gm27244 | Gm15551 |
| Lor | Hist2h2ac | Gm16160 | Gm9857 | Gm16958 |
| Gm43736 | Hist2h2be | Gm43241 | Fam102b | Gm43729 |
| Gm43735 | Hist2h3c2 | A230001M10Rik | Gm43221 | Gm43653 |
| Gm35439 | Gm20632 | Gm43242 | 4930408K08Rik | Gm35585 |
| Sprr2a1 | Hist2h2aa2 | Gm42678 | Slc25a54 | 9830132P13Rik |
| Sprr2a2 | Gm20633 | Gm42682 | C130013H08Rik | Gm35667 |
| Sprr2a3 | Hist2h2aa1 | Tspan2os | Gm6602 | Gm43824 |
| Sprr2b | Hist2h3c1 | Gm42681 | Gm9889 | 4933424H11Rik |
| Sprr2f | Hist2h4 | Gm43062 | Gm43109 | Gm28543 |
| Sprr2h | Gm20628 | Gm10964 | Gm26544 | Gm35986 |
| Sprr2j-ps | Hist2h3b | Gm15886 | Gm31651 | Gm42697 |
| Gm9774 | Gm42743 | Gm43064 | Gm29151 | Gm42650 |
| Sprr2k | Hist2h2bb | Phtf1os | Gm26530 | 6330410L21Rik |
| 2310046K23Rik | Platr30 | Gm5546 | Gm42893 | Gm42997 |
| Gm38119 | Gm26553 | Gm38412 | Gm42457 | Gm43522 |
| Gm38201 | Gm17575 | Gm29561 | Gm43191 | 2010110G14Rik |
| 2310050C09Rik | Gm26689 | Fam19a3 | 4930512P04Rik | Gm36520 |
| Lce1m | 4930477E14Rik | Gm40117 | 4930455H04Rik | Gm43351 |
| Gm4858 | Gm26654 | 4930564D02Rik | 1700061I17Rik | Gm43349 |
| Flg | BC107364 | Kcnd3os | Gm42891 | Gm43352 |
| Gm37245 | Gm10685 | Fam212b | 4921521D15Rik | 4930534D22Rik |
| Gm37596 | Hfe2 | Gm43848 | Gm9916 | Gm29865 |
| Gm5773 | Gm15441 | Gm5547 | 6530403H02Rik | Gm42880 |
| Gm9117 | Gm16253 | Atp5f1 | Tmem56 | Gm29811 |
| Gm9125 | Gm42957 | Pifo | A530020G20Rik | Gm43254 |
| Gm10697 | 4930442L01Rik | Gm42722 | A730020M07Rik | Gm42567 |
| Gm10696 | 9230114J08Rik | Olfr266 | 4930432M17Rik | Gm40153 |

| | | | | |
|---|---|---|---|---|
| Gm6135 | Gm33758 | Gm42947 | 4930480G23Rik | Gm21953 |
| Gm30484 | Gm42704 | 4930570G19Rik | 4930598A11Rik | Fam205a3 |
| Gm26691 | Gm6260 | Gm15577 | Gm11872 | Gm21586 |
| Gm43192 | Gm42705 | 4930592C13Rik | Gm30731 | Gm2163 |
| Gm30648 | Gm42706 | Gm33466 | 1700025O08Rik | Fam205a2 |
| 4930539C22Rik | Gm42707 | 1810013D15Rik | Gm11884 | Gm10593 |
| Gm26820 | Gm42708 | B230334C09Rik | Gm28448 | Gm10591 |
| 4930539J05Rik | Gm34078 | Gm20752 | Gm27243 | Gm2564 |
| Gm31243 | Gm43446 | Vmn1r2 | Gm11899 | 4930578G10Rik |
| 1700030L20Rik | Gm43447 | Vmn1r3 | 4930548K13Rik | Gm12395 |
| Gm43357 | Gm34866 | AI838599 | Gm11915 | Gm12394 |
| Gm43356 | Gm43240 | 2210414B05Rik | Gm11917 | Fam205a1 |
| Gm567 | Gm16233 | Gm26857 | Gm11916 | Fam205c |
| Gm4861 | 9530034A14Rik | Gm11808 | Gm11924 | 1700022I11Rik |
| Gm21962 | Cyr61 | Impad1 | 4930556G01Rik | Gm26643 |
| Gm42610 | Gm17501 | Gm11782 | Bach2os | Fam214b |
| Gm31678 | Gm35409 | 4930423M02Rik | Gm12753 | Gm26881 |
| Gm43403 | Wdr63 | 4930430E12Rik | Gm12364 | Fam166b |
| H2afz | Gm10636 | Gm11802 | 3110043O21Rik | AL732506.1 |
| Gm40155 | Gm16325 | 8430436N08Rik | Gm12371 | Gm12454 |
| Gm43689 | 4930503B20Rik | Gm11795 | Gm12374 | Gm12472 |
| Gm19708 | Gm29771 | Gm26548 | Gm12381 | Gm12473 |
| Gm5105 | Gm36831 | Gm11816 | Gm17094 | Olfr70 |
| Gm43691 | Gm37041 | Gm11817 | Ddx58 | Olfr71 |
| 0610031O16Rik | Gm36888 | 1700123O12Rik | 4930509K18Rik | Olfr159 |
| Gm16559 | Gm43573 | Gm11815 | 4933428C19Rik | Olfr156 |
| Adh6a | Gm10287 | 2610301B20Rik | Gm6297 | Olfr157 |
| Adh6b | Gm30382 | Gm11827 | AI464131 | Olfr155 |
| 4930425O10Rik | Gm5149 | Gm26663 | 1110017D15Rik | Gm12462 |
| Gm32693 | Gm31121 | Gm11831 | Fam219aos | Gm12678 |
| Gm43678 | Gm43470 | Gm11832 | Dnaic1 | Gm12493 |
| Gm43049 | Gm43468 | Gm10604 | Gm12406 | Gm12679 |
| Gm32754 | 4930555A03Rik | Gm26895 | Gm12405 | 1700055D18Rik |
| 4930590L14Rik | Gm42949 | Gm11840 | Gm13307 | Frmpd1os |
| Gm15688 | Gm43617 | Gm11842 | Gm21541 | Gm12408 |
| Gm15689 | Gm43619 | Fam92a | Gm20878 | Gm829 |
| Gm15540 | Gm31881 | Gm11846 | Fam205a4 | Gm12410 |
| 9530052C20Rik | AI115009 | Gm11820 | Gm21093 | Gm12446 |
| 4930519L02Rik | Gm16213 | Gm11823 | Gm3892 | Gm568 |
| Gm43614 | Gm16231 | 1700120G11Rik | Gm3893 | Gm12426 |
| A830019L24Rik | 1700094M23Rik | Tmem55a | Gm13304 | Gm12434 |
| Gm43613 | 5730460C07Rik | Gm11844 | Gm13299 | Gm12436 |
| Gm43612 | Asb17os | Gm11861 | Gm13306 | Acnat2 |
| 1700001N15Rik | Gm43400 | Gm11851 | Gm13305 | Acnat1 |
| Gm43616 | Gm20389 | Gm12353 | Gm10600 | Tmem246 |
| Gm33651 | 9330178D15Rik | Gm12354 | Gm21980 | C630028M04Rik |

| | | | | |
|---|---|---|---|---|
| Smc2os | Mup12 | Gm12914 | Gm12694 | Gm12802 |
| Vma21-ps | Gm21320 | Gm11264 | Junos | Hspb11 |
| Olfr275 | Mup13 | Gm5860 | 4930551L18Rik | Gm12869 |
| Olfr273 | Mup14 | Gm11266 | Gm12708 | 1700047F07Rik |
| Olfr272 | Mup15 | Gm11269 | Gm29064 | Gm12907 |
| Olfr270 | Mup16 | Gm11413 | Gm10192 | 4933424M12Rik |
| 4930522O17Rik | Mup17 | Gm11184 | E130114P18Rik | Lrp8os2 |
| AI427809 | Mup18 | Gm11415 | 0610025J13Rik | Lrp8os1 |
| Gm12496 | Mup19 | 6030471H07Rik | Kank4os | 0610037L13Rik |
| 1700060J05Rik | Mup5 | Gm12414 | I0C0044D17Rik | 4930407G08Rik |
| Gm12484 | Mup20 | Gm12415 | Gm12705 | Fam159a |
| Gm29067 | Mup3 | Saxo1os | Gm12681 | Zcchc11 |
| Gm12478 | Gm12909 | Gm12600 | Gm12689 | Gm17354 |
| Gm12480 | Gm12910 | Gm12631 | Gm12690 | 3110021N24Rik |
| Gm12514 | Mup21 | C87499 | Gm12688 | A730015C16Rik |
| Gm12505 | Gm11209 | Gm26566 | Gm12682 | 8030443G20Rik |
| Gm12510 | Gm11210 | Gm13274 | Gm12702 | Nrd1 |
| Gm12511 | Gm11211 | Gm13271 | Ube2uos | Ttc39aos1 |
| Gm12506 | Aknaos | Gm16686 | Gm10577 | Dmrta2os |
| Gm12519 | Gm11213 | Gm13283 | Gm12801 | Skint8 |
| 2310081O03Rik | Gm11482 | Gm13290 | Gm12796 | Skint7 |
| Gm26657 | Gm11214 | Gm13289 | E130102H24Rik | Skint1 |
| Ikbkap | Gm11216 | Gm13272 | 0610043K17Rik | Skint4 |
| Fam206a | Gm11217 | Ifnz | Gm12798 | Skint3 |
| Gm12530 | Gm11751 | Gm26525 | B020004J07Rik | Skint9 |
| Gm12536 | Gm12911 | Gm13276 | C130073F10Rik | Skint2 |
| 1700042G15Rik | Gm11228 | Gm13277 | Gm12800 | Skint10 |
| Palm2 | Gm11232 | Gm13278 | Gm12794 | Skint6 |
| Akap2 | Gm11234 | Gm13275 | Gm12790 | Gm12823 |
| Gm26889 | Gm11250 | Gm26867 | Gm12789 | Skint5 |
| Gm12580 | Aldoart1 | Gm13279 | Gm12709 | Skint11 |
| Olfr267 | Gm11487 | Gm13285 | Tctex1d1 | Gm28864 |
| AI314180 | Gm11238 | Gm13287 | BB031773 | Gm12829 |
| Gm20503 | Gm11237 | Gm13288 | Wdr78 | 9130410C08Rik |
| AI481877 | Gm11236 | 4930553M12Rik | Gm44037 | Gm12830 |
| Gm12596 | Gm11239 | Wincr1 | 4930456L15Rik | Cyp4x1os |
| Gm12542 | Gm428 | Gm12606 | Gm12718 | Gm12847 |
| Mup4 | Gm11758 | Gm12629 | Gm17662 | Gm29434 |
| Mup6 | Gm11757 | Gm12668 | Gm12728 | 6430628N08Rik |
| Mup7 | Gm13871 | Gm12670 | Gm12729 | 1700042G07Rik |
| Mup2 | Gm11756 | 4930577H14Rik | Gm12730 | Gm49337 |
| Mup8 | 2310002L09Rik | Gm12666 | 2700068H02Rik | Gm12951 |
| Mup9 | A230083N12Rik | Gm12637 | Gm12744 | Gm12953 |
| Mup1 | Gm42303 | Gm12649 | Lexm | Gm12996 |
| Mup10 | Gm11261 | Gm12648 | Gm12746 | 1700021J08Rik |
| Mup11 | Gm11412 | Gm10306 | Gm12786 | Tctex1d4 |

| | | | | |
|---|---|---|---|---|
| Gm12843 | Gm12916 | Gm10300 | Gm13029 | Pramef20 |
| Gm12828 | 1700021L23Rik | Gm28874 | Gm13016 | Pramef12 |
| Gm12827 | 4933435F18Rik | Gm28872 | Gm13017 | Pramef12os |
| Gm12840 | 1700057H15Rik | Gm17300 | Gm13021 | 1700012P22Rik |
| Gm12845 | Gm2164 | Gm12981 | Gm13026 | Gm9944 |
| Gm12841 | 4933407E24Rik | Atpif1 | Gm13025 | 9430007A20Rik |
| 9530034E10Rik | 3100002H09Rik | Gm12999 | 4930515B02Rik | Gm13124 |
| Gm12853 | 1700125G02Rik | Gm26615 | Gm13031 | Gm436 |
| Olfr62 | 9930104L06Rik | Gm13257 | 4921514A10Rik | Gm13178 |
| Olfr1342 | Gm12930 | Fam46b | Rsg1 | Gm438 |
| Olfr1341 | Gm12932 | Gm12974 | Gm13055 | Zfp990 |
| Olfr1340 | 1700041M05Rik | Gm12977 | Gm13056 | Gm13212 |
| Olfr1339 | 2610028E06Rik | Gm7534 | Gm13074 | Gm26763 |
| Olfr1338 | Gm12946 | E130218I03Rik | Gm13075 | Zfp980 |
| Olfr1337 | Adprhl2 | Grrp1 | AI507597 | Gm13236 |
| Olfr1335 | 4930471C06Rik | 1700021N21Rik | Ctrcos | Zfp986 |
| Olfr1333 | 1700080G11Rik | Tmem57 | Gm29367 | Zfp987 |
| Olfr1331 | A630031M04Rik | Gm16225 | Fhad1os2 | Gm26573 |
| Olfr1330 | 1700112K13Rik | Gm16224 | Fhad1os1 | Gm13165 |
| Olfr1329 | Csmd2os | Gm12984 | 4930455G09Rik | Gm13166 |
| Olfr1328 | Hmgb4os | 1700029M20Rik | Tmem51os1 | Zfp992 |
| Gm12863 | Gm12958 | Gm13008 | Gm13062 | Zfp981 |
| Lao1 | Gm12968 | Gm27022 | Gm13052 | Gm26880 |
| Gm12865 | Gm15904 | 6030445D17Rik | Pramel1 | 1700095A21Rik |
| Gm12862 | 1700086P04Rik | 1700037C06Rik | Pramef8 | Zfp993 |
| Gm12866 | Gm15906 | Gm13001 | Oog4 | Gm21411 |
| Gm12867 | Yars | Gm13003 | BC080695 | Zfp989 |
| Gm12868 | C77080 | 2810405F17Rik | Gm13057 | C230088H06Rik |
| Gm12927 | Gm12976 | Gm13010 | Gm13083 | Rex2 |
| AL607142.1 | Gm26722 | 1700013G24Rik | Gm13088 | 4933438K21Rik |
| Foxo6os | Gm12979 | Rap1gapos | Gm13089 | Zfp991 |
| Ctps | E330017L17Rik | Gm13012 | Gm13078 | Gm26624 |
| Gm8439 | 1700003M07Rik | 1700095J12Rik | Gm13023 | Zfp988 |
| Gm12860 | Gm12963 | 2310026L22Rik | Gm13084 | Zfp978 |
| Gm45579 | Gm853 | AB041806 | Gm13103 | Gm16503 |
| Gm12888 | Gm10570 | Gm13030 | Pramef6 | Zfp985 |
| Gm12886 | Gm26516 | Gm45533 | Pramef25 | Zfp979 |
| Gm12887 | Gm12972 | Minos1 | Gm13101 | Gm16211 |
| 9530002B09Rik | Gm12970 | Gm16287 | Pramef17 | Zfp534 |
| Gm12923 | Gm26716 | Pqlc2 | Pramel4 | Zfp984 |
| Gm12925 | Gm12973 | 2700016F22Rik | Gm13102 | Gm20707 |
| 1700121C08Rik | A930031H19Rik | Gm21969 | Oog3 | Fv1 |
| 4933427I04Rik | Gm831 | Gm1667 | Oog2 | Gm13054 |
| Gm26606 | Gm12962 | Gm13028 | C87977 | Gm13201 |
| Gm12905 | Gm16080 | 4933427I22Rik | Gm13128 | Gm13206 |
| 4930535I16Rik | Gm13063 | Gm13027 | Gm13119 | Gm13200 |

| | | | | |
|---|---|---|---|---|
| Gm13209 | Mxra8os | Gm33847 | 5031425E22Rik | Gm29609 |
| Gm572 | Gm16008 | Gm8922 | 4930580E04Rik | Gm9970 |
| Gm13203 | AW011738 | 1700003C15Rik | 2700038G22Rik | 4930548H24Rik |
| Gm15969 | Gm47252 | Gm8926 | Fam126a | 4930566F21Rik |
| Gm13205 | Vmn2r125 | Gm43132 | Nupl2 | Gm43660 |
| Gm17029 | Gm36548 | Gm6465 | Gm15589 | AI839979 |
| Ube4bos3 | Gm17590 | Gm43131 | Gm15587 | Gm17130 |
| Ube4bos2 | Gm42435 | 4933402N22Rik | 2310074N15Rik | Gm36840 |
| Ube4bos1 | 1700109H08Rik | Gm29066 | 4931409K22Rik | Gm7596 |
| Gm13097 | 2510017J16Rik | Gm8953 | Gm26648 | Gm15614 |
| Gm13066 | Gm43825 | 4930568A13Rik | Prkag2os2 | Gm20671 |
| Gm13068 | 4833413G10Rik | Speer3 | Prkag2os1 | Gm20465 |
| Gm16188 | Gm42744 | Gm4128 | 1500035N22Rik | Gm43851 |
| Gm13073 | 1700015F17Rik | Gm26918 | E130116L18Rik | Gm9903 |
| Gm13070 | Gm8773 | Gm26954 | 4831440E17Rik | Gm42849 |
| Gm13067 | Gm26825 | Gm9758 | 1700096K18Rik | Gm42851 |
| Gm13093 | Gm40263 | Speer4e | Gm10062 | Gm1673 |
| Gm13091 | Gm30835 | Gm10354 | Gm43972 | 4930557J02Rik |
| Gm16079 | Gm15731 | Gm17019 | Gm21663 | Gm42846 |
| Gm13049 | Gm5106 | Gm43391 | Gm21671 | Gm42848 |
| 1700045H11Rik | Gm15611 | Gm21149 | Gm21680 | Gm43458 |
| Gm13090 | Gm40264 | Gm21847 | Gm21698 | Gm15522 |
| 4930589P08Rik | Gm15734 | Gm21190 | Gm1979 | Gm42560 |
| Gm16333 | 9330182L06Rik | Gm30613 | Gm5862 | Gm43791 |
| Gm833 | Gm10482 | Gm21083 | Speer4a | 4930478P22Rik |
| Gm13173 | Gm5152 | Speer4d | Gm7347 | E130018O15Rik |
| Gm13174 | Gm31831 | 4930572O03Rik | Gm10471 | 4931431C16Rik |
| Gm13115 | Gm6455 | Speer4cos | 5031410I06Rik | Gm15650 |
| BC039966 | Gm43358 | Speer4c | Gm10220 | 2210406O10Rik |
| Trp73os | Gm8871 | 4930519H02Rik | Gm7361 | Gm43701 |
| Tprgl | Gm8857 | Gm3510 | 4930584F24Rik | Ppp2r2cos |
| B230104I21Rik | Gm43492 | Gm43000 | Speer4b | Gm26647 |
| Gm13133 | 4930558F17Rik | Gm28710 | Gm16058 | Jakmip1.1 |
| Gm13134 | Gm8879 | Gm28710.1 | Gm26608 | Gm43763 |
| Gm13111 | Gm32554 | Speer4f2 | 9530036O11Rik | Gm20052 |
| Gm27202 | Gm5861 | Speer4f1 | Gm26894 | Gm40289 |
| Gm13112 | Gm33093 | Gm31752 | 4632411P08Rik | Gm26560 |
| Fam213b | Gm8890 | 4921504A21Rik | Gm5129 | 4930487D11Rik |
| Gm10564 | Gm32877 | A630072M18Rik | Gm42450 | Clnkos |
| Gm27200 | Speer1 | Gm10475 | A230098N10Rik | Gm40293 |
| Gm10563 | 1700108N06Rik | Gm16110 | Gm4961 | Gm40292 |
| Gm16024 | Gm8897 | Gm43657 | 3110082J24Rik | Gm42524 |
| 1500002C15Rik | 4930437M23Rik | 6030443J06Rik | 1700001C02Rik | 4930513D17Rik |
| Atad3aos | Gm8906 | A930003O13Rik | Gm42765 | Gm35191 |
| Gm26840 | Gm43640 | Gm42948 | Gm15461 | Gm16223 |
| Tmem88b | Gm6460 | Gm42951 | Gm15469 | Gm43700 |

| | | | | |
|---|---|---|---|---|
| Gm42711 | Gm15819 | Gm43101 | 1700063O14Rik | A730035I17Rik |
| Gm7854 | 5830416I19Rik | Gm43100 | Gm43688 | 1700007G11Rik |
| Gm43020 | Gm43838 | Gm19590 | Gm26582 | Gm11111 |
| Gm43699 | Gm43837 | Gm28865 | U90926 | 1700010H22Rik |
| Gm43021 | Gm31363 | Gm32780 | Gm43599 | Gm43251 |
| Gm15866 | 1700027F09Rik | Gm7467 | 4932430I15Rik | Gm35394 |
| Gm42555 | C030018K13Rik | Gm7271 | 11-Sep | Gm35172 |
| Gm16015 | Gm3716 | C530008M17Rik | 2010109A12Rik | 4930405H06Rik |
| Gm7879 | Gm42564 | 2310040G07Rik | Gm16427 | Gm16226 |
| Gm16401 | Gm42565 | 1700112J05Rik | C87414 | 4930522N08Rik |
| Gm42428 | Gm26761 | Thegl | Gm7792 | Gm38413 |
| Gm42982 | Gm43552 | Hopxos | Gm16429 | Gm35911 |
| Gm42984 | Gm42726 | Gm15831 | BC061212 | Gm17092 |
| Gm36017 | 9130230L23Rik | Gm34648 | Gm6351 | 4930524J08Rik |
| Gm43507 | Gm26725 | Gm34728 | Gm3106 | Tmem150cos |
| Gm42750 | 1700126H18Rik | 1700017L05Rik | A430089I19Rik | Gm9932 |
| 4930431F12Rik | Uchl1os | Gm42761 | Gm6502 | 2310034O05Rik |
| D5Ertd615e | Gm15949 | Gm16054 | AA792892 | Gm15777 |
| Gm43303 | Gm15948 | 1700031L13Rik | Gm7682 | 4930458D05Rik |
| Gm42413 | Gm42670 | Gm43532 | Gm3139 | Gm36793 |
| 9630001P10Rik | Gm43698 | Gm43567 | Gm3147 | Gm20484 |
| 4930405L22Rik | Gm26756 | Gm43056 | Gm6509 | Gm42133 |
| 4930518C09Rik | Gm15478 | Gm16579 | Gm6205 | 9430085M18Rik |
| 5730480H06Rik | Gm15477 | Gm43055 | Gm3183 | Gm20548 |
| Gm20700 | Gm21905 | Gm43057 | Gm6367 | Gm29707 |
| 4930448I18Rik | 4930425K10Rik | Gm43638 | Gm7942 | 1700013M08Rik |
| Gm43016 | Gm5108 | Gm42794 | Gm16513 | 4930429D17Rik |
| Gm43177 | Gm38562 | 1700066N21Rik | Gm10424 | 1700021F02Rik |
| C130083M11Rik | Gm20647 | BC037156 | Gm3259 | 5430427N15Rik |
| Gm43179 | AU023070 | BC051076 | D5Ertd577e | Gm42620 |
| Gm3519 | Gm5868 | 2310003L06Rik | Gm3286 | Gm17660 |
| Gm43685 | 4933408A14Rik | Gm28434 | Gm7971 | Gm26703 |
| 8030423F21Rik | Gm10135 | Gm11116 | Gm7978 | Gm31048 |
| Gm30301 | 1700071G01Rik | Gm11115 | Gm7982 | BC005561 |
| Gm17182 | Gm34411 | Gm7714 | E330014E10Rik | D930016D06Rik |
| Gm45495 | 1700025M24Rik | Prol1 | Gm33050 | Gm43302 |
| Gm42745 | 1700019F05Rik | Gm42912 | Gm42604 | Gm26519 |
| Gm10441 | C78283 | Gm9958 | 1700016F12Rik | Gm32051 |
| Gm40304 | Usp46os1 | Gm43449 | Gm42601 | Lrrc8dos |
| Gm10440 | Gm15984 | 5830473C10Rik | Gm33370 | Gm32736 |
| 4930459L07Rik | Gm15985 | Cxcl15 | Gm8013 | Gm32921 |
| 1700029E06Rik | Gm42575 | Gm43086 | Gm9484 | 4930458A03Rik |
| Gm43042 | Gm6116 | Gm42530 | Gm33609 | 4930432H08Rik |
| 4933402J10Rik | Gm19583 | Gm19610 | Gm2861 | Gm42916 |
| 1700014F14Rik | Gm42802 | Gm19619 | Gm33969 | Gm26872 |
| 0610040J01Rik | Gm42800 | Gm28271 | 4930467D21Rik | Gm28050 |

| | | | | |
|---|---|---|---|---|
| Gm29464 | A630023P12Rik | 4930430O22Rik | Gm42878 | Gm42980 |
| Gm17304 | Gm42595 | Gcn1l1 | Gm16552 | Gm42817 |
| Gm33474 | Gm15559 | 1110006O24Rik | 1700008B11Rik | Gm42495 |
| Gm42902 | Gm26515 | Gm14508 | Fam109a | Gm42500 |
| Gm17202 | Gm43118 | Gm13842 | Gm3970 | 4930573I07Rik |
| A830010M20Rik | C130026L21Rik | C330018A13Rik | Gm15637 | 5930412G12Rik |
| Gm42669 | Gm42489 | Gm14507 | Gm43301 | Gm42875 |
| A930041C12Rik | Gm26897 | Gm13837 | A930024E05Rik | Gm43001 |
| Gm26692 | E130006D01Rik | Gm13839 | Gm2479 | Zfp11 |
| 4930428O21Rik | Gm16019 | Gm13838 | Gm26745 | 14-Sep |
| Ube2d2b | Gm36535 | B230112J18Rik | Gm38102 | Nupr1l |
| Fam69a | Gm42488 | 4930562A09Rik | 4932422M17Rik | Gm6598 |
| 9330198I05Rik | 1700016B01Rik | Srrm4os | Gm43409 | A330070K13Rik |
| Gm42518 | 1700028D13Rik | Gm43122 | Wdr66 | Gm29681 |
| Gm42517 | Gm26953 | Gm29926 | Gm15857 | 2810432F15Rik |
| Gm10419 | Gm6583 | 9530046B11Rik | Gm15860 | 4930563F08Rik |
| Atp5k | Gm20636 | BC051077 | Gm15747 | Gm42439 |
| Mfsd7a | Gm6588 | Gm10399 | Gm49027 | Gm28616 |
| 1700047L14Rik | Gm27004 | Gm15728 | Gm43518 | Gm42626 |
| Vmn2r8 | 1700034G24Rik | Gm26995 | Gm34086 | Gm36667 |
| Vmn2r9 | Gm42864 | Gm9754 | Gm43661 | Gm15627 |
| Gm20629 | Gm42865 | 2410131K14Rik | Gm16001 | Gatsl2 |
| Vmn2r10 | 1700095B10Rik | Gm28563 | Pitpnm2os2 | Gm42882 |
| Vmn2r11 | Gm42161 | Gm43782 | Pitpnm2os1 | Gm42884 |
| Vmn2r12 | Tmem211 | Gm43342 | Gm42425 | Gm10369 |
| Vmn2r13 | Aym1 | 4930413E15Rik | 2810006K23Rik | Gm30003 |
| Vmn2r14 | Gm15736 | 1700081H04Rik | 4930404A12Rik | 4933439J24Rik |
| Vmn2r15 | 1700069L16Rik | Gm7538 | Gm43777 | Wbscr25 |
| Vmn2r16 | Gm17122 | Gm31314 | Gm42838 | Gm43500 |
| Vmn2r17 | Gm42797 | Tbx3os1 | Gm42839 | Gm43091 |
| 5430403G16Rik | Gm13790 | Gm16063 | Gm32585 | Srrm3os |
| 4930522L14Rik | Gm42789 | Gm16064 | Gm40323 | Gm15701 |
| A430073D23Rik | 4930515G01Rik | Tbx3os2 | Gm10382 | Gm43604 |
| Gm26779 | 1500011B03Rik | 1700021F13Rik | 1700048F04Rik | Gm16599 |
| Gm26808 | Gm20499 | Gm43050 | Gm42945 | 4933404O12Rik |
| 1010001B22Rik | 2610524H06Rik | Gm43269 | Gm42952 | Muc3 |
| Zfp932 | 4930519G04Rik | Gm38554 | Gm40330 | Gm3054 |
| Gm17655 | Gm9936 | Gm10390 | 1700081B01Rik | Gm40348 |
| Gm35315 | Gm13822 | Gm42654 | Gm33347 | Gm42456 |
| Gm43136 | Hnf1aos1 | Gm27199 | Gm42953 | 9130604C24Rik |
| Gm43137 | Hnf1aos2 | Gm15690 | Gm40331 | Gm8066 |
| Gm26718 | Gm10401 | Gm43579 | 4933438B17Rik | Gm17112 |
| Gm15788 | Gm13830 | Gm15749 | Gm43475 | Gm36266 |
| Gm15792 | Gm42903 | Gm43069 | 4930572K03Rik | Gm20605 |
| Gm26711 | 4930401G09Rik | Gm42917 | Gm43476 | Gm15498 |
| Gm42596 | Gm13832 | Gm42918 | Tmem132cos | Gm36551 |

| | | | | |
|---|---|---|---|---|
| Gm43495 | 1700018F24Rik | Gm21221 | Gm15592 | Gm13860 |
| Smok3a | Atp5j2 | Gm43450 | Gm15594 | Gm13861 |
| Smok3b | Gm5565 | Gm43452 | Gm26809 | Npn2 |
| Smok3c | Gm6309 | Gm42557 | Gm20186 | Gm14547 |
| Gm454 | 1700001J03Rik | Gm42556 | 4930524B17Rik | 3110062M04Rik |
| Gm43645 | B230303O12Rik | Gm5 | Gm30270 | 1810058I24Rik |
| Nxpe5 | Gm43625 | Gm42529 | Gm3289 | 1700065J11Rik |
| BC037034 | Gm6408 | Gm43690 | Gm26719 | Gm42931 |
| 6330418K02Rik | Gm6370 | Gm43598 | Gm20756 | 9330158H04Rik |
| Gm43720 | 4930449I24Rik | Gm43597 | 6530409C15Rik | 1700111E14Rik |
| Gm10874 | Gm3402 | Gm36447 | Gm3294 | Ttc26 |
| Zfp157 | Gm3404 | Gm43298 | Fam71f2 | Gm26699 |
| 1700123K08Rik | Gm3409 | Gm42906 | Fam71f1 | Gm42962 |
| Zfp68 | Gm3415 | 5430435K18Rik | Gm17172 | 1700025N23Rik |
| A430033K04Rik | Gm34333 | D730045B01Rik | Gm9047 | 4930599N23Rik |
| Gm5294 | 4930573C15Rik | Vmn2r18 | Gm26627 | Gm43479 |
| 6330403L08Rik | 1700041I07Rik | Gm43129 | Smkr-ps | Gm10244 |
| 4930520M14Rik | Gm27033 | Gm8579 | Rncr4 | Gm16272 |
| 3110082I17Rik | Gm26597 | 4930528H21Rik | 1700095J07Rik | Gm42420 |
| C130050O18Rik | D5Ertd605e | Gm20714 | 1700012C14Rik | Gm26833 |
| 4930500L23Rik | Gm43156 | 1700019G24Rik | Gm13781 | Gm26807 |
| Gm26938 | Gm15721 | Gm20619 | Gm13782 | E330009J07Rik |
| Gm42423 | Gm36186 | Gm20617 | Gm44296 | Olfr461 |
| Gm16122 | 2210417A02Rik | Gm20685 | 4930412F09Rik | Olfr460 |
| Gm16121 | 4930505K14Rik | Gm20618 | Gm27019 | Trbv1 |
| Gm16120 | Gm15407 | Gm26774 | Gm13833 | 1700074P13Rik |
| Gm4869 | Gm42791 | Gm19272 | Gm14532 | Gm4744 |
| Gm42808 | Gm15410 | 1700012J15Rik | Mkln1os | Gm29588 |
| Gm43703 | Gm15406 | Gm45062 | Gm13844 | 1810009J06Rik |
| Gm16035 | 8430423G03Rik | Gm16055 | Gm6117 | Gm2663 |
| Gm16036 | Gm15409 | Gm16043 | Gm43154 | 2210010C04Rik |
| Olfr718-ps1 | 5930430L01Rik | Gm35822 | Gm13848 | Trbv2 |
| Rbakdn | Gm15408 | AA545190 | Gm43293 | 5830405F06Rik |
| 4930448H16Rik | 5730422E09Rik | Gm44129 | Gm13849 | Trbv6 |
| BC030343 | Gm42787 | 1700016P04Rik | 1700012A03Rik | Trbv7 |
| E130309D02Rik | Gm42788 | Bmt2 | Gm13850 | Trbv8 |
| 0610040B10Rik | Gm43332 | 2610001J05Rik | Plxna4os3 | Trbv9 |
| Gm43397 | 2310047D07Rik | 1110019D14Rik | Plxna4os2 | Trbv10 |
| Gm20635 | Gm29264 | Gm36503 | Plxna4os1 | Trbv11 |
| Gm42504 | Gm19719 | Gm36669 | BC049739 | Trbv12-3 |
| D130017N08Rik | BC028471 | Gm15473 | Gm14540 | Trbv18 |
| Gm17135 | Gm43151 | Gm4876 | Gm13853 | Trbv21 |
| Gm42421 | Gm43150 | D830026I12Rik | Gm13856 | Trbv22 |
| 2900089D17Rik | Gm15997 | Gm15581 | Gm13855 | Trbv24 |
| Gm4871 | Wdr95 | Gm29591 | Gm14546 | Trbv25 |
| Gm43562 | Gm20005 | Gm26738 | Akr1b3 | Trbv27 |

| | | | | |
|---|---|---|---|---|
| Trbv28 | Olfr434 | Gm4872 | Gm43284 | Igkv4-77 |
| Gm5771 | Olfr47 | Gm16499 | Igkv9-129 | Igkv13-76 |
| Gm10334 | Gm44731 | Gm16499.1 | Igkv9-128 | Igkv4-75 |
| Trbd1 | 9430018G01Rik | 9530036M11Rik | Igkv14-126-1 | Igkv13-74-1 |
| Trbj1-1 | Gm35216 | Gm44026 | Igkv9-120 | Igkv13-73-1 |
| Trbj1-2 | Gm26625 | Gm28402 | Igkv9-119 | Igkv4-73 |
| Trbj1-3 | Zfp956 | Gars | Igkv14-118-2 | Igkv4-72 |
| Trbj1-4 | Gm16630 | Gm34358 | Igkv14-118-1 | Igkv13-71-1 |
| Trbj1-5 | Gm44696 | Gm44073 | Gm43739 | Igkv4-71 |
| Trbj1-6 | 1700026J14Rik | 6430584L05Rik | Igkv11-118 | Igkv4-68 |
| Trbj1-7 | Gm5111 | Gm3279 | Igkv2-116 | Igkv12-67 |
| Trbd2 | Gm45021 | Ccdc129 | Igkv1-115 | Rprl1 |
| Trbj2-1 | Gm44965 | Gm8239 | Igkv11-114 | Igkv12-66 |
| Trbj2-2 | Gm38804 | Vmn1r13 | Igkv2-113 | Igkv4-65 |
| Trbj2-3 | AI854703 | Vmn1r20 | Igkv1-108 | Igkv13-64 |
| Trbj2-4 | Gimap9 | 4930533I22Rik | Igkv2-107 | Igkv4-63 |
| Trbj2-6 | Gm28053 | Vmn1r23 | Igkv11-106 | Igkv13-62-1 |
| Trbj2-7 | Gm44226 | Gm43994 | Igkv2-105 | Igkv4-62 |
| Olfr459 | 4833403J16Rik | Vmn1r27 | Igkv15-102 | Igkv13-61-1 |
| Sval1 | Gm30781 | Gm35077 | Igkv20-101-2 | Igkv4-61 |
| Sval3 | Gm7932 | Gm45060 | Igkv15-101-1 | Igkv4-59 |
| Sva | 1600015I10Rik | BB365896 | Igkv15-101 | Igkv4-60 |
| Gm44284 | Doxl2 | Gm5570 | Igkv1-99 | Igkv13-57-2 |
| 2010310C07Rik | 4930563H07Rik | Gm44410 | Igkv15-97 | Igkv4-57-1 |
| Tas2r143 | Svs1 | Gm35386 | Igkv2-95-2 | Igkv13-57-1 |
| Olfr458 | Gm3455 | Gm43893 | Igkv2-95-1 | Igkv13-56-1 |
| Olfr457 | Mpp6 | Gm43894 | Igkv10-95 | Igkv4-56 |
| Olfr456 | 5430402O13Rik | Gm44071 | Igkv2-93-1 | Igkv13-55-1 |
| Olfr455 | 4921507P07Rik | Gm44072 | Gm42925 | Igkv13-54-1 |
| Tcaf3 | C530044C16Rik | C130060K24Rik | Igkv4-91 | Igkv4-54 |
| Olfr453 | Gm44109 | Gm16838 | Gm42924 | Igkv4-53 |
| Olfr38 | G930045G22Rik | 4930544G11Rik | Igkv1-88 | Gm42606 |
| Olfr452 | Gm32479 | 2610300M13Rik | Gm42539 | Igkv4-51 |
| Olfr450 | Gm26677 | Vmn1r33 | Gm42543 | Gm42958 |
| Olfr449 | Gm6559 | Vmn1r34 | Gm42542 | Igkv12-49 |
| Olfr448 | 0610033M10Rik | Vmn1r35 | Igkv13-87 | Igkv12-47 |
| Olfr447 | Gm38811 | Vmn1r36 | Igkv4-86 | Igkv12-42 |
| Olfr446 | Gm44077 | Vmn1r37 | Igkv4-83 | Igkv5-40-1 |
| Olfr444 | Halr1 | Vmn1r38 | Igkv13-82 | Igkv12-40 |
| Gm17472 | Gm28308 | Vmn1r39 | Igkv4-81 | Gm43220 |
| Gm20730 | Gm15050 | Gm36816 | Igkv13-80-1 | Gm43586 |
| Olfr441 | 5730596B20Rik | Gm38825 | Igkv4-80 | Igkv8-31 |
| Olfr237-ps1 | Gm29430 | Gm42889 | Igkv4-79 | Gm43218 |
| Olfr437 | Evx1os | Igkv1-136 | Igkv13-78-1 | Igkv8-22 |
| Olfr13 | 1700094M24Rik | Igkv14-134-1 | Igkv4-78 | Gm42667 |
| Olfr435 | Gm44028 | Igkv17-134 | Gm43583 | Gm10360 |

| | | | | |
|---|---|---|---|---|
| Gm42720 | Gm38843 | Gm15612 | Gm15631 | D330020A13Rik |
| Igkv3-12-1 | Gm32591 | 4933427D06Rik | Gm34933 | Gm20692 |
| Igkv3-11 | 2310069B03Rik | Gm1965 | 4933431M02Rik | 1700072O05Rik |
| Igkv3-8 | Gm45145 | Gm6507 | Gm44040 | Gm43916 |
| Igkv3-6 | Gm42688 | Gm839 | 0610040F04Rik | Gm43915 |
| Gm30211 | Ccdc142os | Gm44204 | Gm20387 | Gm15856 |
| Igkj1 | Gm44287 | 4930512J16Rik | 1700054K19Rik | Gm44148 |
| Igkj2 | Gm15624 | Gm26811 | 5031434C07Rik | Gm7298 |
| Igkj3 | Gm21284 | Gm44117 | Gm26799 | 1700063H04Rik |
| Igkj4 | Vax2os | Gm44421 | Gm44081 | Gm26826 |
| Igkj5 | 1700124L16Rik | Gm45901 | Gm16161 | Vmn2r19 |
| Tex37 | Gm33024 | Gm15756 | Gm15492 | Vmn2r20 |
| Gm44427 | Gm15475 | Gm44123 | Gm44199 | Vmn2r21 |
| Gm1070 | Gm44127 | Gm44278 | 1700015O11Rik | Vmn2r22 |
| Gm44110 | Gm10445 | Gm14573 | Gm43964 | Vmn2r23 |
| Gm44174 | 4930504D19Rik | 4930471M09Rik | Gm26982 | Vmn2r24 |
| Gm44175 | Gm44012 | 1810044D09Rik | Gm44053 | Vmn2r25 |
| Gm44172 | Gm5878 | Gm45218 | Gm15082 | Vmn2r26 |
| 1700040L08Rik | Gm19265 | 4930466I24Rik | Gm15083 | Vmn2r27 |
| Gm26628 | 4930553P18Rik | A730049H05Rik | Gm17482 | Gm43971 |
| 9130221F21Rik | Gm44089 | Gm15737 | Gm44079 | 1700013D24Rik |
| 4933431G14Rik | Gm44941 | 4930511E03Rik | Gm36355 | 1700027F06Rik |
| Gm38832 | Gm44790 | Kbtbd8os | Gm44206 | Gm44006 |
| Gm45051 | A430078I02Rik | Gm10234 | H1foo | Gm44096 |
| Gm15401 | 2310040G24Rik | Fam19a1 | Gm20404 | Gm16567 |
| Gm15402 | Gm43936 | 1700123L14Rik | 8-Mar | Gm15884 |
| Gm26640 | Gm44386 | Fam19a4 | Olfr211 | Gm20531 |
| Gm20560 | 1600020E01Rik | Gm32592 | Olfr212 | Gm45234 |
| 1700065L07Rik | Gm44214 | Gm765 | Olfr213 | A230083G16Rik |
| 4931417E11Rik | Gm44036 | Gm44196 | Gm5580 | 4930557K07Rik |
| Gm31747 | D6Ertd527e | 4930595L18Rik | Olfr214 | Gm28809 |
| Gm44113 | Gm44418 | Gm20696 | Olfr215 | Gm10415 |
| Gm9008 | Gm34312 | Gm26748 | 8430408G22Rik | Gm29009 |
| Gm44166 | Gm44097 | Gm43067 | D030044L04Rik | Gm45869 |
| Gm29005 | 1810020O05Rik | Gm44442 | Gm4640 | Gm26673 |
| Gm20362 | Gm44210 | Gm33201 | 6820426E19Rik | 1700018A23Rik |
| Gm44271 | Gm43904 | Gm44107 | Gm9946 | Platr31 |
| Gm20594 | Gm45140 | 1700049E22Rik | Gm45083 | Gm43126 |
| Gm4409 | Gm26636 | Gm43950 | 1700030F04Rik | 9330179D12Rik |
| Gm38839 | H1fx | Gm43948 | Gm7292 | Gm43635 |
| 1700009C05Rik | Gm5577 | Gm26911 | Gm29509 | Gm38404 |
| Gm20383 | Gm38708 | Gm44169 | 1700069P05Rik | Gm43634 |
| Gm44202 | 1700031F10Rik | Gm9871 | Gm30557 | Gm42738 |
| Gm15864 | Gm44264 | Gm44176 | Gm30498 | Gm38901 |
| Gm44145 | Gm44170 | 4930587E11Rik | Gm44317 | Gm15870 |
| Gm17034 | Gm26588 | Gm19757 | 4930540M05Rik | 9330102E08Rik |

| | | | | |
|---|---|---|---|---|
| Gm10010 | 4933406J09Rik | Casc1 | Gm15873 | Vmn2r34 |
| Gm26770 | Gm44448 | Gm15543 | Gm44877 | Vmn2r35 |
| Gm44596 | Gm38910 | Gm15706 | Fam71e2 | Vmn2r36 |
| Gm15862 | Gm36328 | 2010013B24Rik | Gm44983 | Vmn2r41 |
| Gm10069 | 2810454H06Rik | Gm15499 | Gm44973 | Vmn2r42 |
| Pzp | Gm44238 | Gm15704 | Zfp865.1 | Vmn2r43 |
| A2ml1 | Lockd | Sspnos | Gm15510 | Vmn2r44 |
| Gm44009 | Gm19434 | Gm32914 | Vmn1r55 | Vmn2r45 |
| Gm44511 | Gm36640 | 4930479D17Rik | Vmn1r56 | Vmn2r40 |
| Gm44066 | Gm44275 | Arntl2 | Vmn1r57 | Vmn2r39 |
| Gm26656 | 4930425L21Rik | Gm44085 | Gm5065 | Vmn2r38 |
| Gm44067 | Gm14329 | E330012B07Rik | Vmn1r58 | Vmn2r37 |
| BC035044 | Gm14330 | Gm15767 | Vmn1r59 | Vmn2r46 |
| Gm15987 | Gm26653 | Gm6288 | Vmn2r28 | Vmn2r47 |
| Gm44000 | Gm8994 | 1700049E15Rik | Vmn1r60 | Gm45351 |
| 2310001H17Rik | E330021D16Rik | Gm15762 | Vmn1r61 | Vmn2r48 |
| Gm47861 | Hist4h4 | Far2os2 | Vmn1r62 | Vmn2r49 |
| 4922502D21Rik | H2afj | Far2os1 | Vmn1r63 | Vmn2r50 |
| Klre1 | A830011K09Rik | Gm17216 | Vmn1r64 | Vmn2r51 |
| Klri1 | Gm30332 | Gm44020 | Vmn1r65 | Vmn2r52 |
| Klri2 | B230110G15Rik | Gm6313 | 4933402C05Rik | Vmn1r72 |
| Gm156 | 4922502N22Rik | 4930528G23Rik | Gm44733 | Vmn1r73 |
| Klra17 | 4930404I20Rik | Gm44115 | Epp13 | Vmn1r74 |
| Gm17631 | Gm11077 | Gm43913 | Gm45069 | Vmn1r75 |
| Klra5 | Gm30524 | Fam60a | Gm38944 | Vmn1r76 |
| Klra6 | Gm28523 | 3010003L21Rik | Gm3854 | Vmn1r77 |
| Klra4 | Gm5724 | Gm26540 | Gm20715 | Vmn1r78 |
| Klra8 | Gm30784 | Gm10203 | Olfr1344 | Vmn1r79 |
| Klra9 | Gm6614 | 1700003I16Rik | Olfr1336 | Vmn1r80 |
| Klra7 | Gm20400 | 2810474O19Rik | Olfr1346 | Gm4224 |
| Klra10 | 5330439B14Rik | Gm21814 | Olfr5 | Vmn1r81 |
| Klra3 | A930014E10Rik | Speer9-ps1 | Olfr1347 | Vmn1r82 |
| Klra1 | 4930434O05Rik | AU018091 | Olfr1348 | Vmn1r83 |
| Klra2 | Gm31108 | Gm44257 | Olfr1349 | Vmn1r84 |
| 5430401F13Rik | 1700126G02Rik | Gm44151 | Olfr1350 | Vmn2r53 |
| Gm6619 | D6Ertd474e | 3300002P13Rik | Zim1 | Vmn2r54 |
| Gm43974 | 4930579D09Rik | Gm35037 | Zim3 | Vmn2r55 |
| Prb1 | 1700060C16Rik | Gm44920 | Zfp418 | Vmn2r56 |
| Prpmp5 | Sox5os1 | Gm15929 | Gm45844 | Gm45593 |
| Gm8882 | Sox5os2 | Gm15448 | Vmn2r29 | Gm31152 |
| A630073D07Rik | Sox5os3 | Gm14548 | Gm45631 | 1810019N24Rik |
| Tas2r122 | Sox5os4 | Gm44581 | Vmn2r30 | Gm29638 |
| 5530400C23Rik | Sox5os5 | 1700095K22Rik | Gm45783 | Vmn1r85 |
| Kap | Gm26666 | 9430041J12Rik | Vmn2r31 | Gm45436 |
| Gm17088 | Gm15687 | D030047H15Rik | Vmn2r32 | Vmn1r86 |
| Gm17089 | Lrmp | Gm15494 | Vmn2r33 | Vmn1r87 |

| | | | | |
|---|---|---|---|---|
| Vmn1r88 | Gm16251 | Gm4177 | Vmn1r174 | Gm38979 |
| Gm38947 | Vmn1r91 | Vmn1r135 | Vmn1r175 | Gm26604 |
| Vmn1r89 | Gm16442 | Gm5725 | Vmn1r176 | Gm16282 |
| Bsph2 | Vmn1r93 | Gm5891 | Vmn1r177 | 1110035H17Rik |
| Obox8 | Vmn1r94 | Vmn1r137 | Vmn1r178 | Gm44632 |
| Vmn1r90 | Vmn1r95 | Vmn1r138 | Vmn1r179 | Gm29326 |
| Obox7 | Gm6164 | Gm6176 | Vmn1r180 | A330087D11Rik |
| Obox2 | Gm4498 | Vmn1r139 | Vmn1r181 | Zfp940 |
| Obox1 | Gm5890 | Gm16451 | V1rd19 | Gm26554 |
| Obox3 | Gm4133 | Gm10666 | Gm36694 | Zfp568 |
| Obox5 | Gm5728 | Vmn1r142 | Vmn1r183 | Gm26920 |
| Obox4-ps35 | Vmn1r100 | Vmn1r143 | Gm44793 | Gm5113 |
| Obox6 | Vmn1r101 | Gm8653 | Tescl | Gm26810 |
| Crxos | Gm10670 | Gm8453 | Irgc1 | Thap8 |
| Gm38948 | Vmn1r103 | Gm4187 | Gm36159 | E130208F15Rik |
| Gm45509 | Vmn1r104 | Gm8660 | Gm26550 | 4930479H17Rik |
| Gm45510 | Gm5726 | Vmn1r148 | Gm4598 | Nphs1os |
| Gm29443 | Gm4141 | Vmn1r149 | BC049730 | Gm49396 |
| Gm29442 | Gm4513 | Gm10665 | Gm4763 | Gm26610 |
| 9330104G04Rik | Vmn1r107 | Vmn1r151 | Gm9844 | 2200002J24Rik |
| Pnmal2 | Vmn1r111 | Vmn1r152 | Gm4881 | Tmem147os |
| Pnmal1 | Gm10668 | Gm8677 | Gm44970 | Gm26935 |
| Gm42372 | Vmn1r112 | Gm4201 | 4933430L12Rik | Gm29439 |
| Gm32772 | Vmn1r113 | Gm4565 | 9130221H12Rik | D7Ertd128e |
| Gm32846 | Vmn1r114 | Vmn1r155 | 4732471J01Rik | Gm44662 |
| Gm44613 | Vmn1r115 | Gm8693 | Gm29918 | Gm17077 |
| Gm26821 | Vmn1r116 | Gm4567 | Gm38569 | Gm4673 |
| Gm45058 | Vmn1r117 | Vmn1r157 | Gm26707 | G630030J09Rik |
| Mypopos | Vmn1r118 | Vmn1r158 | Gm30146 | Gm10640 |
| Gm4969 | Vmn1r119 | Vmn1r159 | BC026762 | Scgb2b2 |
| 1700058P15Rik | Vmn1r120 | Vmn1r160 | Gm42375 | Scgb1b2 |
| Gm26802 | Vmn1r121 | Gm4214 | Gm15883 | Scgb2b3 |
| D830036C21Rik | Vmn1r122 | Gm4216 | Gm21983 | Scgb1b3 |
| Gm44698 | Vmn1r123 | Vmn1r163 | BC024978 | Scgb2b7 |
| Cd3eap | Gm5157 | Gm8720 | Gm15567 | Scgb1b7 |
| Gm26852 | Vmn1r124 | Gm10662 | Gm20479 | Scgb1b10 |
| A930016O22Rik | Vmn1r125 | Vmn1r165 | Gm15541 | Scgb2b11 |
| Gm26890 | Vmn1r126 | Vmn1r166 | Gm26891 | Scgb1b11 |
| Gm44805 | Vmn1r127 | Gm6902 | Gm44744 | Scgb2b12 |
| Gm20512 | Vmn1r128 | Vmn1r167 | 9530053A07Rik | Scgb1b12 |
| Gm26600 | Vmn1r129 | Vmn1r168 | Gm44709 | Scgb1b29 |
| Gm16175 | Gm6882 | Vmn1r169 | Gm44710 | Scgb2b15 |
| Gm16174 | Vmn1r130 | Vmn1r170 | Gm44618 | Scgb1b15 |
| Gm44658 | Vmn1r131 | Vmn1r171 | Gm44702 | Scgb2b17 |
| Gm19345 | Vmn1r132 | Vmn1r172 | Gm10648 | Scgb1b17 |
| Gm44659 | Gm4175 | Vmn1r173 | Gm44700 | Scgb2b18 |

| | | | | |
|---|---|---|---|---|
| Scgb1b18 | Gm37494 | Gm36864 | Gm44644 | Gm35325 |
| Scgb2b19 | Gm26526 | Gm15517 | Gm44643 | Gm29327 |
| Scgb1b19 | Gm45839 | Gm44780 | A230006K03Rik | Gm44946 |
| Scgb2b20 | 4930435C17Rik | 5430431A17Rik | B230209E15Rik | Gm44948 |
| Scgb1b20 | 4930558N11Rik | Fam71e1 | A230057D06Rik | Gm28258 |
| Scgb2b21 | Gm44992 | Gm15396 | Gm9801 | Gm44532 |
| Gm12763 | 4933404I11Rik | Gm44832 | Gm32061 | Gm35842 |
| Scgb2b24 | Gm28807 | 2310016G11Rik | A26c2 | Gm44813 |
| Scgb1b24 | 4933402C06Rik | 6530437J22Rik | 4930554H23Rik | Gm44812 |
| Scgb2b26 | A230077H06Rik | Gm15545 | Gm44718 | Gm44808 |
| Scgb2b27 | Gm28307 | Gm45669 | E030018B13Rik | Gm42397 |
| Scgb1b27 | Gm38997 | Gm45552 | Gm45052 | Gm36011 |
| Scgb1b30 | 1700110I07Rik | Gm45713 | Gm32633 | Gm39033 |
| Gm26762 | Gm44839 | Ccdc155 | Gm45054 | 6030442E23Rik |
| Gm12762 | Gm44748 | Mtag2 | Gm20670 | Gm29328 |
| Gm12758 | Gm34811 | Gm45619 | Gm20457 | Gm10295 |
| 4931406P16Rik | Gm17102 | Gm45808 | Fam189a1 | Gm29295 |
| Gm6096 | Gm20449 | Gm45564 | 1810049I09Rik | Gm44689 |
| Gm12764 | Vmn2r57 | 0610005C13Rik | BC046251 | 4930441H08Rik |
| Gm12766 | 2610021A01Rik | Gm31479 | Tarsl2 | Gm36633 |
| Gm12784 | Zfp788 | Gm45437 | 1810008I18Rik | 1700011C11Rik |
| Gm44837 | Vmn2r58 | A030001D20Rik | Gm45213 | Gm36696 |
| Gm12781 | Vmn2r59 | Gm31597 | Gm45081 | Gm28745 |
| Gm12756 | Vmn2r60 | Gm45441 | Gm26827 | Gm28744 |
| Gm45095 | Vmn2r61 | Gm45442 | Gm44752 | Gm44725 |
| Gm45096 | 4933421I07Rik | Ccdc114 | Gm10974 | Gm6567 |
| Gm35611 | Zfp141 | Gm45486 | Gm44753 | 4933435G04Rik |
| Gm45091 | Gm21028 | 1700025L06Rik | Gm33570 | Gm28746 |
| Gm35665 | Zfp975 | Gm15700 | 1700112J16Rik | Gm20083 |
| B230322F03Rik | Vmn2r62 | C86187 | Gm33926 | Gm30075 |
| Gm26790 | Gm2381 | Gm9999 | 4833412C05Rik | 4930429H19Rik |
| E130304I02Rik | Vmn2r63 | Gm45474 | Gm34350 | A730056A06Rik |
| Gm28514 | Gm9268 | Gm14377 | Gm39027 | Gm44686 |
| Gm38991 | Gm17768 | Gm44969 | Gm16157 | Gm44734 |
| Gm29129 | EU599041 | Gm45086 | Gm16158 | Gm44738 |
| Gm28078 | Zfp715 | Gm45282 | Gm44889 | Gm44737 |
| Gm28076 | Gm45233 | Gm32849 | 4933436H12Rik | Gm44736 |
| Gm28075 | 4931406B18Rik | 9130015G15Rik | Gm34549 | Gm44739 |
| Gm28077 | Gm2511 | Gm26856 | Gm44886 | Gm10619 |
| Gm45768 | Gm44818 | Nell1os | Gm44692 | Gm21269 |
| 4930505M18Rik | Zfp719 | 4933405O20Rik | Gm34664 | Gm45004 |
| Gm30684 | Zfp819 | Gm29296 | Gm44691 | Gm7580 |
| Gm30771 | Gm36546 | 1700015G11Rik | Gm34838 | Gm30459 |
| D530033B14Rik | Gm44707 | Gm44506 | Gm38584 | 0610006L08Rik |
| Gm44767 | Gm44756 | Gm9962 | 4930402F11Rik | 4930533N22Rik |
| Gm29087 | 1700028J19Rik | Gm38393 | Gm29683 | AU020206 |

| | | | | |
|---|---|---|---|---|
| 4921513I08Rik | Gm30873 | Gm45159 | Olfr547 | Olfr601 |
| Gm9885 | Gm44933 | 4930567K12Rik | Olfr548-ps1 | Olfr603 |
| 5330411O13Rik | Gm45175 | Gm26944 | Olfr549 | Olfr605 |
| 1700011D18Rik | Gm2115 | Gm26981 | Olfr550 | Olfr606 |
| Gm44730 | 2610206C17Rik | Gm9934 | Olfr551 | Olfr607 |
| Gm26633 | Olfr291 | Gm44947 | Olfr552 | Olfr608 |
| Gm35040 | Olfr290 | Gm45037 | Olfr553 | Olfr609 |
| Gm45168 | Olfr290.1 | Gm32647 | Olfr554 | Olfr610 |
| Gm45169 | Vmn2r65 | Gm44601 | Olfr555 | Olfr611 |
| 9330171B17Rik | Vmn2r66 | Gm15412 | Olfr556 | Olfr612 |
| Gm44706 | Vmn2r67 | Gm15414 | Olfr557 | Olfr613 |
| Gm26646 | Vmn2r68 | Gm15413 | Olfr558 | Olfr615 |
| Gm21057 | Vmn2r69 | Gm15416 | Olfr33 | Olfr616 |
| Gm44949 | Vmn2r70 | Gm44633 | Olfr559 | Olfr617 |
| Gm45202 | Vmn2r71 | Rsf1os1 | Olfr78 | Olfr618 |
| 6330403N20Rik | Vmn2r72 | Rsf1os2 | Olfr560 | Olfr619 |
| Gm44851 | Vmn2r73 | Gm44507 | Olfr561 | Olfr620 |
| Gm15880 | Vmn2r74 | Gucy2d | Olfr564 | Olfr622 |
| Gm44649 | Vmn2r75 | A630091E08Rik | Olfr566 | Olfr623 |
| Gm44899 | Vmn2r76 | Gm19656 | Olfr568 | Olfr624 |
| Gm44968 | Olfr310 | Gm44934 | Olfr569 | Olfr625-ps1 |
| BC048679 | Olfr309 | Gm45188 | Olfr570 | Olfr243 |
| 2900076A07Rik | Olfr308 | Gm45187 | Olfr571 | Olfr628 |
| Gm44570 | Olfr307 | Gm45012 | Olfr572 | Olfr629 |
| Fam103a1 | Olfr305 | Gm26705 | Olfr573-ps1 | Olfr630 |
| Gm45016 | Olfr304 | Gm10605 | Olfr574 | Olfr69 |
| Gm10160 | Olfr303 | Gm15635 | Olfr575 | Olfr68 |
| Gm45014 | Olfr301 | Gm34280 | Olfr576 | Olfr67 |
| Gm20744 | Olfr299 | Olfr520 | Olfr577 | Olfr66 |
| Gm45056 | Olfr298 | Olfr521 | Olfr578 | Olfr64 |
| Gm32850 | Olfr297 | F730035P03Rik | Olfr582 | Olfr65 |
| Gm45090 | Olfr295 | Gm38405 | Olfr583 | Olfr631 |
| 4933406J10Rik | Olfr294 | Gm34821 | Olfr584 | Olfr632 |
| 1700010L04Rik | Olfr293 | Gm10603 | Olfr585 | Olfr633 |
| Gm36584 | Olfr292 | Gm39059 | Olfr586 | Olfr635 |
| 4933430H16Rik | Vmn2r77 | Gm35082 | Olfr588-ps1 | Olfr638 |
| A530021J07Rik | Vmn2r78 | Gm20476 | Olfr589 | Olfr643 |
| 5930435M05Rik | Vmn2r79 | Gm45620 | Olfr591 | Olfr639 |
| Gm44724 | Gm26522 | Gm45837 | Olfr592 | Olfr640 |
| Gm44991 | Gm44751 | Gm35363 | Olfr593 | Olfr641 |
| Gm10610 | Gm45144 | Gm10602 | Olfr594 | Olfr642 |
| Gm16638 | Gm44995 | Xndc1 | Olfr596 | 4930516K23Rik |
| 2310044K18Rik | AI314278 | Xntrpc | Olfr597 | Olfr644 |
| Gm44993 | Prss23os | Olfr543 | Olfr598 | Olfr645 |
| Gm39043 | Gm45064 | Olfr544 | Olfr599 | Olfr646 |
| Gm44827 | A230065N10Rik | Olfr545 | Olfr600 | Ubqln5 |

| | | | | |
|---|---|---|---|---|
| Olfr648 | Gm20663 | Olfr482 | Gm33586 | Gm45847 |
| Olfr649 | Gvin1 | Olfr483 | 4930543E12Rik | Gm27040 |
| Gm47248 | Olfr693 | Olfr484 | Arntl | Gm15339 |
| Trim12a | Olfr694 | Olfr485 | Gm45355 | 3100003L05Rik |
| Gm15133 | Olfr695 | Olfr486 | Far1os | Gm45092 |
| Olfr651 | Olfr697 | Olfr487 | Gm5600 | 4933440M02Rik |
| Olfr652 | Olfr698 | Olfr488 | Gm29507 | 4930571K23Rik |
| Olfr653 | Olfr699 | Olfr490 | Gm45615 | Gm30717 |
| Olfr654 | Olfr700 | Olfr491 | 4933406I18Rik | Gm44876 |
| Olfr655 | 4930458B22Rik | Olfr492 | Gm45017 | D430042O09Rik |
| Olfr656 | Olfr701 | Olfr493 | Gm45020 | 1700123J17Rik |
| Olfr657 | Olfr702 | Olfr494 | A730082K24Rik | Gm30928 |
| Olfr658 | Olfr703 | Olfr495 | Gm34225 | 2510046G10Rik |
| Olfr659 | Olfr704 | Olfr497 | Gm45025 | 4930448A20Rik |
| Olfr661 | Olfr705 | Olfr498 | Sox6os | Fam57b |
| Gm45556 | Olfr706 | Olfr502 | Gm4353 | Aldoa.1 |
| Olfr665 | Olfr707 | Olfr503 | Gm44777 | Gm44854 |
| Olfr666 | Olfr1532-ps1 | Olfr504 | Gm45151 | Pagr1b |
| Gm45272 | Olfr709-ps1 | Olfr506 | 4732496C06Rik | Gm42742 |
| Olfr667 | Olfr710 | Olfr507 | Gm39075 | Gm31749 |
| Olfr668 | Olfr6 | Olfr508 | Gm45155 | Gm44939 |
| Olfr669 | Olfr711 | Olfr509 | Gm35309 | 1-Sep |
| Olfr670 | Olfr2 | Olfr510 | B230311B06Rik | Gm31897 |
| Olfr671 | Olfr713 | Olfr512 | 9030624J02Rik | 9130019O22Rik |
| Olfr672 | Olfr714 | Olfr513 | Gm44763 | E430018J23Rik |
| Olfr675 | Olfr17 | Olfr514 | Gm44588 | Gm45184 |
| Olfr676 | Olfr715b | Olfr516 | Gm39078 | 1700008J07Rik |
| Olfr677 | Gm45868 | Olfr517 | Gm19950 | Gm42715 |
| Olfr678 | Olfr715 | Olfr518 | Gm45792 | Gm28198 |
| Gm45557 | Olfr716 | Olfr519 | Gm44866 | Ccdc189 |
| Olfr679 | Platr28 | Gm26599 | Tmem159 | Gm44759 |
| Olfr680-ps1 | Gm44773 | Gm44781 | 4930505K13Rik | 1700120K04Rik |
| Gm45558 | 5330417H12Rik | Gm45024 | Abca14 | Gm39090 |
| Olfr681 | Gm45040 | 1700095J03Rik | Abca16 | Gm49388 |
| Olfr683 | Olfr467 | St5 | Gm32916 | Gm21974 |
| Olfr684 | Olfr469 | Gm44864 | 4930588G17Rik | Gm26690 |
| Olfr685 | Olfr470 | 1600010M07Rik | Gm5737 | Gm15533 |
| Olfr686 | Olfr472 | Gm45515 | Gm15774 | Gm49368 |
| Olfr688 | Olfr473 | Gm28863 | Gm45079 | Gm39091 |
| Olfr689 | Olfr474 | Gm21123 | Gm36736 | Gm44850 |
| Olfr690 | Olfr476 | Mrvi1 | 4933432K03Rik | 9130023H24Rik |
| Olfr691 | Olfr477 | Gm16336 | 1700025J12Rik | BC017158 |
| Olfr692 | Olfr478 | 5430402P08Rik | 1700069B07Rik | Gm40457 |
| Fam160a2 | Olfr479 | Gm49324 | Gm44987 | Gm45121 |
| Gm45799 | Olfr480 | Micalcl | Gm15489 | Gm44674 |
| Gm4070 | Olfr481 | Parvaos | Gm45846 | Gm44672 |

| | | | | |
|---|---|---|---|---|
| Gm32884 | Gm45686 | Gm45889 | Gm48759 | Gm32926 |
| Gm33027 | 4930543N07Rik | Gm49369 | Gm46210 | Gm4922 |
| Gm39094 | Gm30808 | Gm33148 | Gm28905 | 4930444F02Rik |
| Gm44778 | Gm45507 | Gm49394 | Gm48278 | Gm48483 |
| Gm4265 | Gm45613 | Gm6471 | Gm48281 | 1700124M09Rik |
| Gm44647 | Gm32486 | Gm39117 | Gm48279 | Gm48545 |
| Etos1 | Gm2044 | Tspan32os | Gm31904 | Gm20139 |
| Gm5602 | 6430531B16Rik | Gm28821 | Gm48324 | Gm33056 |
| 1700029B22Rik | Msx3 | 4933417O13Rik | Gm10944 | Gm33104 |
| 5430419D17Rik | Olfr522 | Gm44732 | 4930567K20Rik | 9230106D20Rik |
| Gm44623 | Olfr523 | Cars | Gm16577 | AC153891.1 |
| Gm44546 | Olfr524 | Gm15579 | Gm48680 | 4930405J17Rik |
| Gm15677 | Cd163l1 | Tnfrsf26 | Gm48697 | Gm48249 |
| Gm10584 | 5830411N06Rik | E230032D23Rik | Gm48721 | 4933406P04Rik |
| Gm44892 | Olfr525 | Tnfrsf22 | Gm48722 | Gm48651 |
| Gm34908 | Olfr527 | Tnfrsf23 | Gm48723 | Gm20149 |
| Gm44893 | Olfr60 | Gm498 | Gm48724 | Gm48189 |
| Gm35147 | Olfr530 | Gm14372 | B230208H11Rik | Gm26577 |
| Gm16764 | Olfr531 | Faddos | Gm47691 | Gm40608 |
| Gm15582 | Olfr532 | Gm44930 | Gm48773 | Gm33619 |
| Gm45502 | Olfr533 | Gm44929 | Gm32172 | Gm33728 |
| Gm15718 | Olfr535 | Gm34964 | Gm32105 | 4930455C13Rik |
| 4930483O08Rik | Olfr536 | Gm26793 | Gm26835 | 1700021A07Rik |
| Gm45672 | Olfr538 | Gm39119 | Gm28289 | 1700020N01Rik |
| Gm35625 | Olfr46 | Bc1 | Gm48843 | Gm5420 |
| Gm45671 | Olfr61 | Oraov1 | Gm48845 | Gm29323 |
| Gm28747 | Olfr53 | Gm45181 | Gm48844 | Platr6 |
| Gm45670 | Olfr539 | 1810010D01Rik | Gm40603 | 4930444G20Rik |
| Gm4593 | Olfr45 | 9230019H11Rik | Gm48854 | E030030I06Rik |
| Fam196a | Olfr541 | B020014A21Rik | Gm46212 | Gm26740 |
| 1700120G07Rik | Gm45785 | H60c | Gm16234 | Gm26581 |
| Gm45339 | C330022C24Rik | Gm26752 | B230364G03Rik | Gm10825 |
| Gm36356 | Odf3 | Gm30570 | Gm47698 | Gm10824 |
| 5830432E09Rik | Gm45717 | Gm21781 | Gm47710 | Gm47843 |
| Gm36431 | Gm17387 | Gm10097 | Gm47712 | 4930520K02Rik |
| Gm45241 | Gm16982 | Gm10945 | Gm40604 | Gm40614 |
| 6330420H09Rik | Gm45618 | B430219N15Rik | Gm32283 | Gm15271 |
| Gm36737 | Gm39115 | Gm48614 | C330004P14Rik | Ctgf |
| C030029H02Rik | Gm45337 | Gm30906 | Gm47729 | Gm47715 |
| C230079O03Rik | Gm29735 | Gm48654 | Gm47760 | Gm40617 |
| Gm36849 | Gm10153 | Gm26674 | Gm47761 | Gm29571 |
| Gm6249 | Krtap5-3 | Gm48727 | Gm47769 | Gm36908 |
| Gm45278 | Gm7579 | Gm48728 | Gm47770 | Gm36543 |
| Gm10578 | Gm40460 | Gm48748 | Gm47771 | 4930401C15Rik |
| Gm45397 | Gm2431 | Gm48749 | Gm20655 | Gm47939 |
| Gm45682 | Gm4559 | Gm48757 | Gm10827 | Gm9767 |

| | | | | |
|---|---|---|---|---|
| Gm30034 | A930033M14Rik | Lilr4b | 4933428P19Rik | Gm48022 |
| Gm48893 | Cdk19os | Gm40645 | Gm16135 | Gm40689 |
| Gm48084 | Gm48057 | Lilrb4a | Gm16143 | Gm48023 |
| 4930579H20Rik | Gm32284 | Gm46224 | Gm48072 | Gm48024 |
| Gm30165 | AC166165.1 | 1700042O05Rik | Gm26789 | 4930533K18Rik |
| Gm30228 | Gm17196 | 4933411G06Rik | Gm16145 | Gm48036 |
| Soga3 | BC048559 | Gm47613 | Gm7075 | Gm15647 |
| Gm49353 | Gm15200 | Gm47614 | Gm10118 | C730027H18Rik |
| Gm46218 | Gm15199 | Gm40646 | Gm28881 | AC079680.2 |
| Gm40621 | Gm26860 | Gm26741 | Gm30853 | AC079680.1 |
| Gm48159 | Gm34006 | Gm36065 | 1700023F02Rik | Gm34376 |
| Gm48215 | Gm48671 | Gm40649 | 4930407I19Rik | Gm20609 |
| Gm20300 | Gm47815 | Gm36173 | Gm28139 | 4930551I15Rik |
| Gm30676 | Gm29246 | Gm19395 | Gm31181 | Gm47028 |
| Gm48094 | 9030612E09Rik | Gm36229 | Gm47895 | AC121788.1 |
| Gm48093 | Gm47889 | AC155941.1 | Gm47896 | Gm34609 |
| Gm47823 | Gm9803 | Gm20597 | Gm47897 | Gm34776 |
| D830005E20Rik | 1700021F05Rik | Gm47738 | Gm47900 | A330049N07Rik |
| Gm15939 | F930017D23Rik | Gm47915 | Gm47903 | Gm15398 |
| Sult3a2 | Gm34481 | Gm47917 | 1110002J07Rik | Gm20441 |
| Sult3a1 | Gm47927 | Gm36595 | Gm26576 | Gm5134 |
| Zufsp | Gm47926 | 4930452L12Rik | Gm31763 | Gm16220 |
| Gm48045 | Gm40634 | Gm48053 | AC155712.1 | Gm16222 |
| Fam26d | 1700027J07Rik | Gm36827 | Gm32023 | Gm16240 |
| Fam26e | 4933404K13Rik | Gm40652 | Gm47531 | 1700094J05Rik |
| Fam26f | Gm35028 | 4930467K11Rik | Gm32255 | 4930483K19Rik |
| Gm26564 | Gm48061 | AW822073 | 4930563J15Rik | Gm15343 |
| Gm47512 | Gm48065 | Gm4981 | Gm32364 | Gm35608 |
| 1700054O05Rik | Gm35154 | BB019430 | Gm32515 | Gm48276 |
| AC123035.1 | Gm48066 | Gm17542 | 1700030E10Rik | Gm35721 |
| Gm47049 | Gm47389 | 10-Sep | AC127568.1 | Gm3137 |
| 4930543K20Rik | 4930431F10Rik | Gm10273 | 4930521O17Rik | Gm10787 |
| Gm26535 | Gm47392 | Gm10322 | Gm16212 | AC153830.1 |
| Gm31378 | Gm47398 | Gm17059 | Gm47078 | Fam207a |
| 4930591E09Rik | Gm40638 | Pldi | 4930545H06Rik | 2810425M01Rik |
| Gm48197 | Gm40639 | Gm26947 | Gm32872 | Gm49325 |
| Wisp3 | Gm15934 | Gm20611 | 1700048P04Rik | Gm19402 |
| D030034A15Rik | Bves | Gm20625 | Gm33035 | Gm9508 |
| Gm31562 | D030045P18Rik | Gm20610 | Gm47107 | Gm10272 |
| Gm48486 | AC158799.1 | D830039M14Rik | Gm33263 | Gm10024 |
| Gm16364 | Gm35552 | Gm47595 | A930033H14Rik | Gm10142 |
| Gm16365 | Gm47568 | Gm47593 | Gm40685 | Gm10100 |
| Gm26824 | Gm47354 | H2afy2 | Gm7097 | Gm18596 |
| Mfsd4b3 | Gm48275 | Hk1os | Gm47946 | Gm9736 |
| Gm31992 | Gm46189 | 4930507D05Rik | Gm33979 | Gm3233 |
| Gm47598 | Gm49339 | Kif1bp | Gm48021 | Gm3238 |

| | | | | |
|---|---|---|---|---|
| Gm7138 | Gm15124 | 4930486F22Rik | 1110019B22Rik | Gm35722 |
| Gm3250 | Gm26710 | Gm15990 | Gm47708 | Gm47033 |
| Gm7137 | Gm49322 | Gm6729 | 4732465J04Rik | Gm47101 |
| Gm9639 | Gm26541 | Gm48485 | Gm47718 | Gm47143 |
| Gm19668 | Gm17151 | Gm49358 | Gm47719 | Gm17028 |
| Gm9507 | Gm16099 | Gm16268 | Gm33543 | Gm47223 |
| Gm2696 | 4930442H23Rik | Gm16280 | Gm47725 | Gm47221 |
| Gm36176 | Atcayos | Gm16271 | Gm5426 | Gm47224 |
| Gm10318 | Gm16315 | 4930555G07Rik | 4930459C07Rik | Gm47226 |
| Gm3285 | 4930404N11Rik | Igf1os | Gm48427 | Gm21293 |
| 1700009J07Rik | Gm48551 | Gm48306 | Gm33782 | Gm6763 |
| 1810043G02Rik | Gm16104 | 4921515L22Rik | Gm48428 | Gm8764 |
| Gm47922 | Gm16105 | Gm26546 | Gm48505 | Gm21304 |
| D10Jhu81e | Gm47697 | Gm16235 | Gm33843 | Gm21312 |
| Gm30122 | Aes | Uhrf1bp1l | Gm48521 | Gm20765 |
| 4921516A02Rik | Gm15917 | Gm26765 | Gm33981 | Gm4340 |
| Gm47944 | BC025920 | 4930534H03Rik | 4930556N09Rik | C230021G24Rik |
| Olfr1358 | Gm26896 | Gm47082 | 4930473O22Rik | 4930532I03Rik |
| Olfr1357 | Gm3055 | Gm31592 | Gm10754 | Gm29674 |
| Gm47459 | Gm4767 | Rmst | 4930525C09Rik | Gm29685 |
| Ccdc105 | Zfp781 | Gm26700 | C030005K15Rik | Gm36283 |
| Olfr1356 | BC024063 | Gm20757 | Gm8633 | Gm5136 |
| Olfr1355 | AU041133 | 4930401A07Rik | Gm34297 | 9230102K24Rik |
| Olfr1354 | AC158605.1 | Gm31822 | Gm5427 | 1700020G17Rik |
| Olfr8 | Zfp938 | Gm32468 | Phxr2 | Gm40761 |
| Olfr1353 | 4932415D10Rik | Gm32688 | Gm16239 | Gm28592 |
| Gm47343 | Gm4924 | Gm17745 | Gm34574 | 4933440J02Rik |
| Olfr1352 | Gm1553 | 4933408J17Rik | Gm48884 | Gm30262 |
| Olfr1351 | 1190007I07Rik | Gm47156 | Gm34777 | Gm30624 |
| Olfr57 | Gm47561 | Gm15915 | Gm34921 | Gm47865 |
| 2610008E11Rik | 1700028I16Rik | Gm15963 | Gm48089 | Gm20758 |
| Vmn2r80 | 1700025N21Rik | 4930471D02Rik | B530045E10Rik | Gm47419 |
| Vmn2r81 | Gm40723 | Gm4792 | Gm35035 | 1700010J16Rik |
| Vmn2r82 | D10Wsu102e | Tmcc3os | Gm34983 | Gm3942 |
| Gm47015 | C230099D08Rik | Gm48657 | Gm20110 | Gm47524 |
| Vmn2r83 | Gm34184 | Gm48689 | Gm35101 | Gm47532 |
| Theg | Gm34113 | Gm48867 | Gm47578 | Gm40765 |
| Odf3l2 | Gm47253 | Gm29684 | Gm35206 | 4930473D10Rik |
| Gm48495 | 4930463O16Rik | Gm33091 | AC167229.1 | Gm15723 |
| Gm17134 | E230014E18Rik | Gm48882 | Gm4301 | Gm31182 |
| Gm26602 | Platr7 | 2310039L15Rik | Gm4302 | Gm31267 |
| Atp5d | Fhl4 | Gm47599 | Gm4303 | A930009A15Rik |
| 1600002K03Rik | Btbd11 | Gm33336 | Gm4305 | AC153364.1 |
| Mum1 | Gm47358 | Gm47667 | Gm4307 | 4933416C03Rik |
| Gm15122 | Gm1559 | Anapc15-ps | Gm4312 | Kcnmb4os1 |
| Gm15123 | A230060F14Rik | Gm47671 | 4930430F08Rik | Kcnmb4os2 |

| | | | | |
|---|---|---|---|---|
| Gm26579 | 4930471E19Rik | A430046D13Rik | Olfr806 | Gm31401 |
| 1700030O20Rik | Gm48202 | Gm29585 | Olfr807 | Gm45076 |
| 4930579P08Rik | 4921513I03Rik | Gm47540 | Olfr808 | A630009H07Rik |
| Gm10271 | Gm15961 | Gm26876 | Olfr809 | Gm31343 |
| D630029K05Rik | Gm15910 | AC117232.1 | Olfr810 | D530014G21Rik |
| Gm10747 | Gm48341 | Gm47093 | Olfr811 | Gm31463 |
| 4930423D24Rik | Gm48410 | Mettl7b | Olfr812 | Gm45075 |
| 9530003J23Rik | 4930432O09Rik | 9030616G12Rik | Olfr813 | 4921522P10Rik |
| Gm48903 | Gm35404 | Olfr9 | Olfr814 | Gm44621 |
| Gm40770 | Gm48435 | Olfr763 | Olfr815 | Gm44626 |
| Gm32141 | Gm40787 | Olfr764-ps1 | Olfr816 | Gm44622 |
| Gm32235 | Gm46204 | Olfr765 | Olfr818 | Gm44515 |
| Gm32365 | Gm35696 | Olfr766-ps1 | Olfr819 | 4930453L07Rik |
| Gm32552 | Gm35865 | Olfr767 | Olfr247 | Gm44516 |
| Gm40773 | BC048403 | Olfr768 | Gm10310 | Gm39128 |
| 4933411E08Rik | Tmem5 | Olfr769 | Olfr820 | Fam155a |
| 5330439M10Rik | Gm4489 | Olfr770 | Olfr821 | Gm44761 |
| Gm47958 | Gm36041 | Olfr771 | Olfr822 | Gm10217 |
| Gm32717 | Gm48877 | Olfr772 | Olfr823 | Gm10067 |
| Gm32802 | A130077B15Rik | Olfr773 | Olfr824 | Gm44788 |
| Iltifb | Gm36208 | Olfr774 | Olfr825 | 5330413D20Rik |
| Gm9040 | Fam19a2 | Olfr775 | Olfr826 | Gm44786 |
| Gm9044 | Gm47839 | Olfr776 | Olfr827 | Gm44785 |
| Gm9045 | Gm36719 | Olfr777 | Vmn2r84 | Gm44784 |
| Gm9046 | Gm45604 | Olfr780 | Vmn2r85 | Gm45045 |
| Gm9048 | 1700021G15Rik | Olfr781 | Vmn2r86 | Gm39129 |
| Gm9049 | 1700025K04Rik | Olfr782 | Vmn2r87 | Gm32540 |
| Gm33337 | Gm40797 | Olfr784 | Gm31793 | B930025P03Rik |
| Gm47425 | Gm47966 | Olfr785 | Gm16180 | Gm45044 |
| Gm4065 | 4930484H19Rik | Olfr786 | A430078G23Rik | Gm2814 |
| Gm38403 | Gm49335 | Olfr787 | Tex45 | 3930402G23Rik |
| Gm47461 | Gm47972 | Olfr788 | C330021F23Rik | Gm44955 |
| Gm33677 | 9-Mar | Olfr789 | Fcor | Gm44956 |
| 1700064J06Rik | A730063M14Rik | Olfr790 | Gm39121 | B830042I05Rik |
| Gm47480 | F420014N23Rik | Olfr791 | Gm45231 | Gm44714 |
| Gm34045 | AC144852.1 | Olfr792 | Gm16553 | Gm44717 |
| 4930477N07Rik | Mars | Olfr794 | Gm26750 | Gm15418 |
| 1700025F24Rik | Gm47200 | Olfr796 | Cd209f | Gm15419 |
| Grip1os3 | Gm16217 | Olfr798 | Cd209g | 1700016D06Rik |
| Gm16321 | Gm16229 | Olfr799 | 4932443L11Rik | 4930465I24Rik |
| Gm47027 | Gm16230 | Olfr800 | Gm49320 | 1700128E19Rik |
| Grip1os2 | Rdh16f1 | Olfr801 | Gm6410 | Gm39132 |
| Grip1os1 | Gm47438 | Olfr802 | Gm44693 | Gm45680 |
| Gm47009 | Atp5b | Olfr803 | Gm17215 | B020031H02Rik |
| 1700006J14Rik | Gm26847 | Olfr804 | Gm30954 | A230072I06Rik |
| 9230105E05Rik | Gm17201 | Olfr805 | Gm31135 | Gm29175 |

| | | | | |
|---|---|---|---|---|
| Gm33175 | Gm6040 | Gm32050 | B930018H19Rik | Gm30931 |
| Gm33326 | Gm14851 | Gm45584 | Gm34096 | Gm45253 |
| 4933439N14Rik | Gm15293 | Gm32098 | Gm45627 | Gm45607 |
| Gm15348 | Gm15292 | Gm45646 | Gm34368 | Gm45479 |
| Gm15350 | Defa33 | 2310008N11Rik | Gm34419 | Gm31172 |
| Gm15347 | Gm7849 | Gm45572 | Gm45625 | 1700011L03Rik |
| Gm15353 | Gm7861 | Gm45371 | Gm34474 | Gm45600 |
| Gm15351 | AY761184 | Gm45370 | Gm34533 | 4930579M01Rik |
| Gm17023 | Defb50 | Gm32389 | Gm34597 | E330018M18Rik |
| Gm5608 | Defb2 | Proscos | Gm45301 | Gm45576 |
| 4932443I19Rik | Defb10 | 4933416M07Rik | Gm34853 | E030037K01Rik |
| AF366264 | Defb9 | Gm45267 | 1700015I17Rik | Gm31786 |
| Gm5907 | Defb11 | Gm45470 | Gm19410 | Gm45824 |
| C030037F17Rik | Gm36879 | Tex24 | 5430403N17Rik | Gm45864 |
| Gm16350 | Defb35 | Gm45861 | Gm38414 | 1700061N14Rik |
| 5830468F06Rik | 1810012K16Rik | Poteg | Gm39164 | Gm32052 |
| BB014433 | Gm26909 | Gm1698 | 4933430A20Rik | Gm45830 |
| Gm45274 | Gm15346 | 1700008N11Rik | Gm35520 | Gm45829 |
| Gm7706 | 1700041G16Rik | Gm8110 | 6430573F11Rik | 4930555F03Rik |
| Gm45271 | Gm15816 | Gm45571 | G630064G18Rik | Gm45321 |
| Gm35392 | A730045E13Rik | Gm26795 | Gm45524 | Gm45322 |
| Gm35453 | Gm45163 | Gm32567 | Gm45368 | Gm45323 |
| Gm45676 | Gm45164 | Gm39154 | Gm40493 | Gm45341 |
| Gm45408 | Gm30692 | Gm45490 | Gm6213 | Gm45455 |
| Gm16346 | Gm26714 | Gm45489 | Adam20 | Gm2516 |
| Gm16347 | Gm16933 | 4933433F19Rik | Gm26584 | C130073E24Rik |
| 1700014L14Rik | 5430421F17Rik | Fut10 | B430010I23Rik | Gm32975 |
| Gm16725 | Gm30978 | 7420700N18Rik | Gm16192 | Gm45389 |
| Gm35934 | Gm45237 | Gm45303 | Gm16193 | Gm45388 |
| Defb40 | Gm45238 | Gm45304 | 2810404M03Rik | 1190028D05Rik |
| Defb37 | Gm31045 | Gm26578 | Adam26b | Gm45554 |
| Defb38 | Gm16159 | Gm3985 | Adam26a | 5330439A09Rik |
| Defb39 | D830025C05Rik | Gm45587 | Gm5346 | 4930518J21Rik |
| Gm45826 | Gm17484 | 5930422O12Rik | Adam34 | Gm33501 |
| Spag11a | Gm45411 | 1700104B16Rik | Gm45488 | Gm34030 |
| Gm39139 | 1700047A11Rik | Gm26632 | Gm45659 | 5033428I22Rik |
| Defb46 | Gm45692 | Gm39157 | Gm2366 | Gm34099 |
| Gm35998 | Gm45250 | Gm9951 | AY512931 | Gm45359 |
| 4930467E23Rik | Gm45305 | Gm26768 | Gm30504 | Gm45356 |
| Gm26853 | Gm31727 | Gm10131 | Gm6329 | Gm26905 |
| Gm31371 | Gm31784 | Gm39158 | Gm45753 | Gm15882 |
| Gm26804 | Gm45307 | Gm29243 | Gm45542 | BC030500 |
| Gm15319 | Gm31898 | Gm45350 | Gm16351 | Gm34370 |
| Gm21119 | Gm45580 | Gm33831 | Sorbs2os | 4930470O06Rik |
| Defb33 | Gm31983 | Gm45349 | 1700029J07Rik | Gm34623 |
| Gm15056 | Gm39149 | Gm33968 | Gm39168 | Gm10283 |

| | | | | |
|---|---|---|---|---|
| Gm45650 | Fam129c | D830024N08Rik | Gm3235 | 1700047G07Rik |
| B230317F23Rik | Gm35572 | Ddx39 | Gm36325 | Gm45703 |
| Gm16178 | Zfp709 | Gm26721 | Gm36380 | Gm45788 |
| 1700001D01Rik | Zfp882 | Gm10644 | Irx3os | Gm39231 |
| 4930512H18Rik | Zfp961 | 2210011C24Rik | Gm28511 | Gm15679 |
| Gm45460 | Olfr372 | Gm10643 | Gm28515 | A330008L17Rik |
| Gm45459 | Olfr373 | C330011M18Rik | Gm36670 | Gm39232 |
| Gm45462 | Olfr374 | 4930432K21Rik | Gm36843 | Gm45391 |
| Gm35368 | Gm26586 | Gm26887 | Gm28516 | Gm32531 |
| Gm45589 | Gm11034 | Gm26532 | Gm45332 | 1600027J07Rik |
| Gm35521 | Gm10282 | Ccdc130 | Gm45335 | Gm45277 |
| Gm45518 | Gm17435 | Gm16183 | Gm45334 | 4933400L20Rik |
| Gm45345 | Gm45821 | Gm26664 | Gm21817 | Gm29682 |
| Gm32507 | Gm11033 | G430095P16Rik | Gm45336 | Gm10631 |
| Gm10663 | Gm45822 | THSD8 | Gm45505 | Gm45892 |
| BC030870 | Gm45820 | Asna1 | Gm45472 | Gm45711 |
| 1-Mar | Gm45899 | Gm5741 | Gm45765 | Gm11020 |
| Gm15354 | Gm45903 | Gm42031 | Gm45764 | K230015D01Rik |
| Gm15356 | Gm10358 | Cks1brt | Gm30132 | Gm45843 |
| Gm32568 | Gm45740 | Olfr371 | Gm45766 | Gm33023 |
| Gm45911 | Gm45741 | AA672651 | Gm45708 | Fam96b |
| Gm39185 | Gm45798 | Gm45425 | Gm26843 | 1700082M22Rik |
| Gm45827 | Gm45406 | Gm39214 | Gm30606 | 4932416K20Rik |
| Gm10660 | 4933421D24Rik | Gm10638 | 4930488L21Rik | Gm3830 |
| Gm15991 | Gm45285 | 2010110E17Rik | Gm45774 | D230025D16Rik |
| Gm26659 | 0610038B21Rik | Gm27167 | Gm39228 | 4931428F04Rik |
| 1700125H03Rik | 1700092C02Rik | Gm27168 | C78859 | Gm20163 |
| Gm15656 | Gm29895 | Gm5118 | Gm15889 | Lrrc29 |
| Gm33103 | Gm45326 | Gm27246 | Gm15890 | Gm38250 |
| Gm15716 | Gm39204 | Gm10637 | 9330175E14Rik | Gm5914 |
| D130040H23Rik | Gm30329 | Gm27169 | Fam192a | Gm16156 |
| Gm45848 | Gm45601 | Gm27207 | Gm45767 | Gm45752 |
| Gm9495 | Gm45602 | 4930535O05Rik | Gm31224 | Dpep2.1 |
| Gm43263 | Gm45435 | Gm2694 | Gm45812 | Dpep2nb |
| Gm7697 | Gm45431 | Gm19872 | Gm31036 | Gm26786 |
| Zfp868 | Gm45430 | Gm27016 | Tepp | Gm10629 |
| Zfp964 | 4930505O20Rik | 1700120C18Rik | 4933406B17Rik | 6030452D12Rik |
| Zfp869 | Gm31105 | 9430002A10Rik | Gm31518 | Gm10073 |
| Zfp963 | Gm27048 | Papd5 | Gm45702 | 1110028F18Rik |
| Gm20422 | Gm9725 | Gm45639 | 4930513N10Rik | Gm16208 |
| Gm11175 | Gm45538 | Gm35850 | Gm31659 | 5033426E14Rik |
| Gm16486 | Gm38590 | Gm35963 | Sap18b | Gm39244 |
| 2010320M18Rik | Gm31545 | Gm45641 | Gm31805 | Lncbate1 |
| 1700026F02Rik | Gm45449 | Gm45666 | Gm45895 | Gm20686 |
| Gm35256 | Gm10645 | Gm36243 | Gm32005 | Gm17344 |
| C430049E01Rik | Olfr370 | 4831440D22Rik | Gm32122 | 8030455M16Rik |

| | | | | |
|---|---|---|---|---|
| Gm17720 | Gm26784 | D830030K20Rik | Gm3500 | 4933406F09Rik |
| Gm21964 | Gm27030 | 1700110I01Rik | Gm3194 | Gm3755 |
| Gm26832 | Gm45833 | Gm2956 | Gm8246 | Gm3752 |
| Mtss1l | Gm20406 | Gm10413 | Gm3187 | Gm3558 |
| Gm15894 | Gm42047 | Gm10409 | Gm3488 | Gm10338 |
| Gm15895 | Gm26812 | Gm3005 | Gm3532 | Gm10128 |
| Gm26816 | Gm45353 | Gm10408 | Gm3383 | Gm16741 |
| Fuk | Trhr2 | Gm3012 | Gm8265 | Gm26661 |
| Aars | Gm20681 | Gm3020 | Gm9603 | Gm45521 |
| Kars | Gm20735 | Gm3043 | Gm3373 | Oit1 |
| Gm6793 | Gm10612 | Gm3002 | Gm3424 | 2610318M16Rik |
| Gm26994 | C230057M02Rik | Gm3033 | Gm8237 | 4930452B06Rik |
| Gm45904 | A530010L16Rik | Gm2916 | Gm3248 | 4930455B14Rik |
| Gm10280 | Gm45894 | Gm3029 | Gm3453 | Gm48370 |
| Gm30052 | Sult5a1 | Gm3095 | Gm8206 | Gm48371 |
| 4933408N05Rik | Gm4316 | Gm3099 | Gm6337 | Gm48439 |
| Gm16116 | Gm45842 | Gm5796 | Gm3468 | Fhitos |
| Gm16117 | 4933417D19Rik | Gm3115 | Gm3476 | Gm3839 |
| Gm16118 | Gm45781 | Gm8108 | Gm21560 | Gm48594 |
| 4930488N15Rik | 6030466F02Rik | Gm3127 | Gm3411 | Gm48595 |
| Gm15655 | 4732419C18Rik | Gm2974 | Gm21103 | Gm3848 |
| Gm45733 | Gm45866 | Gm3138 | Gm3618 | Gm49355 |
| 4930422C21Rik | 2810455O05Rik | Gm3149 | Gm16440 | 3830406C13Rik |
| 1700018P08Rik | Gm29773 | 2610042L04Rik | Gm3591 | D030051J21Rik |
| Gm15395 | Gm45890 | Gm3159 | Gm3594 | Gm48239 |
| Gm31717 | Gm45732 | Gm3164 | Gm8356 | Gm48526 |
| Gm38416 | 1810008B01Rik | Gm3173 | Gm3629 | Gm281 |
| Gm31774 | Gm15775 | Gm3182 | Gm3460 | Olfr720 |
| Gm45721 | 2810004N23Rik | Gm3047 | Gm10251 | Gm48857 |
| Gm45720 | Gm45709 | Gm8159 | Gm8050 | Olfr31 |
| Gm45723 | Gm16237 | Gm3239 | Gm3636 | Gm48860 |
| Gm32352 | 4930567H12Rik | Gm7876 | Gm8362 | Olfr721-ps1 |
| Tldc1 | A730098A19Rik | Gm3252 | Gm3642 | AC117608.1 |
| Gm20388 | Gm45875 | Gm9602 | Gm3667 | 4921522A10Rik |
| Gm15684 | Gm26759 | Gm3269 | Gm6356 | Gm47758 |
| Gm15898 | Gm45805 | Gm3264 | Gm10406 | Gm31577 |
| Fam92b | Gm31718 | Gm3278 | Gm3685 | Gm47782 |
| A330074K22Rik | A630001O12Rik | Gm8297 | Gm3696 | Gm31804 |
| Gm26739 | Gm3952 | Gm3298 | Gm6676 | Gm47780 |
| Gm27021 | Gm10999 | 4930555G01Rik | Gm3512 | 1700062C10Rik |
| Gm26537 | 2610044O15Rik8 | Gm8281 | Gm16434 | Gm47794 |
| Gm26878 | Gm32856 | Gm8279 | Gm3727 | Gm48163 |
| Gm28324 | Gm2888 | Gm3317 | Gm3739 | Gm48162 |
| 5033426O07Rik | Gm2897 | Gm8271 | Gm5797 | Gm48321 |
| Gm26815 | Gm10340 | Gm3526 | Gm8374 | Gm48099 |
| Gm26747 | Gm5795 | Gm3542 | Gm10339 | Gm21738 |

| | | | | |
|---|---|---|---|---|
| Gm48105 | Tmem254c | Gm5460 | Gm9611 | Gm5930 |
| Gm2244 | Cphx2 | A630023A22Rik | Gm7995 | Gm8229 |
| Gm2237 | Cphx3 | Gm30556 | Gm8005 | Gm3371 |
| Gm5458 | Duxbl2 | Gm49201 | Gm8011 | Gm8232 |
| Gm41102 | Plac9b | Fam35a | Gm8020 | AC242269.1 |
| 4930469B13Rik | Tmem254b | Fam25c | Gm8024 | BC061237 |
| D14Ertd670e | Duxbl3 | Gm17110 | Gm17124 | Gm17093 |
| Gm48283 | Gm34059 | 9230112D13Rik | Gm3633 | AC160336.1 |
| Gm48395 | 4933413J09Rik | Gm49012 | Gm10378 | Gm5624 |
| Gm30054 | Slmapos2 | Gm49024 | Gm3015 | Gm8247 |
| 7330404K18Rik | Arf4os | Gm49025 | Gm10377 | Gm8257 |
| 1810062O18Rik | 9930004E17Rik | A930038B10Rik | Gm21977 | Gm8267 |
| 2810402E24Rik | Fam208a | Gm49032 | Gm8082 | Gm34060 |
| 6230400D17Rik | Gm45645 | 4930596D02Rik | Gm10376 | Gm49145 |
| Fut11 | Gm2670 | 4930474N05Rik | Gm8094 | Gm17173 |
| Gm30108 | Gm35164 | 2610528A11Rik | Gm8104 | Gm49180 |
| Dupd1 | Gm35217 | Gm31393 | Gm49135 | Ero1l |
| Dusp13 | Gm48004 | Gm31456 | Gm7233 | Gm34250 |
| Gm15935 | Gm48006 | Gm20641 | Gm8122 | Gm41144 |
| A430057M04Rik | Gm48003 | Nrg3os | Gm8126 | Gm49123 |
| 4931407E12Rik | Gm48058 | Gm20642 | AC166344.1 | Gm15601 |
| Gm7480 | Gm41118 | 1700109I08Rik | Gm8127 | Gm49124 |
| 4930405A10Rik | Gm35281 | Fam213a | 1700001F09Rik | A530076I17Rik |
| Gm10248 | Gm47414 | Gm47547 | Gm8138 | Gm15218 |
| Gm29626 | AC154646.1 | Gm2832 | Gm5799 | Gm49192 |
| CT025624.1 | 1700087M22Rik | Gm5798 | Gm17654 | Gm15217 |
| E330034G19Rik | Tmem110 | Gm7945 | Gm10375 | Gm15222 |
| Gm47774 | Smim4 | Gm6482 | Gm21154 | Gm15221 |
| Gm47814 | Gm35823 | Gm7954 | Gm8165 | Gm48910 |
| 4930542C16Rik | Gm49217 | Gm3486 | Gm16506 | Gm48911 |
| 4930428N03Rik | AC108416.1 | Gm7970 | Gm6526 | Gm34756 |
| Gm47906 | Gm49387 | 1700049E17Rik2 | AC154806.1 | Gm48912 |
| Gm10398 | D830044D21Rik | Gm3072 | Gm8180 | E130120K24Rik |
| Gm38408 | 4930425P05Rik | Gm3676 | Gm41138 | Rubie |
| Gm32123 | Gm17210 | Gm8068 | Gm49083 | 4933425B07Rik |
| Gm32224 | Gm46447 | Gm7929 | Gm8113 | D330046F09Rik |
| 4930572O13Rik | Gm48914 | Gm47189 | Gm49087 | Gm48924 |
| Zmiz1os1 | Gm2990 | Gm7980 | Gm3287 | Gm48933 |
| Gm26660 | Prrxl1 | 1700049E17Rik1 | Gm49099 | Gm34934 |
| Gm17747 | Gm28651 | 1700024B05Rik | Gm49101 | 2810457G06Rik |
| Gm26772 | 1810011H11Rik | Gm6401 | Gm3327 | Gm10371 |
| Ppifos | Gm49030 | Gm3543 | 4930503E14Rik | Gm49150 |
| Anxa11os | Gm15512 | Gm7951 | 1700047E10Rik | Gm49189 |
| Plac9a | Gm49108 | Gm17027 | Gm8212 | Gm48975 |
| Tmem254a | 4930503F20Rik | Gm3573 | Gm32857 | Gm35166 |
| Gm9780 | Gm30083 | Gm17026 | Gm8220 | Gm49004 |

| | | | | |
|---|---|---|---|---|
| 4930538L07Rik | Olfr748 | Trav3n-2 | Traj30 | Gm26751 |
| Gm41148 | Olfr749 | Trav4n-3 | Traj29 | Gm48925 |
| Gm49302 | Gm26782 | Trav5n-2 | Traj28 | Cma2 |
| Gm35360 | Tmem55b | Trav5n-3 | Traj27 | Mcpt1 |
| 4930447J18Rik | Gm49342 | Trav4n-4 | Traj26 | Mcpt9 |
| Gm49303 | Olfr750 | Gm43650 | Traj25 | Mcpt2 |
| Gm49304 | 1810028F09Rik | Trav15n-3 | Traj24 | Mcpt4 |
| Gm49305 | Gm21718 | Trav4-2 | Traj22 | Mcpt8 |
| Gm49306 | AC125117.1 | Trav3-2 | Traj21 | Gm4022 |
| 1700128I11Rik | Gm7247 | Trav4-3 | Traj20 | Gm10873 |
| 4930572G02Rik | Vmn2r88 | Trav5-2 | Traj19 | Cenpj |
| Gm49120 | Vmn2r89 | Trav4-4-dv10 | Traj18 | Gm16573 |
| Gm35766 | Gm5622 | Trav13-3 | Traj16 | Gm16973 |
| Gm49151 | Gm17175 | Trav8-2 | Traj15 | B020004C17Rik |
| Gm49156 | Gm17174 | Trav15-3 | Traj14 | Gm4491 |
| Gm48935 | Gm17078 | Trav12-4 | Traj13 | Ift88os |
| 1700011H14Rik | Gm4181 | B230359F08Rik | Traj12 | Gm49361 |
| Gm48940 | Gm17079 | Trav20 | Traj11 | Gm33321 |
| 3632451O06Rik | Gm5800 | Gm43647 | Traj9 | Gm48447 |
| Gm36146 | AY358078 | Trav22 | Traj8 | Gm33472 |
| Olfr722 | Gm49076 | Trdv3 | Traj6 | Gm41162 |
| Olfr723 | Gm16617 | Gm30275 | Traj5 | Gm48453 |
| Olfr724 | Gm49086 | Trdd1 | Traj4 | 1700129C05Rik |
| Olfr725 | Gm49256 | Trdd2 | Traj3 | Gm5142 |
| Olfr726 | Olfr221 | Gm43434 | Traj2 | Gm6904 |
| Olfr727 | Gm26590 | Traj61 | Traj1 | 4930563I02Rik |
| Olfr728 | Olfr1513 | Traj60 | Olfr49 | Nupl1 |
| Olfr729 | Olfr1512 | Traj59 | 5330426L24Rik | Gm49336 |
| Olfr730 | Olfr1511 | Traj58 | Gm10366 | Gm29266 |
| Olfr731 | Olfr1510 | Traj56 | Gm17606 | Mipepos |
| Olfr732 | Olfr1509 | Traj55 | 4930579G18Rik | AC124555.1 |
| Olfr733 | Olfr1508 | Traj54 | Gm29776 | 1700109G14Rik |
| Olfr734 | Olfr1507 | Traj51 | Gm20521 | 4930556J02Rik |
| Olfr735 | Trav4-1 | Traj50 | Gm49130 | Gm27010 |
| Tlr11 | Trav4d-2 | Traj49 | Gm29015 | Gm26916 |
| Olfr736 | Trav14d-1 | Traj47 | Gm17428 | Gm27017 |
| Olfr738 | Trav3d-2 | Traj46 | Gm31251 | Gm35419 |
| Olfr739 | Trav4d-3 | Traj45 | Zfhx2os | Gm26969 |
| Olfr740 | Trav5d-2 | Traj44 | Gm20687 | 4931440J10Rik |
| Olfr741 | Trav5d-3 | Traj43 | Gm49022 | Gm26536 |
| Olfr742 | Trav4d-4 | Traj42 | Gm10876 | Gm49390 |
| Olfr743 | C920008G01Rik | Traj41 | A730061H03Rik | Gm5463 |
| Olfr744 | Gm30214 | Traj37 | Gm15932 | Gm17232 |
| Olfr745 | Trav15d-3 | Traj36 | Gm49378 | Gm15918 |
| Olfr746 | Trav9n-1 | Traj35 | AC174678.1 | 4930471C04Rik |
| Olfr747 | Trav11n | Traj34 | Gm49067 | Gm17116 |

| | | | | |
|---|---|---|---|---|
| 9630015K15Rik | Gm49164 | Gm41225 | Gm4675 | Gm32710 |
| Gm47202 | Gm49171 | Gm48941 | Gm9376 | Gm47001 |
| Gm30008 | 4933402J15Rik | 4930474H20Rik | Gm32093 | Gm47002 |
| Gm41177 | Gm49175 | Gm33246 | Gm26791 | Gm47034 |
| Gm48433 | Gm10847 | Gm48942 | 1700044C05Rik | 4930568E12Rik |
| Gm48449 | Gm4278 | 4930433E05Rik | Gm27198 | Gm47051 |
| Gm41183 | Gm15628 | Gm33450 | Gm49179 | Gm17571 |
| Gm10032 | Gm15629 | 4921530L21Rik | AC154760.1 | Gm47535 |
| Gm5464 | Gm49200 | Gm49007 | Gm4681 | Gm47557 |
| AC166061.1 | Spert | Gm33525 | Gm48997 | 1700019J19Rik |
| Gm20675 | Gm4285 | Gm49018 | Gm26679 | Gm47559 |
| Gm6878 | 2900040C04Rik | Gm41230 | Gm49008 | Gm47564 |
| 4930438E09Rik | 4930444M15Rik | Gm49225 | Gm17613 | Gm47565 |
| Gm10860 | E130202H07Rik | 4930517O19Rik | Gm41253 | Gm34069 |
| Gm38409 | 4930431P22Rik | Gm41231 | Gm33299 | Gm47326 |
| Gm31107 | Gm30246 | 6330576A10Rik | Gm49035 | Gm47319 |
| Gm47212 | Gm48968 | Gm49237 | 1810041H14Rik | Gm34263 |
| Gm31227 | Gm48969 | Gm9922 | Timm8a2 | Gm7607 |
| Gm41192 | Gm49002 | Gm49292 | Gm49236 | Gm10706 |
| 1700092C10Rik | Gm6994 | Gm26778 | A330035P11Rik | Gm16302 |
| Gm27222 | Gm49003 | 4933432I03Rik | 1700108J01Rik | Gm47214 |
| Gm27223 | Gm30467 | AC102815.1 | AC154683.1 | Gm16568 |
| Gm16677 | Gm1587 | AC114585.1 | Gm5089 | Gm34973 |
| Gm27221 | Gm48954 | Gm34907 | Gm10837 | 4930540M03Rik |
| Gm16867 | Gm19301 | Slain1os | 4930594M22Rik | Olfr24 |
| Gm27175 | Gm49016 | 4930432J09Rik | Gm28932 | Olfr828 |
| Gm27179 | Gm49015 | D130079A08Rik | 1700024B18Rik | Olfr829 |
| Gm27174 | Gm4632 | Rnf219 | 4930404O17Rik | Olfr830 |
| Gm27176 | Zfp957 | D130009I18Rik | Gm26870 | Olfr832 |
| Gm27177 | Gm49042 | 5430440P10Rik | Gm10722 | Olfr834 |
| 4930480K23Rik | 4930452G13Rik | Gm35909 | Gm11168 | Olfr835 |
| Gm33524 | Gm9748 | Gm10076 | Gm10721 | Olfr836 |
| AC151836.1 | Gm6999 | Gm48970 | Gm10720 | Olfr837 |
| Fam160b2 | Gm10845 | Gm48972 | Gm10719 | Olfr839-ps1 |
| 2410012E07Rik | AC121960.1 | Gm36107 | Gm10718 | Olfr843 |
| Gm49289 | AC121960.2 | 1700128A07Rik | Gm10717 | Olfr844 |
| Gm4251 | Gm49046 | Gm48977 | Gm17535 | Olfr845 |
| 4930429C20Rik | Gm41219 | Gm48992 | Gm10715 | Olfr846 |
| 4930434J06Rik | Gm49048 | Tpm3-rs7 | Gm31138 | Olfr847 |
| Gm9195 | Gm49054 | 4930524C18Rik | AV064505 | Olfr849 |
| Gm16549 | Gm32455 | Gm49010 | 9230110C19Rik | Olfr850 |
| Gm49293 | 9630013A20Rik | 1700100I10Rik | Gm32014 | Olfr851 |
| Gm41206 | Gm32729 | Gm31072 | Gm16485 | Olfr853 |
| Gm17233 | 4930529K09Rik | 4930435M08Rik | Gm16833 | Olfr854 |
| Gm21750 | Gm32913 | Gm26773 | Gm46102 | Olfr855 |
| Gm49165 | Gm33203 | 4933431J24Rik | Gm47334 | Olfr857 |

| | | | | |
|---|---|---|---|---|
| Olfr58 | Gm48256 | 4933422A05Rik | Olfr250 | Olfr148 |
| Olfr859 | Gm10181 | Gm33838 | Olfr251 | Olfr960 |
| Gm45699 | Gm29642 | Gm1113 | Olfr147 | Olfr961 |
| Olfr860 | 7-Sep | Gm17677 | Olfr901 | Olfr963 |
| Olfr862 | Gm48362 | Gm17727 | Olfr902 | Olfr149 |
| Olfr77 | Gm39312 | Gm27235 | Olfr904 | Olfr965 |
| Olfr866 | Gm30313 | D730048I06Rik | Olfr905 | Olfr150 |
| Olfr867 | Gm48393 | 9230110F15Rik | Olfr906 | Olfr967 |
| Olfr868 | Gm1110 | 9230113P08Rik | Olfr907 | Olfr968 |
| Olfr869 | Gm48796 | Gm27187 | Olfr911-ps1 | Olfr969 |
| Olfr870 | Gm48801 | Gm5916 | Olfr910 | Olfr970 |
| Olfr871 | Gm48803 | Gm3434 | Olfr912 | Olfr971 |
| Olfr872 | Gm48901 | Gm7257 | Olfr913 | Olfr972 |
| Olfr39 | Gm48902 | Gm9513 | Olfr914 | Olfr229 |
| Olfr873 | Gm48091 | Gm5615 | Olfr916 | Olfr974 |
| Olfr18 | Gm15520 | Gm17689 | Olfr917 | Olfr975 |
| Gm26733 | Gm47677 | A630095E13Rik | Olfr918 | Olfr976 |
| Fbxl12os | Gm47716 | Gm26787 | Olfr919 | Olfr978 |
| Gm26521 | Gm31013 | Gm3896 | Olfr920 | Olfr979 |
| A230050P20Rik | Gm29724 | Gm48716 | Olfr921 | Olfr980 |
| Gm47079 | Gm17508 | Gm48081 | Olfr922 | Olfr981 |
| Gm26592 | CT025678.1 | Gm47633 | Olfr923 | Olfr982 |
| Gm49373 | Gm31408 | Gm34885 | Olfr924 | Olfr983 |
| Gm38431 | Gm47436 | Olfr160 | Olfr26 | Olfr984 |
| Fdx1l | Gm31497 | Olfr874 | Olfr926 | Olfr985 |
| 1700084C06Rik | Gm47465 | Olfr875 | Olfr930 | Olfr986 |
| Gm16754 | Gm39317 | Olfr876 | Olfr933 | 1700110K17Rik |
| Gm39302 | Gm27201 | Olfr877 | Olfr934 | Gm17540 |
| Gm36198 | Gm47680 | Olfr145 | Olfr935 | Gm35177 |
| Gm16853 | Gm27240 | Olfr878 | Olfr146 | Gm48277 |
| Gm26511 | Gm3331 | Olfr881 | Olfr937 | Gm48284 |
| Ccdc151 | Gm27162 | Olfr883 | Olfr938 | Gm35371 |
| Gm49318 | Gm27166 | Olfr884 | Olfr27 | Gm21915 |
| Gm16845 | Gm32171 | Olfr885 | Olfr943 | Gm16095 |
| 1810064F22Rik | Gm47775 | Olfr887 | Olfr944 | Gm48293 |
| Zfp809 | Gm47777 | Olfr888 | Olfr1537 | Gm35657 |
| Zfp599 | Gm47778 | Olfr889 | Olfr945 | 1700063D05Rik |
| Zfp810 | Gm47784 | Olfr890 | Olfr948 | Gm48742 |
| 9530077C05Rik | Gm47785 | Olfr891 | Olfr951 | Gm48740 |
| Gm17545 | Gm27164 | Olfr893 | Olfr952 | CT025619.1 |
| Gm47663 | 7630403G23Rik | Olfr894 | Olfr954 | Gm48784 |
| Gm15600 | Gm47858 | Olfr143 | Olfr955 | Gm35835 |
| Gm10701 | Gm26662 | Olfr895 | Olfr44 | Gm40513 |
| Cypt4 | Kirrel3os | Olfr896-ps1 | Olfr957 | Gm16214 |
| Gm29824 | Gm39318 | Olfr25 | Olfr958 | Gm39321 |
| E130101E03Rik | Srpr | Olfr898 | Olfr959 | 4930546K05Rik |

| | | | | |
|---|---|---|---|---|
| Gm36435 | Gm10680 | Gm48346 | 4933433G08Rik | Gm27241 |
| Gm47629 | Gm47544 | Gldnos | I730028E13Rik | Gm27232 |
| Gm16322 | 4931429L15Rik | Wdr61 | Gm34105 | Gm27204 |
| Gm47931 | Gm39329 | AY074887 | B930092H01Rik | Dyx1c1 |
| Tmem136 | Gm31374 | Gm47877 | Gm10655 | Gm44503 |
| D630033O11Rik | Gm47141 | Gm47887 | Gm34322 | Ccpg1os |
| Gm36799 | Gm47144 | Gm17226 | Gm47262 | Gm20509 |
| Gm47475 | Gm31432 | 4930563M21Rik | A730043L09Rik | Gm27211 |
| Gm28215 | Gm48945 | Peak1os | Gm35288 | Gm28622 |
| Gm36855 | Gm31557 | Gm47176 | Gm26609 | Gm20649 |
| Gm29909 | Gm31614 | Gm26868 | Gm48193 | Gm28182 |
| Gm29961 | 2900052N01Rik | Gm47235 | Gm36033 | Fam214a |
| Gm3898 | Gm47153 | Odf3l1 | 1110036E04Rik | Gm19531 |
| Gm30373 | Gm4791 | Gm47237 | Gm48855 | Gm48677 |
| Gm10688 | Gm31698 | 4930442G15Rik | Gm47350 | 4933433G15Rik |
| Gm39323 | Gm31816 | Gm10658 | Gm39363 | Gm26903 |
| Gm49380 | A730065G17Rik | 2700012I20Rik | Gm47270 | Gm27234 |
| Gm39325 | Gm40518 | Man2c1os | Fam96a | Gm34158 |
| Gm26737 | Gm32335 | 1700041C23Rik | Gm28379 | Gm47800 |
| Gm10687 | 1700042D02Rik | Trcg1 | Gm15563 | Gm19541 |
| Pdzd3 | Gm4894 | Gm17231 | Gm10647 | 5730403I07Rik |
| Gm48604 | Gm11149 | Gm17322 | BC050972 | Gm47790 |
| Gm48853 | Gm47543 | Gm16130 | Gm17098 | Gm47789 |
| H2afx | 2310003N18Rik | Gm16131 | Gm10646 | Gm47795 |
| Ccdc84 | Gm39336 | 1700072B07Rik | Gm19299 | Gm34654 |
| Gm47186 | Rpl10-ps3 | 4930461G14Rik | Gm47047 | Gm47818 |
| C030014I23Rik | Plet1os | 6030419C18Rik | M5C1000I18Rik | Gm34829 |
| Gm47204 | Gm47077 | Gm47271 | Gm20745 | Gm39375 |
| Gm47230 | AU019823 | Gm33053 | Gm28195 | Ick |
| Gm47231 | 1110032A03Rik | Gm33180 | 4930502A04Rik | C920006O11Rik |
| Gm47232 | 4833427G06Rik | Gramd2 | Gm15511 | Gsta4 |
| BC049987 | Gm32742 | Gm20275 | Gm16144 | Gm3776 |
| Gm48702 | Colca2 | Gm7616 | B230323A14Rik | Gm10639 |
| AC061963.1 | 2010007H06Rik | Gm15507 | Gm28731 | Omt2a |
| Atp5l | 1810046K07Rik | 2010001M07Rik | 2210414F02Rik | Omt2b |
| Gm30934 | Gm47854 | 9230112J17Rik | Gm26849 | 4930542C12Rik |
| Gm48719 | Arhgap20os | Gm49334 | Gm10642 | Mb21d1 |
| Gm10684 | Gm6980 | B930082K07Rik | Gm32511 | Gm17324 |
| Gm39327 | 4933407I05Rik | 1700036A12Rik | Gcom1 | Gm47430 |
| Gm17099 | Gm47957 | Gm9869 | Gm27159 | 4921509A06Rik |
| BC049352 | Gm47960 | Gm47923 | Gm27188 | Gm35024 |
| 1700003G13Rik | AC166902.1 | Gm5122 | Gm27253 | Gm47499 |
| 4833428L15Rik | Gm1715 | AC160562.1 | BC065403 | Gm47501 |
| Gm16536 | 4930510E17Rik | Gm39348 | Gm27255 | Gm28620 |
| Gm48529 | Kdelc2 | Gm34004 | Gm27231 | Gm39377 |
| A830035O19Rik | Gm16124 | Gm33914 | 4930509E16Rik | Gm10635 |

| | | | | |
|---|---|---|---|---|
| 4930562D21Rik | Gm29562 | Gm28087 | Gm34106 | Gm4665 |
| 4930429F24Rik | Gm26611 | Gm16004 | Gm9917 | Gm26962 |
| Gm17477 | 4930579C12Rik | Gm47757 | Gm16343 | Gm48007 |
| Gm27227 | Gm29094 | 4930519F24Rik | Nat6 | Gm16142 |
| Gm48387 | Gm47403 | Gm28167 | Gm38150 | Gm4668 |
| Gm39380 | AF529169 | Gm34397 | Gm37850 | Gm48038 |
| Gm27224 | Gm47409 | Gm28166 | Gm37974 | Gm31410 |
| Gm26713 | 1700017I07Rik | Gm28979 | Gm20661 | D730003K21Rik |
| Gm27226 | 4930524O08Rik | 9630041A04Rik | Fam212a | Gm45881 |
| D430036J16Rik | Gm26882 | Gm26846 | 4921523L03Rik | Gm26614 |
| Gm26907 | Gm30849 | Gm29387 | Gm20662 | Gm17399 |
| Gm35548 | Gm28703 | 4930533D04Rik | 4930535L15Rik | Platr11 |
| 1700063H06Rik | Gm29408 | Gm5627 | Gm38134 | 4933432G23Rik |
| Gm39383 | Gm28652 | Gm47416 | Gm37401 | Gm33460 |
| Gm38398 | Gm29478 | Gm47468 | Ccdc36 | Gm45897 |
| Gm27216 | Gm31409 | Gm20425 | Qars | Gm2415 |
| Gm46123 | Gm29139 | 1300017J02Rik | 4833445I07Rik | Gm16295 |
| Gm2087 | 1700057G04Rik | Gm47328 | Gm38004 | Gm47289 |
| Gm39384 | 4933400C23Rik | 5830418P13Rik | Gm7628 | Gm47320 |
| Gm36120 | Gm28054 | Gm16252 | Gm42493 | 4930516B21Rik |
| Gm2109 | 1700034K08Rik | Gm32743 | Gm42756 | 1700019L13Rik |
| Gm36278 | 1190002N15Rik | 4932413F04Rik | Gm43732 | Gm47043 |
| Gm46124 | Gm29395 | Gm29154 | Gm42775 | AC117245.1 |
| Gm39388 | Gm16262 | Gm33054 | Ngp | Gm33858 |
| 4930554C24Rik | Gm28424 | Gm28305 | Gm42470 | Gm5922 |
| Gm48820 | Gm32281 | Acpp | Gm35715 | Gm38642 |
| Gm28625 | Gm16126 | Gm28548 | Tdgf1 | Gm47059 |
| Gm48821 | 1700065D16Rik | Gm47106 | 1700061E17Rik | D830035M03Rik |
| Gm48830 | Gm16794 | 1700080E11Rik | Gm42523 | 5830454E08Rik |
| Gm48831 | C78334 | Gm20643 | Gm43236 | Gm39456 |
| Gm11114 | Gm28085 | Gm15619 | 4930545L08Rik | Gm47064 |
| Fam46a | BC043934 | Gm19667 | 2310075C17Rik | Gm47066 |
| Gm48832 | Gm28729 | Gm47575 | 2900079G21Rik | Gm39459 |
| Gm48833 | A930006L05Rik | Gm29208 | 4933411E02Rik | Gm47068 |
| Gm48834 | Gm16010 | D030055H07Rik | Gm36251 | Gm47069 |
| Gm48842 | Gm19325 | 4930500F10Rik | Gm4657 | Gm19385 |
| Gm47604 | Gm26767 | Gm29123 | Gm36367 | Gm34425 |
| Ube2cbp | Gm16185 | Gm28959 | AC164881.1 | Gm47070 |
| Dopey1 | Gm15891 | Iqcf3 | Gm36485 | Gm39460 |
| A330041J22Rik | 4921534H16Rik | Gm28111 | Gm47875 | 1700020M21Rik |
| Gm36539 | 2610303G11Rik | 4930524O07Rik | AU023762 | 4930593C16Rik |
| Gm28231 | Gm37637 | 1700039M15Rik | Gm36745 | Gm47095 |
| Gm20537 | E330023G01Rik | Gm17041 | Gm47950 | Gm38661 |
| Zfp949 | 7420426K07Rik | 4930429P21Rik | Gm29825 | Lyzl4os |
| 9430037G07Rik | Gm1123 | Gm17141 | C130032M10Rik | E530011L22Rik |
| Gm10634 | 4930422M22Rik | Gm17118 | Gm9888 | Zfp651 |

| | | | | |
|---|---|---|---|---|
| Gm17163 | Gm11998 | Gm6685 | Gm12159 | Gm12660 |
| Gm47112 | 4930512M02Rik | Gm12081 | Gm12160 | 4933414I15Rik |
| Gm47113 | Gm12000 | 2810471M01Rik | Gm12162 | Olfr1378 |
| 1700048O20Rik | 1700042O10Rik | Gm12082 | Gm12166 | Olfr1377 |
| Gm49343 | Gm12002 | Gm12089 | Gm16033 | Olfr51 |
| Fam198a | 4930554G24Rik | Gm12088 | Gm16034 | Olfr54 |
| Gm39463 | 1700046C09Rik | Gm12092 | Gm12167 | Olfr1375 |
| Gm47117 | Egfros | 4931440F15Rik | Fam71b | 9230009I02Rik |
| Gm39465 | Wdr92 | Gm29237 | 2310031A07Rik | Platr20 |
| Gm39464 | Etaa1os | 4930505A04Rik | Gm12169 | Gm12569 |
| Gm26797 | Gm12018 | Gm17332 | Gm12171 | BC049762 |
| A730085K08Rik | Gm12022 | D630024D03Rik | BC053393 | D930048N14Rik |
| Gm47128 | Gm12023 | Gm12107 | Dppa1 | 0610009B22Rik |
| Gm35454 | Gm28401 | Gm12108 | Gm4926 | Gm39822 |
| Gm47131 | Gm16140 | 4930555O08Rik | Gm12184 | Gm26551 |
| Gm47133 | Gm16141 | Gm12111 | Gm16170 | Gm12204 |
| 9530059O14Rik | 4933406G16Rik | Gm12114 | Gm5431 | Gm12205 |
| Gm47134 | Gm12027 | Gm12116 | Gm12185 | 9430098F02Rik |
| Gm35501 | Gm12031 | Gm12117 | 9930111J21Rik1 | Olfr1373 |
| Gm47136 | Gm12037 | D130052B06Rik | Olfr56 | Olfr1371 |
| Gm9856 | 4932414J04Rik | Gm12119 | Tgtp1 | Skp1a |
| Gm47162 | Gm12050 | Gm12120 | 9930111J21Rik2 | Gm12207 |
| Gm17120 | Gm12051 | Gm12121 | Tgtp2 | A630014C17Rik |
| Gm17200 | Gm26829 | 4930403D09Rik | Ifi47 | Gm11186 |
| Gm39469 | Gm12052 | Fam196b | Olfr1396 | Gm10447 |
| Gm47181 | Gm12055 | Gm12711 | Olfr1395 | Gm12212 |
| Gm21836 | Gm12056 | Rars | Olfr1394 | Gm9837 |
| Gm11399 | Gm28048 | Gm12126 | Olfr1393 | Gm9945 |
| Gm12735 | Zrsr1 | Gm12127 | Olfr1392 | 8-Sep |
| 4930556J24Rik | 1700061J23Rik | Gm12128 | Olfr10 | Gm12214 |
| Gatsl3 | 0610010F05Rik | Gm12130 | Olfr1391 | Gm17334 |
| Gm11959 | Gm12061 | Gm12132 | Olfr1390 | Gm12216 |
| Gm11958 | A830031A19Rik | 3110004A20Rik | Olfr1389 | 4933405E24Rik |
| Gm11032 | Gm12063 | Platr13 | Olfr1388 | Gm12223 |
| Gm11961 | Gm12064 | Gm9972 | Olfr1387 | Gm12224 |
| Gm11963 | 4930538E20Rik | Gm12145 | Olfr1386 | Gm12227 |
| Ankrd36 | Gm12065 | Gm12146 | Olfr1385 | Lyrm7os |
| Gm11967 | Gm12066 | Gm12144 | Olfr1384 | Slc36a3os |
| A730071L15Rik | Gm12068 | Gm12147 | Olfr1383 | Slc36a1os |
| H2afv | 4933430M04Rik | Gm12148 | Olfr1382 | Gm12236 |
| Gm11973 | Gm10466 | Gm12150 | Olfr1381 | Gm12239 |
| Wap | Gm12665 | 4933415A04Rik | Olfr1380 | Gm12243 |
| Gm11983 | 4933427E13Rik | Gm12153 | Gm12195 | 4933424L21Rik |
| Gm11985 | Gm12069 | Gm12154 | Gm26542 | 4933426K07Rik |
| Gm11986 | 5730522E02Rik | 4930597A21Rik | Gm12199 | Gm12249 |
| Spata48 | Gm6899 | Gm12158 | Gm12200 | Gm12248 |

| | | | | |
|---|---|---|---|---|
| Gm12246 | Gm12265 | Kdm6bos | Zfp616 | Gm11418 |
| 4930438A08Rik | Myo15 | Dnah2os | Olfr401 | 5530401A14Rik |
| 1810065E05Rik | Gm26837 | Tnfsf13os | Olfr402 | Gm31522 |
| Gm12253 | Gm26534 | Tnfsfm13 | Olfr403 | Gm11419 |
| 4930504O13Rik | Map2k3os | Gm39566 | Olfr43 | Gm17268 |
| Olfr30 | Gm45605 | 2810408A11Rik | Olfr406 | Gm38577 |
| Olfr332 | B9d1os | 0610010K14Rik | Olfr59 | Gm11426 |
| Olfr331 | Gm12272 | Gm21988 | Olfr410 | Gm11423 |
| Olfr330 | A530017D24Rik | Alox12e | Olfr411 | Slfn5os |
| Olfr329-ps | Gm26964 | Gm12316 | Olfr412 | 1190001M18Rik |
| Olfr328 | Gm12273 | Gm12315 | Gm28180 | Gm20234 |
| Olfr224 | Gm12278 | Gm12319 | Gm12333 | Gm48462 |
| Olfr325 | Gm12949 | Gm12320 | Gm12336 | Rpl9-ps1 |
| Olfr324 | Gm12279 | 6330403K07Rik | Gm45606 | Wfdc17 |
| 2210407C18Rik | Gm12280 | Gm16013 | Gm26836 | Wfdc18 |
| Olfr323 | Lrrc75aos2 | 9230020A06Rik | Gm16075 | Gm12576 |
| Olfr322 | Lrrc75aos1 | Gm33351 | 1700016K19Rik | Lhx1os |
| Olfr320 | Mmgt2 | Gm12324 | Fam57a | 1700109G15Rik |
| Olfr319 | Gm27194 | Gm12325 | Gm26780 | Gm11431 |
| Olfr318 | Gm10428 | Gm12326 | Tusc5 | 1700125H20Rik |
| Olfr317 | Zfp286os | 4930401O10Rik | Gm12343 | Appbp2os |
| Olfr316 | Tvp23bos | Gm43951 | Gm45027 | Bcas3os1 |
| Olfr315 | Cdrt4os2 | 4732414G09Rik | 2210008F06Rik | Bcas3os2 |
| Olfr314 | Cdrt4os1 | Gm49340 | Gm10392 | Gm11444 |
| Fam183b | B430319H21Rik | Olfr20 | Gm10277 | Brip1os |
| Olfr313 | B130011K05Rik | Olfr376 | Gm11190 | Gm11479 |
| Olfr312 | 9630013K17Rik | Olfr1 | Dhrs13os | Gm11491 |
| Olfr311 | Gm12289 | Olfr378 | 2610507B11Rik | Gm11492 |
| Gm12258 | 2810001G20Rik | Olfr380 | Gm11192 | 4-Sep |
| Gm12259 | Gm12290 | Olfr381 | BC030499 | Mir142hg |
| 2810021J22Rik | F930015N05Rik | Olfr382 | Slc13a2os | Gm11505 |
| Zfp39 | Gm12292 | Olfr385 | Gm11195 | Olfr462 |
| Gm12256 | 1700086D15Rik | Olfr384 | Fam58b | Olfr463 |
| Gm12257 | Gm12295 | Zfp735 | Gm11201 | Olfr464 |
| Hist3h2ba | Gm12298 | Olfr389 | Gm44787 | Gm11508 |
| Hist3h2a | 2310065F04Rik | Olfr390 | Gm9964 | 2010015M23Rik |
| Gm10435 | 9430073C21Rik | Olfr391-ps | AU040972 | Gm15892 |
| Gm15755 | Myhas | Olfr392 | Rab11fip4os1 | Gm38534 |
| Olfr222 | Gm12301 | Olfr393 | Rab11fip4os2 | Gm15893 |
| Olfr223 | C78197 | Olfr394 | Gm11205 | C030037D09Rik |
| Gm12264 | Ccdc42os | Olfr395 | 9130204K15Rik | Gm11493 |
| Gm16062 | 9330160F10Rik | Olfr23 | Gm26563 | 4930556N13Rik |
| Gm12714 | 9130213A22Rik | Olfr397 | Adap2os | Scpep1os |
| 1810063I02Rik | Cntrobos | Olfr398 | 4930507D10Rik | Gm11496 |
| Med9os | Kcnab3os | Olfr139 | Gm11417 | Dgkeos |
| 4930412M03Rik | Chd3os | Olfr399 | 4930527B05Rik | Gm11497 |

| | | | | |
|---|---|---|---|---|
| Gm2018 | Gm11562 | Gm20511 | 4732490B19Rik | Gm11767 |
| Gm11494 | Krtap4-1 | 1700023F06Rik | Fam104a | Gm11766 |
| Gm45716 | Gm11555 | Arhgap27os3 | D11Wsu47e | Gm11770 |
| 4930405D11Rik | Gm11563 | Arhgap27os2 | 1700025D23Rik | Gm11772 |
| Gm20390 | Gm14190 | Arhgap27os1 | 1700001J04Rik | Hgs.1 |
| Gm21885 | Gm14180 | Gm39397 | Gm11690 | Gm16755 |
| Gm11541 | Gm11595 | Gm884 | Gm11691 | Gm11788 |
| Hils1 | Gm11596 | Gm11659 | 4932435O22Rik | Gm11789 |
| Gm11513 | Gm11569 | Gm11665 | Dnaic2 | Gm17586 |
| Dlx4os | Gm11554 | Gm11639 | Slc9a3r1 | Myadml2os |
| Gm11515 | Krtap4-13 | Gm10842 | Ict1os | Notumos |
| Gm11520 | Gm11564 | 1700052K11Rik | Atp5h | Gm17178 |
| Gm11521 | 2300003K06Rik | 10-Mar | Gm11695 | Cbr2 |
| Gm29477 | Gm11568 | Gm11640 | Gm26613 | Gm11773 |
| Gm11528 | Gm11559 | Gm11651 | Smim6 | Gm28192 |
| Phb | Gm11567 | Gm11646 | Recql5os1 | Gm11775 |
| Gm26830 | Krtap31-1 | Taco1os | Sap30bpos | Gm11791 |
| Gm9796 | Gm11565 | Gm11672 | B230344G16Rik | Hexdc |
| Zfp652os | Krtap31-2 | Gm10840 | Gm26730 | BC017643 |
| B130006D01Rik | Gm11571 | Gm11706 | Gm11739 | Gm12589 |
| Gm11527 | Gm12347 | Gm11707 | Gm29292 | Gm29513 |
| Gm11534 | Krt42 | Gm11715 | 1810032O08Rik | Gm15643 |
| Gm29202 | Gm14206 | Gm11716 | St6galnac2.1 | Gm12590 |
| Atp5g1 | Gm12349 | Gm11712 | BC018473 | Gm46400 |
| Gm11535 | Ttc25 | Gm11713 | Gm11728 | Gm40653 |
| Gm53 | Dhx58os | C330019F10Rik | Gm11731 | Gm48530 |
| Hoxb5os | Gm11615 | Gm11657 | Gm11730 | 2810429I04Rik |
| Gm11536 | Gm28156 | Gm11655 | 9-Sep | Fam208b |
| Hoxb3os | G6pc | Cep112os1 | Gm16045 | Gm47695 |
| 2010300F17Rik | Gm27029 | Gm11670 | Gm11729 | Gm35190 |
| Gm11537 | Gm11626 | E030025P04Rik | Gm34418 | Gm47904 |
| Gm11525 | Gm11634 | 1700096J18Rik | Gm11733 | Gm47925 |
| Gm11583 | Gm11635 | Gm11696 | Gm11734 | Gm48010 |
| Gm11592 | Gm11551 | 9930022D16Rik | Gm11723 | Gm40658 |
| 4933428G20Rik | 4930417O22Rik | Gm15642 | Gm11724 | Gm5444 |
| Gm11614 | Gm20659 | Gm11685 | Gm20708 | Gm47450 |
| 1700001P01Rik | 1700006E09Rik | Gm11684 | Gm11725 | AC151530.1 |
| Gm11630 | E130111B04Rik | 1700012B07Rik | Gm11738 | AC133498.1 |
| Gm12352 | Gm11586 | Gm11683 | Gm11747 | 1700016G22Rik |
| Gm12355 | BC030867 | 1700023C21Rik | Gm11750 | Gm47507 |
| Gm12359 | Bloodlinc | Gm11697 | Gm26508 | Gm46401 |
| Gm11940 | Gm11627 | Gm11682 | Gm11754 | Gm47514 |
| A830036E02Rik | Gm11636 | Gm11674 | Gm11755 | Gm47548 |
| Gm11939 | 2810433D01Rik | Gm11680 | Gm26888 | Gm47549 |
| Gm11938 | Gm26668 | Gm11681 | Gm11752 | Gm35615 |
| Gm11937 | 2410004I01Rik | 4933434M16Rik | Rptoros | Gm28465 |

| | | | | |
|---|---|---|---|---|
| Gm40662 | Tcrg-V3 | Gm11290 | C230035I16Rik | Prl3d2 |
| CT010465.1 | Trgj3 | Hist1h2ah | Gm46404 | Prl3d3 |
| Gm36074 | Tcrg-C3 | Hist1h2bk | Gm11335 | Prl3c1 |
| Gm48257 | Tcrg-C2 | Hist1h4i | Hist1h4h | Prl3b1 |
| Gm9745 | Trgj2 | Hist1h2ag | Hist1h2af | Prl3a1 |
| Gm36264 | Tcrg-V1 | Hist1h2bj | Hist1h3g | 4930511O05Rik |
| Gm48825 | Trgj4 | Vmn1r188 | Hist1h2bh | Prl6a1 |
| Gm26601 | Tcrg-C4 | Vmn1r189 | Hist1h3f | Prl8a2 |
| Gm36423 | 9330199G10Rik | Vmn1r191 | Hist1h4f | Prl2b1 |
| Gm47802 | A530099J19Rik | Vmn1r192 | Hist1h1d | Prl8a6 |
| Gm26861 | Gm32036 | Vmn1r193 | Hist1h3e | Prl8a8 |
| Gm47407 | Olfr1370 | Vmn1r194 | Hist1h2ae | Prl8a9 |
| Gm36525 | Olfr1369-ps1 | Vmn1r195 | Hist1h2bg | Prl8a1 |
| Gm47408 | Olfr263 | Vmn1r196 | Hist1h2bf | Prl7b1 |
| Gm47486 | Olfr1368 | Vmn1r197 | Hist1h2ad | Prl7a1 |
| Gm30239 | Gm47806 | Vmn1r198 | Hist1h3d | Prl7a2 |
| Ero1lb | Olfr1367 | Vmn1r199 | Hist1h4d | Prl7d1 |
| Prl2c3 | Gm10065 | Vmn1r200 | Hist1h2be | Prl7c1 |
| Prl2c2 | Gm11273 | Vmn1r201 | Hist1h1e | Prl2a1 |
| Gm48544 | Olfr1366 | Vmn1r202 | Hist1h2ac | Prl2c1 |
| Prl2c5 | Olfr1535 | Vmn1r203 | Hist1h2bc | Prl4a1 |
| Gm48682 | Olfr1364 | Gm11300 | Hist1h1t | Prl5a1 |
| Gm47882 | Olfr1362 | Vmn1r204 | Hist1h4c | Gm11361 |
| Gm30893 | Olfr11 | Vmn1r205 | Hist1h1c | Gm40841 |
| Gm20043 | Olfr1361 | Vmn1r206 | Hist1h3c | Gm34639 |
| Gm26645 | Olfr1360 | Gm11309 | Hist1h2bb | Gm26735 |
| 4933412O06Rik | Olfr1359 | Vmn1r208 | Hist1h2ab | A330102I10Rik |
| A530046M15Rik | Hist1h2bl | Gm11314 | Hist1h3b | Gm11365 |
| Gm31450 | Hist1h2ai | Vmn1r209 | 4930558J22Rik | Gm11368 |
| Gm48444 | Hist1h3h | Vmn1r210 | Hist1h4b | 4930519D14Rik |
| Gm48491 | Hist1h2bm | Vmn1r211 | Hist1h4a | Gm29675 |
| Gm48492 | Hist1h4j | Vmn1r212 | Hist1h3a | Gm5447 |
| Gm48503 | Hist1h4k | Vmn1r213 | Hist1h1a | Gm48662 |
| Gm48754 | Hist1h2ak | Vmn1r214 | Gm11337 | 4930401O12Rik |
| Gm48799 | Hist1h2bn | Vmn1r215 | Hist1h2ba | Gm11373 |
| 4930448F12Rik | Gm11274 | Vmn1r216 | Hist1h2aa | G630018N14Rik |
| Gm48829 | Hist1h1b | Vmn1r216.1 | Gm11339 | Gm11376 |
| Gm47658 | Hist1h3i | Vmn1r217 | Gm11342 | A530084C06Rik |
| Gm47661 | Hist1h2an | Vmn1r218 | C530050E15Rik | Gm11377 |
| Gm31887 | Hist1h2bp | Vmn1r219 | 1700016G14Rik | 1700018A04Rik |
| Tcrg-V7 | Hist1h2bq | Vmn1r220 | 4932702P03Rik | Gm11378 |
| Tcrg-V4 | Hist1h2ao | Vmn1r222 | Gm11351 | Gm11379 |
| Tcrg-V6 | Hist1h4m | Vmn1r223 | 1700092E19Rik | Gm35732 |
| Tcrg-V5 | Hist1h4n | 4930557F10Rik | 4921520N01Rik | Gm11381 |
| Trgj1 | Hist1h2ap | 4933404K08Rik | Gm47888 | Gm48073 |
| Tcrg-C1 | Hist1h2br | 4930586N03Rik | Prl3d1 | D930007J09Rik |

| | | | | |
|---|---|---|---|---|
| Gm47662 | Gm47732 | Gm48570 | 4931429P17Rik | Gm16249 |
| Gm36099 | Gm47734 | Gm32401 | Gm36101 | Gm29431 |
| Gm11397 | Gm29590 | Gm32184 | A330033J07Rik | 5330429C05Rik |
| Gm6093 | Gm30918 | Gm5082 | Gm36346 | 4930526F13Rik |
| Gm47916 | Gm47751 | 1700061E18Rik | Gm49291 | Gm17617 |
| 1110046J04Rik | Gm47754 | Gm28564 | A330048O09Rik | Gm46416 |
| Gm47076 | Gm40915 | Gm28707 | Gm48648 | Gm15911 |
| Gm36500 | CT009713.1 | Gm47118 | A830005F24Rik | 4930451E10Rik |
| Gm40909 | AI463229 | Gm15810 | 6720427I07Rik | Gm28760 |
| Gm47127 | Gm49350 | Gm15813 | Gm36550 | Gm47071 |
| Gm15908 | Gm40918 | Gm47683 | Phf2os1 | H2afy |
| Gm47150 | Gm31600 | Gm32939 | Fam120aos | 4930550C17Rik |
| Gm47151 | Gm46411 | Gm47728 | Iars | Gm10782 |
| Gm47152 | Gm47990 | Gm20751 | Gm30302 | Gm3045 |
| Gm47157 | 4930579J19Rik | Gm33115 | Gm47977 | 2010203P06Rik |
| Gm36839 | Gm31834 | Gm33195 | Gm48051 | Gm45623 |
| 4933417A18Rik | Gm48056 | Gm2233 | 4933433N18Rik | Gm48441 |
| Gm16984 | Gm40922 | Gm33489 | Gm906 | Gm26555 |
| Gm48626 | Gm47400 | Gm33630 | Gm31126 | 2210016F16Rik |
| Gm40910 | Gm46392 | Gm5083 | Gm8739 | Gm47918 |
| Gm48629 | Gm47510 | Gm33684 | Fbxw17 | 4930455J16Rik |
| 4930529N20Rik | Gm47511 | A330076C08Rik | Gm47429 | Gm47947 |
| 1700019C18Rik | Gm47509 | Gm47781 | Gm904 | Gm40968 |
| Gm48694 | Gm26514 | Gm29676 | Gm8765 | Gm47423 |
| Gm48703 | 5033403F01Rik | Gm33958 | 4930518P08Rik | 4930415C11Rik |
| Gm48704 | Gm47348 | Gm27007 | Gm32834 | Gm34307 |
| Gm48707 | Gm3509 | 1700029N11Rik | Gm26651 | Gm34354 |
| Gm48708 | Gm47349 | Gm34084 | Gm48190 | Gm47359 |
| Gm48746 | Gm47351 | Gm47792 | Gm48199 | Gm34558 |
| Gm30127 | Gm47352 | Gm34276 | BB123696 | Gm34672 |
| Gm48770 | Gm32243 | Gm47805 | Gm33424 | Gm47360 |
| Gm48765 | Gm40923 | Gm9817 | Gm48336 | Gm34721 |
| Gm48767 | Gm47316 | Gm40932 | Gm2762 | Gm34788 |
| Gm30177 | Gm47039 | Gm47448 | Gm43262 | 4930455M05Rik |
| Gm49144 | Gm9979 | Gm34466 | AC154808.1 | Gm46419 |
| Gm47675 | Gm47061 | Gm47460 | 9530014B07Rik | 4930528D03Rik |
| Gm30489 | Gm26688 | 4930453C13Rik | 4930555G21Rik | 1700014D04Rik |
| Gm47707 | Gm47067 | Gm45949 | Gm34278 | Gm49354 |
| Gm30600 | Gm48107 | Gm10113 | Gm48548 | Etohd2 |
| Gm47709 | Gm31683 | Gm47523 | Gm34557 | Zcchc6 |
| Gm47711 | A730081D07Rik | Gm48250 | Gm48550 | Gm34961 |
| Gm4035 | Gm48510 | A930002C04Rik | Gm16578 | Gm48384 |
| Gm47730 | Gm32063 | Gm48612 | Gm48615 | Gm19866 |
| Gm29459 | Gm17364 | Gm35733 | Gm48623 | A530065N20Rik |
| Gm29458 | Gm26877 | 4930471G24Rik | Gm48622 | Gm48396 |
| Gm47731 | Gm48571 | G630093K05Rik | Gm16248 | Gm48397 |

| | | | | |
|---|---|---|---|---|
| Gm48488 | Gm47194 | Gm48900 | Eprn | Gm20379 |
| Gm5084 | Olfr465-ps1 | Zfp459 | Slc6a19os | C030017D09Rik |
| Gm35333 | Olfr466 | Zfp874b | Gm41002 | Gm4814 |
| Gm48500 | Gm47249 | Gm48899 | Zfp72 | A630019I02Rik |
| A530001N23Rik | Gm36445 | Gm26965 | Ftl1-ps1 | Papd4 |
| Ctsll3 | Gm47251 | Zfp748 | Gm47428 | Gm15622 |
| 4930486L24Rik | Gm10775 | 9430065F17Rik | E430024I08Rik | AW495222 |
| Gm49392 | Gm7762 | Gm49345 | CT009718.2 | Gm26527 |
| Gm49393 | Gm40983 | Gm48095 | CT009718.1 | Gm47216 |
| Gm49391 | Gm48168 | Zfp65 | Gm47467 | Gm15907 |
| Ctla2b | Gm48166 | Zfp493 | Gm47469 | Gm32089 |
| Tpbpb | Gm10139 | 4930525G20Rik | Gm46430 | Gm48102 |
| Ctla2a | Gm26639 | Gm10037 | 1700037F03Rik | Gm48287 |
| Tpbpa | Gm48170 | BC048507 | Ttc37 | Gm32305 |
| Gm40975 | Platr2 | Gm7969 | Gm31219 | Gm32351 |
| Ctsq | Gm40987 | Gm48419 | Fam172a | Gm48350 |
| Ctsr | Gm48221 | Gm48423 | Gm38604 | Gm48420 |
| Cts6 | Gm48222 | Gm8016 | 3110006O06Rik | Gm16243 |
| Gm49398 | Gm10324 | Gm48436 | A830082K12Rik | Gm48730 |
| Cts7 | Gm48223 | Gm48556 | Gm48399 | Gm20075 |
| Gm49352 | Gm17514 | 1700001L19Rik | Gm31946 | Gm17190 |
| Gm49357 | Gm10772 | Gm35161 | Gm32067 | Gm29543 |
| Gm48116 | Gm26754 | Gm26844 | Gm48402 | Gm17622 |
| Gm48228 | Gm26715 | Gm35514 | Gm29318 | Col4a3bp |
| Gm40977 | 2410141K09Rik | Papd7 | Gm28526 | Gm48133 |
| Gm19792 | Gm26806 | A530095I07Rik | 5430425K12Rik | Gm48597 |
| Gm49359 | Gm40988 | Gm35618 | Gm4211 | 1700029F12Rik |
| 6720489N17Rik | Gm48412 | Gm48819 | 9330111N05Rik | Gm6169 |
| Platr25 | Gm10767 | 4930547H16Rik | Gm49375 | C430039J16Rik |
| Gm48795 | 4933433G19Rik | Gm26819 | Gm17259 | Gm33447 |
| Gm48812 | Gm46440 | 4933416O17Rik | Gm48155 | Gm2379 |
| 2010111I01Rik | Gm28557 | 1700084F23Rik | 2310067P03Rik | Gm41030 |
| Gm16907 | Rslcan18 | Gm3772 | C130071C03Rik | Gm5086 |
| Gm16133 | Zfp759 | AU017674 | Gm26803 | 5330416C01Rik |
| Gm30655 | Gm48732 | 4930520P13Rik | 2810049E08Rik | Gm26619 |
| Gm30709 | Rsl1 | 1700100L14Rik | Gm4241 | Gm10260 |
| 1700024I08Rik | Zfp455 | D030007L05Rik | Gm17750 | Gm41031 |
| Gm47390 | Gm49064 | Gm36377 | A230107N01Rik | Gm34388 |
| Gm47418 | F630042J09Rik | Gm36426 | Gm34585 | Gm21976 |
| 1810034E14Rik | Gm48824 | Gm36607 | Gm4117 | 2310005E17Rik |
| Aaed1 | Gm28044 | Gm36529 | Gm29680 | Gm29501 |
| Gm47003 | Zfp953 | 8030423J24Rik | Gm47520 | Gm10320 |
| Gm47123 | Gm28041 | 1700112M02Rik | Gm47381 | Gm9465 |
| 1700015C15Rik | Gm17039 | Gm20554 | A830009L08Rik | Gm47551 |
| Gm31218 | Zfp456 | Gm47902 | 4833422C13Rik | Gm35215 |
| Gm47193 | Zfp429 | D730050B12Rik | 1700119I11Rik | 1700024P04Rik |

| | | | | |
|---|---|---|---|---|
| Gm35279 | Gm15326 | Gm36161 | Gm47833 | AC164424.1 |
| A930014D07Rik | Gm15322 | Gm36079 | Gm34237 | AC153140.2 |
| 2310020H05Rik | Gm15323 | Gm21188 | Fam49a | Gm4419 |
| Gm807 | Gm48802 | Gm20767 | 4921511I17Rik | Gm4425 |
| BC001981 | Gm48837 | Gm21818 | Platr19 | Gm47997 |
| Gm48596 | 2810403G07Rik | Gm21762 | Gm48187 | 2410018L13Rik |
| Gm29502 | Gm48876 | Gm21731 | 4930519A11Rik | Gm10330 |
| 5930438M14Rik | Gm48879 | AF067061 | Gm40271 | 9030624G23Rik |
| 4932411K12Rik | Gm47827 | BC147527 | Gm35208 | Gm48896 |
| Gm47533 | Gm47850 | Tcstv3 | Gm48209 | Gm47701 |
| Gm29341 | Skiv2l2 | Rab10os | Gm35298 | Gm17746 |
| Gm47007 | AC159207.1 | Gm26520 | Gm48213 | Gm36287 |
| Gm29927 | 1700084D21Rik | Gm48512 | Gm48311 | Gm47705 |
| 1700099I09Rik | BC067074 | 1110002L01Rik | Gm35725 | Gm47713 |
| Gm17160 | Gm41071 | Dtnbos | Gm35890 | AC163354.1 |
| Gm30551 | 4921509O07Rik | Dnmt3aos | Gm48479 | Gm47733 |
| Gm47849 | Gm34471 | Gm48001 | Gm48480 | Gm29687 |
| Gm47851 | Gm34586 | 4921501I09Rik | Fam84a | 4930480M12Rik |
| 2610204G07Rik | A430090L17Rik | 2900045O20Rik | Gm16497 | Gm29968 |
| Fam159b | Gm47040 | Gm31938 | 4930448C13Rik | 4930549C15Rik |
| 4933425L06Rik | 4930467J12Rik | Gm46332 | Gm48539 | 1700020D12Rik |
| Gm10739 | 4930544M13Rik | Gm48610 | Gm48558 | Gm47847 |
| Gm30411 | Gm49395 | Gm3625 | Gm48584 | 4933409F18Rik |
| Gm28989 | Gm10734 | Gm48678 | Gm36235 | Gm9866 |
| Gm28988 | Gm47914 | 2810032G03Rik | Gm48607 | Gm4166 |
| Gm48684 | Gm47913 | AC159282.1 | Gm36372 | Gm45941 |
| Gm31452 | 4930435F18Rik | Gm48619 | Gm48606 | Gm31025 |
| AI197445 | Gm6416 | 1700101O22Rik | Gm48605 | Gm47872 |
| Gm32090 | Gm17509 | Gm32828 | Gm48140 | Gm47871 |
| 1700006H21Rik | 4933413L06Rik | Gm48633 | Gm48227 | Gm31508 |
| B230220B15Rik | B430218F22Rik | Gm33037 | Gm36495 | 4833405L11Rik |
| AC139580.1 | 1700003P14Rik | Gm48898 | Pqlc3 | Gm31333 |
| Gm33045 | Gm10732 | Gm48071 | 2410004P03Rik | C630031E19Rik |
| Gm33172 | Gm16263 | Gm48075 | Gm48538 | Gm15691 |
| 4930526H09Rik | Gm47336 | 5033421B08Rik | Gm36752 | 2310016D03Rik |
| Gm38397 | Nnt.1 | 9930038B18Rik | Gm48313 | Gm28806 |
| 3110015C05Rik | 3110070M22Rik | Gm47391 | AC124772.1 | Gm32443 |
| Gm6270 | Gm48265 | Gm46323 | Gm40849 | 6030469F06Rik |
| AC118475.1 | 1700074H08Rik | Gm48762 | Gm10479 | Gm29542 |
| Gm15290 | Gm48342 | Gm48764 | Gm5784 | Gm48809 |
| Gm15288 | Gm41077 | Gm48790 | AC124739.1 | Gm33111 |
| Gm10198 | AF067063 | Gm48791 | AC124739.2 | 5430401H09Rik |
| Gm15287 | D13Ertd608e | Gm38407 | Gm49371 | Gm33308 |
| Gm15325 | Tcstv1 | 7420701I03Rik | Gm21863 | AC155255.1 |
| AC118704.1 | Gm21761 | 9530020I12Rik | AC163633.2 | Gdap10 |
| Gm15324 | B020031M17Rik | 4930511A02Rik | AC163633.1 | Gm47948 |

| | | | | |
|---|---|---|---|---|
| 4933406C10Rik | Gm33680 | Gm48421 | Gm48823 | Pcnx |
| Sypl | Gm48522 | Gm48422 | Gm15283 | AC124484.1 |
| Gm16267 | Gm48535 | Gm48559 | Gm49321 | Gm48242 |
| Atxn7l1os1 | B230217J21Rik | 4930471E15Rik | Gm29587 | 1700085C21Rik |
| Atxn7l1os2 | Gm48578 | Gm31063 | Dbpht2 | Gm29530 |
| Twistnb | Gm34304 | Gm47979 | Gm34552 | Gm26623 |
| Gm48236 | Gm43517 | Gm47432 | Gm11042 | Gm29361 |
| Gm48237 | 3110039M20Rik | Gm47453 | Gm39473 | Gm47666 |
| Gm34047 | 1810007C17Rik | Gm47456 | 4930442G10Rik | Gm26571 |
| 9130015A21Rik | Gm40418 | Gm47489 | Gm47689 | Gm49366 |
| Gm40392 | Gm48779 | Gm47515 | Gm34868 | 4732463B04Rik |
| Gm34215 | Gm40421 | Gm2912 | 4930426I24Rik | Elmsan1 |
| D630036H23Rik | 1700008C04Rik | Gm47518 | Gm10451 | Gm31513 |
| Ispd | Gm26517 | Gm31447 | AI463170 | Gm48573 |
| Gm40394 | Gm47431 | Gm49383 | Gm35041 | D030025P21Rik |
| Gm48613 | Gm35135 | Gm49384 | Gm35189 | Gm48709 |
| Gm48616 | Gm35188 | Gm47545 | Gm35240 | Gm17139 |
| Gm29007 | 1700031P21Rik | Gm15561 | 4930458K08Rik | Prox2os |
| Gm34408 | Gm35239 | Gm9887 | Gm6657 | Gm47819 |
| Gm48715 | 1700030L22Rik | Atp5s | Fam71d | Gm40477 |
| 4930428E07Rik | 1700060O08Rik | 4931403G20Rik | Mpp5 | Gm32296 |
| Gm34611 | Gm35818 | F730035M05Rik | 9230116L04Rik | Gm26531 |
| Gm34662 | 1700104L18Rik | Gm48747 | Gm48780 | Gm805 |
| Gm34809 | Gm47552 | Gm48847 | 9430078K24Rik | 4732487G21Rik |
| Gm47855 | Gm7550 | Gm32219 | Gm36660 | Gm26698 |
| Gm34923 | Gm46328 | Gm24474 | Gm47752 | Gm6566 |
| Gm47859 | Gm10465 | Gm48866 | Gm26669 | Gm26764 |
| Gm47868 | Gm40884 | Gm32369 | 1300014J16Rik | Gm29362 |
| Arl4aos | Gm20403 | Gm40437 | Gm47767 | Gm5788 |
| Gm17056 | 2700097O09Rik | Gm46355 | Gm47766 | Gm8300 |
| Gm7008 | 1700047I17Rik2 | Gm10457 | Gm30025 | Oog1 |
| Gm47375 | 1110008L16Rik | Gm33016 | 2310002D06Rik | Gm4027 |
| Gm47376 | Aldoart2 | Gm40438 | 4933406B15Rik | Gm16381 |
| Gm47373 | Gm47682 | 1700083H02Rik | Gm47879 | Gm21319 |
| Gm47368 | Gm19990 | 4930404H11Rik | Plekhd1os | Gm21936 |
| Gm47371 | Gm26973 | 9630002D21Rik | Gm26796 | BB287469 |
| Gm46348 | C87198 | Gm4756 | Gm26545 | Gm2001 |
| Gm47013 | Gm15524 | Gm33785 | 1700052I22Rik | Gm2016 |
| Gm47020 | Gm16246 | Gm26709 | Gm3693 | Gm2022 |
| AC113204.1 | 4921518K17Rik | Gm48653 | Gm20498 | Gm2035 |
| Gm47030 | BC042761 | D830013O20Rik | Gm4787 | Gm2042 |
| Gm10165 | Gm46329 | Gm33929 | Adam4 | Gm2075 |
| Gm46349 | Gm47645 | Gm48656 | Gm28370 | Gm6803 |
| Gm47096 | Gm17529 | Gm34016 | Gm16572 | Gm2056 |
| Gm48508 | Gm48301 | 2210039B01Rik | AC125351.1 | Gm16368 |
| Gm9921 | Gm20063 | Gm8075 | Gm47080 | Gm10436 |

| | | | | |
|---|---|---|---|---|
| Gm8332 | Gm28051 | B830012L14Rik | Ighd5-3 | Ighv16-1 |
| Gm5662 | Gm21971 | Mirg | Ighd2-5 | Ighv6-2 |
| Gm5039 | Gm20604 | 4930511J24Rik | Gm37327 | Ighv12-1 |
| 3200001D21Rik | Tmem251 | Gm34667 | Ighd5-2 | Ighv12-2 |
| 4930473H19Rik | Gm47167 | Gm34719 | Ighd2-4 | Ighv15-1 |
| Gm48665 | Gm29508 | 3110009F21Rik | Ighd2-3 | Ighv13-1 |
| Gm48664 | Gm15523 | Gm34785 | Ighd1-1 | Ighv3-7 |
| 1700040E09Rik | 9330161L09Rik | Gm40576 | Adam6b | Ighv5-21 |
| Gm2270 | Gm47267 | AC152827.1 | Ighd3-1 | Ighv8-1 |
| Gm47684 | Serpina16 | Gm35558 | Gm16968 | Ighv1-1 |
| Gm40538 | B430119L08Rik | Gm35558.1 | Ighv5-1 | Ighv6-4 |
| 4930544I03Rik | AC122556.1 | Gm17111 | Ighv2-1 | Ighv1-2 |
| 1700105G05Rik | 4930408O17Rik | 1700001K19Rik | Ighv5-2 | Ighv1-3 |
| 5430427M07Rik | Gm28875 | Gm26912 | Ighv2-2 | Ighv10-2 |
| Gm47804 | Gm47648 | 6030440G07Rik | Ighv5-3 | Ighv1-6 |
| Gm16876 | Gm2721 | 4930595D18Rik | Ighv6-1 | Ighv10-4 |
| Gm40548 | Gm46376 | A230087F16Rik | Ighv2-3 | Ighv1-8 |
| 4930559C10Rik | AU015791 | Gm10425 | Ighv5-5 | Ighv15-2 |
| Gm47396 | Gm47796 | Gm40578 | Ighv5-7 | Ighv1-10 |
| Gm47397 | Gm48807 | Gm266 | Ighv2-4 | Ighv1-13 |
| Gm40552 | Gm19554 | Gm36635 | Ighv5-8 | Ighv1-13.1 |
| Gm35274 | AC163040.1 | Apopt1 | Ighv5-8.1 | Ighv1-14 |
| Gm35326 | Gm32635 | Gm15996 | Ighv5-10 | Ighv1-17-1 |
| Gm47415 | Gm47646 | 5033406O09Rik | Ighv2-5 | Ighv1-17 |
| Gm47439 | Gm17032 | Gm36757 | Ighv5-11 | Ighv1-19-1 |
| Gm47566 | 1700121N20Rik | 2010107E04Rik | Ighv2-6 | Ighv1-21-1 |
| Gm40893 | Gm17033 | E330035G20Rik | Gm37976 | Ighv1-21 |
| Gm47109 | 4933406K04Rik | B020018J22Rik | Gm37418 | Ighv1-25 |
| Gm47116 | 1700013N06Rik | Adssl1 | Gm38203 | Ighv1-27 |
| 4930474N09Rik | Gm16087 | BC022687 | Gm38205 | Ighv1-28 |
| 3300002A11Rik | Gm16086 | 1700127F24Rik | Ighv2-9-1 | Ighv1-29 |
| Gm47177 | Gm16085 | Gm26583 | Gm37722 | Ighv1-30 |
| 1700064M15Rik | Gm16084 | 9230104M06Rik | Ighv5-13 | Ighv1-32 |
| Gm19951 | Gm2800 | Gm37944 | Ighv2-6-8 | Ighv1-33 |
| Gm47207 | Gm3234 | Ighj4 | Gm7003 | Ighv1-35 |
| 4930477G07Rik | 3110018I06Rik | Ighj3 | Ighv2-7 | Ighv1-38 |
| Gm26723 | 4930465M20Rik | Ighd4-1 | Ighv5-18 | Ighv1-40 |
| Gm40557 | Gm15208 | Ighd3-2 | Ighv2-8 | Ighv1-41 |
| Gm48383 | 4930478K11Rik | Ighd5-6 | Ighv2-9 | Ighv1-44 |
| A630072L19Rik | Gm16596 | Ighd2-8 | Ighv5-19 | Ighv1-45 |
| Gm36756 | Gm33467 | Ighd5-5 | Ighv4-1 | Ighv1-46 |
| Gm47639 | Gm33682 | Ighd2-7 | Gm37961 | Ighv8-3 |
| D130020L05Rik | Wars | Ighd5-8 | Ighv11-1 | Ighv1-48 |
| Gm16339 | Gm34081 | Ighd5-4 | Ighv4-2 | Ighv1-51 |
| Gm30198 | Gm26906 | Ighd2-6 | Ighv3-2 | Ighv8-7 |
| Gm20069 | Gm26945 | Ighd5-7 | Ighv11-2 | Gm30948 |

| | | | | |
|---|---|---|---|---|
| Ighv1-57 | Gm19276 | Gm41289 | 9330182O14Rik | Gm48946 |
| Ighv1-60 | Tars | Gm32618 | Gm33301 | Fam84b |
| Ighv1-62 | Gm49106 | Gm32764 | Gm41311 | 4930402D18Rik |
| Ighv1-62-1 | Gm49107 | 4930592A05Rik | Gm16294 | AC140304.2 |
| Gm19331 | 1700047G03Rik | Gm41290 | Gm26854 | AC140304.1 |
| Gm37511 | Gm34759 | 9430069I07Rik | AC136372.1 | AC124097.1 |
| Gm38184 | Gm49240 | 4930413F20Rik | AC158972.2 | AC160335.1 |
| Ighv8-10 | 1810049J17Rik | Gm33497 | AC158129.1 | AC125455.1 |
| Ighv1-65 | Gm49113 | BC048602 | Gm17473 | AC134453.1 |
| Ighv1-68 | Gm49116 | Gm48960 | Gm49271 | Gm30159 |
| Ighv1-70 | 4930557F08Rik | Gm41293 | Gm10373 | Gm41335 |
| Ighv1-73 | Gm41276 | Gm34150 | 1700022A22Rik | Gm49014 |
| Ighv8-14 | Gm49126 | Gm34093 | 4930523O13Rik | 4930449C09Rik |
| Ighv8-15 | Gm49127 | Gm49224 | 4930548G14Rik | Gm41336 |
| Ighv8-16 | Gm49128 | Gm49262 | Gm41318 | Gm49019 |
| Gm42643 | Gm41277 | Gm46515 | Gm49198 | Fam49b |
| Ighv1-79 | C030047K22Rik | Gm10385 | Gm48913 | Gm20717 |
| Gm42990 | Gm35496 | Gm49263 | Gm19303 | 1700010G06Rik |
| Ighv1-83 | Gm49160 | Gm49282 | Gm34562 | Gm30563 |
| Ighv1-86 | Gm49162 | Gm10384 | Gm34678 | Gm30691 |
| Zfp386 | Gm49166 | Gm41300 | Gm48923 | Gm21798 |
| Gm20658 | AC124561.1 | Gm34590 | Gm41322 | Gm21961 |
| Wdr60 | Gm49191 | Gm26766 | Gm34794 | Gm27153 |
| Gm11027 | AC131068.1 | Gm16136 | 1700015H07Rik | Gm27242 |
| Gm48079 | Gm35769 | Gm16137 | Gm7489 | Gm30929 |
| Gm5441 | Gm2824 | Gm28221 | Nov | Lrrc6 |
| AC163032.1 | Gm35996 | Gm15941 | AC137950.1 | Gm17140 |
| D230030E09Rik | 9230109A22Rik | 4930447A16Rik | Gm41325 | Wisp1 |
| Gm48681 | 4930445E18Rik | Gm15942 | 1700040F17Rik | Gm17035 |
| Gm49231 | Gm5468 | Gm35019 | Gm46516 | AC087116.1 |
| AW549877 | Gm48956 | Gm49085 | Gm49211 | 1700012I11Rik |
| Gm2093 | Gm36642 | Gm35167 | Gm49212 | Gm20732 |
| Gm15632 | Gm41279 | Gm49313 | Gm2582 | AC141893.2 |
| Gm15938 | 11-Mar | Gm35248 | Slc22a22 | Gm7125 |
| Gm16311 | Gm49267 | Gm26863 | Gm16006 | AC122275.2 |
| Fyb | Gm36899 | Gm41307 | 9330154K18Rik | AC127697.2 |
| Gm2245 | Fam105a | G930009F23Rik | Gm29394 | Gm16308 |
| Gm16029 | Gm31458 | Gm49077 | Wdyhv1 | AC100400.1 |
| Gm49207 | Gm49233 | 2310043O21Rik | Gm15943 | AC118008.1 |
| 2410089E03Rik | 4930570B17Rik | 1100001I12Rik | Gm2675 | 1700085D07Rik |
| Gm49247 | 9630009A06Rik | Gm45924 | Gm49356 | AC116769.1 |
| Gm31282 | 4930430F21Rik | Gm49097 | 4930544F09Rik | 4933427E11Rik |
| Gm49249 | 6-Mar | AC164883.2 | Gm36617 | D730001G18Rik |
| Gm2310 | Fam173b | Gm16291 | AC161172.1 | Gm17189 |
| Gm21973 | 4930518I17Rik | AU022793 | Gm19510 | 2010109I03Rik |
| Gm10389 | 4930465K09Rik | Gm33251 | Gm2682 | AC139671.1 |

| | | | | |
|---|---|---|---|---|
| Ly6i | Gm16059 | AC142474.1 | Gm44579 | AC163018.1 |
| Ly6a | Gm26884 | AC142474.2 | Olfr287 | AC157583.2 |
| Gm28068 | Gm16575 | AC153010.1 | Olfr286 | AC103674.1 |
| Ly6c1 | Npcd | AC124455.1 | H1fnt | AC103674.4 |
| Gm28502 | AC113595.2 | Fam19a5 | Olfr285 | AC120787.5 |
| Ly6c2 | AC113595.1 | AC124691.1 | Olfr284 | AC139844.1 |
| Ly6g | AC161199.1 | AC157944.1 | Olfr283 | Gm9918 |
| BC025446 | AC140267.1 | AC147230.1 | Olfr282 | Gm26518 |
| Ly6f | AC123059.1 | AC147230.2 | Olfr281 | Gm10337 |
| AC119264.1 | AC147041.1 | Zdhhc25 | OR5BS1P | Gm28047 |
| Gm15945 | AC141880.1 | 1810021B22Rik | Olfr279 | AC123870.1 |
| Pycrl | Mkl1 | Gm26798 | 9330020H09Rik | Atp5g2 |
| Tsta3 | AC102334.1 | Odf3b | 4930415O20Rik | AC124532.2 |
| K230010J24Rik | Gm17025 | AC137513.1 | Gm29331 | Gm28876 |
| Gm48952 | Gm17597 | C730034F03Rik | AC156543.1 | Gm10830 |
| AC116487.2 | AC102103.1 | C230037L18Rik | B130046B21Rik | Gm28265 |
| BC024139 | 1700029P11Rik | Gm15609 | AC161165.6 | D930007P13Rik |
| AC110211.1 | AC104325.1 | AC119932.1 | AC161165.2 | AC162693.1 |
| Gm35339 | 1500009C09Rik | Alg10b | AC161165.4 | AC164069.1 |
| AC157566.4 | 3-Sep | AC158922.1 | AC161165.3 | Gtsf2 |
| Cyhr1 | Fam109b | CN725425 | AC157610.1 | Gm49272 |
| AC157554.2 | AC118710.3 | 4933438A12Rik | Nckap5los | Mucl1 |
| AC157554.4 | Gm20324 | Smgc | Gm17349 | Mucl2 |
| AC157554.3 | Gm29019 | Gm26760 | AC134548.2 | AC131776.4 |
| 1700109K24Rik | AW121686 | AC101921.1 | Gm16537 | AC191865.2 |
| 1700025B11Rik | Gm17206 | AC140403.3 | Gm17058 | AC127687.1 |
| AL592187.3 | AL591952.3 | Gm30085 | Gm17057 | AC127687.2 |
| AL592187.4 | AL591952.1 | AC109617.1 | Gm21917 | Olfr161 |
| Gm17638 | 1700001L05Rik | AC109617.2 | 2310068J16Rik | AC139347.1 |
| A730060N03Rik | AL603867.1 | AC121517.2 | AC133868.2 | Olfr15 |
| AL589692.1 | AL626769.1 | AC121517.1 | AC125526.1 | Gm26675 |
| Tex33 | 1810041L15Rik | AC102910.1 | Mettl7a1 | Gm20695 |
| AL590144.1 | AL611986.1 | Gm49169 | Mettl7a3 | Gm15537 |
| AL590144.3 | AL611987.1 | Gm6961 | Methig1 | Gm15879 |
| AL590144.2 | AL513352.1 | A130051J06Rik | Mettl7a2 | Gm15859 |
| AL591946.1 | Gm29666 | Gm49173 | Gm5475 | Gm15835 |
| Gm26634 | AL583891.1 | Gm17546 | Gm27209 | Gm16861 |
| AL592169.1 | 7530416G11Rik | D030018L15Rik | AC123724.1 | 12-Sep |
| 1700027A07Rik | AU022754 | 2610037D02Rik | Galnt6os | Gm42477 |
| AL589670.5 | AC162302.1 | AC158769.2 | AC113587.1 | Gm15983 |
| AL589670.1 | Gm15722 | AC158769.1 | Gm47841 | AC124490.1 |
| AL589670.3 | Gm15569 | AC158554.1 | Gm16031 | AC136518.1 |
| Gcat.1 | AC146911.1 | Rapgef3os1 | A330009N23Rik | Gm5767 |
| AL589670.4 | AC158974.1 | Rapgef3os2 | AC161812.1 | 1810013L24Rik |
| Pick1.1 | AC158974.2 | Gm26513 | Grasp | AC156026.1 |
| Gm20420 | AC139637.2 | Olfr288 | 6030408B16Rik | AC165274.3 |

| | | | | |
|---|---|---|---|---|
| AC136987.2 | Dgcr14 | Gm37419 | AC110166.2 | 4930404A05Rik |
| AC136987.1 | Vpreb2 | AC158397.1 | 1600019K03Rik | AC122551.1 |
| AC154256.1 | AC087064.2 | AC169509.5 | Dirc2 | Zbtb11os1 |
| AC154311.1 | 4933432I09Rik | AC169509.3 | Gm26838 | Gm15839 |
| Gm15558 | AC118542.1 | Lppos | Gm15564 | Gm16892 |
| CT010583.1 | AC084822.2 | A230028O05Rik | Ccdc58 | Gm26800 |
| Gm11172 | Rtl10 | Tprg | Gm5483 | CT025521.3 |
| Gm26822 | 4930588K23Rik | Gm4524 | Gm5416 | CT025521.2 |
| Gm21859 | AC133488.1 | Gm20319 | 2010005H15Rik | AC156023.3 |
| Gm4262 | 5-Sep | AC154512.1 | BC117090 | AC156023.2 |
| AC164093.2 | Gm28539 | AC154786.1 | BC100530 | AC172892.1 |
| 4930509G22Rik | 2010309G21Rik | Hrasls | Gm15845 | 4930461C15Rik |
| 2610020C07Rik | Iglj1 | AC154438.1 | AC117662.4 | Gm813 |
| Gm4279 | Iglj3p | Gm26834 | 4930565N06Rik | E330017A01Rik |
| AC154509.1 | Iglj3 | CT025592.1 | AC154762.2 | AC159200.1 |
| Gm9961 | Iglj4 | Gm26569 | Gm36028 | Olfr172 |
| AC117197.3 | Iglj2 | Gm1968 | Maats1 | Olfr173 |
| AC117197.2 | Gm43388 | 4632428C04Rik | Maats1os | Olfr175-ps1 |
| AC117197.1 | Iglv2 | 9030404E10Rik | Gm21987 | Olfr177 |
| 1700003L19Rik | Olfr164 | 1700025H01Rik | D930030I03Rik | Olfr178 |
| Mkl2 | Olfr165 | Gm26562 | Gm17103 | Olfr180 |
| Gm15738 | Olfr166 | CT009704.1 | 4930435E12Rik | Olfr181 |
| 2310015D24Rik | Olfr167 | AC163720.1 | Gm15802 | Olfr183 |
| 3110001I22Rik | Olfr168 | AC163720.3 | AC091463.1 | Olfr186 |
| Pla2g10os | Olfr169 | AC163720.2 | AC161816.1 | Olfr187 |
| AC154607.1 | Olfr170 | AC126280.1 | Gm28750 | Olfr190 |
| 2900011O08Rik | Olfr171 | AC130815.3 | 4932412D23Rik | Olfr191 |
| Fopnl | A930003A15Rik | AC130815.2 | AC161607.1 | Olfr193 |
| Gm15868 | AC164558.1 | AC125371.4 | Gm9968 | Olfr194 |
| A630010A05Rik | AC120150.1 | AC125371.3 | Gm15713 | Olfr195 |
| AC154606.1 | Gm16618 | Gm15743 | AC154408.2 | Olfr196 |
| Efcab1 | AC087898.2 | AC125371.2 | Gm26732 | Olfr198 |
| Gm21897 | AC087898.3 | AC161376.1 | Gm609 | Olfr199 |
| AC112943.2 | Gm49333 | AC161376.2 | Gm17783 | Olfr201 |
| Olfr19 | CT010490.3 | AC126055.1 | Gm15591 | Olfr202 |
| 4933404G15Rik | AC158985.3 | 0610012G03Rik | Gm15640 | Olfr203 |
| Vpreb1 | AC123977.1 | Gm15729 | Gm15638 | Olfr204 |
| AC166832.3 | Gm16863 | Tctex1d2 | Gm4737 | Olfr205 |
| 1700056N10Rik | 1300002E11Rik | AC087556.1 | CT025584.1 | Olfr206 |
| AC154667.1 | AC114990.2 | AC140186.1 | C330027C09Rik | Olfr209 |
| 2610318N02Rik | Gm26744 | Tnk2os | 1700026J12Rik | 1700022E09Rik |
| Gm26635 | 2610020F03Rik | Gm26769 | Gm15518 | Gm9017 |
| Tmem191c | 9230117E06Rik | Gm15657 | G730013B05Rik | AC154292.1 |
| AC078895.1 | Gm15651 | Gm15829 | AC138306.1 | AC163285.1 |
| Car15 | CT027991.1 | 1700119H24Rik | 1700116B05Rik | CT025155.1 |
| Gm20518 | B630019A10Rik | AC110166.1 | AC126455.1 | AC154568.1 |

| | | | | |
|---|---|---|---|---|
| Csnka2ip | 2810407A14Rik | Gm10785 | Gm2885 | AC151299.2 |
| AC118240.2 | AC093479.1 | Gm15976 | AC168090.1 | BC002059 |
| 1700010K23Rik | AC110241.2 | Atp5o.1 | 1700122H20Rik | Zfp960 |
| 4933411O13Rik | AC125199.2 | D430001F17Rik | AC122413.1 | Zfp97 |
| AC154428.1 | 2310079G19Rik | AC144408.3 | AC122413.2 | AC129328.1 |
| 4930428D20Rik | 2310061N02Rik | AC144408.2 | Gm1604a | Gm6712 |
| Speer2 | 2310034C09Rik | AC144408.1 | AC117241.1 | AC154200.1 |
| AC154719.1 | 2310057N15Rik | 1700048M11Rik | Gm9992 | Gm26873 |
| D16Ertd519e | Gm5965 | 4930563D23Rik | AC119998.3 | Vmn2r90 |
| AC154345.1 | AC125199.4 | AC162305.1 | AC119998.5 | Fpr-rs4 |
| 4931420L22Rik | AC125199.3 | AC174448.1 | E430024P14Rik | Vmn2r124 |
| AC154242.1 | Krtap14 | AC117775.1 | AC164314.2 | Vmn2r91 |
| AC154782.1 | Krtap15 | Gm26626 | Fgfr1op | Vmn2r92 |
| AC129186.1 | Krtap19-9b | CT025774.1 | T2 | Vmn2r94 |
| AC166995.1 | Krtap22-2 | AC140346.1 | Gm16702 | Vmn2r93 |
| Gm15555 | Gm10229 | 1700029J03Rik | Gm17087 | Vmn2r95 |
| AC121777.1 | Gm10228 | AC135673.1 | 6530411M01Rik | Vmn2r96 |
| AC145744.1 | 1110025L11Rik | AC160993.1 | Gm17728 | Vmn2r-ps117 |
| Gm45030 | Gm6358 | Gm5678 | Qk | Vmn2r97 |
| AC122509.1 | Gm10061 | Dopey2 | 1700110C19Rik | Vmn2r98 |
| Gm45029 | Gm9789 | 2310043M15Rik | A230009B12Rik | CT573043.1 |
| Gm30790 | Gm7735 | AC168220.3 | Gm16168 | Vmn2r99 |
| 1700041M19Rik | 1110057P08Rik | AC165961.3 | Park2 | CT030713.2 |
| Gm11146 | Krtap21-1 | Gm48984 | Gm28505 | Vmn2r100 |
| Gm17333 | Krtap6-2 | CT030190.1 | 4732491K20Rik | Vmn2r101 |
| AC098883.2 | AC122381.2 | Dscr3 | Mrgprh | Vmn2r102 |
| AC122375.2 | AC131339.1 | AC165271.1 | Smok2a | Vmn2r103 |
| 1700066C05Rik | AC131339.4 | Gm7976 | Smok2b | Vmn2r104 |
| AC122392.2 | AC131339.3 | AC154546.2 | Tcte2 | Fpr-rs7 |
| AC122817.1 | AC133505.1 | AC154691.1 | Gm16050 | Fpr-rs6 |
| Gm21833 | 1110008E08Rik | 2810404F17Rik | Gm16052 | Vmn2r105 |
| AC108826.1 | AC133505.2 | Gm15340 | Gm16049 | Vmn2r106 |
| AC173486.1 | AC160759.1 | Wrb | Gm7168 | Vmn2r107 |
| A730009L09Rik | AC160759.2 | C030010L15Rik | Gm7356 | Vmn2r108 |
| CT027693.2 | Gm17518 | B230307C23Rik | CT010437.2 | Vmn2r109 |
| Atp5j | AC134560.1 | A630089N07Rik | Gm3417 | Gm5145 |
| Gm10791 | 1110004E09Rik | AC165953.2 | 9030025P20Rik | Vmn2r110 |
| Gm49227 | Gm15965 | AC165953.3 | Gm3448 | Fpr-rs3 |
| Gm49226 | 4931406G06Rik | 1700102H20Rik | Gm3435 | AC109204.1 |
| AC126936.1 | 4932438H23Rik | AC097366.1 | Tcte3 | Gm5092 |
| AC163352.1 | AC034116.3 | AC173485.1 | AC154507.2 | CT010433.1 |
| AC164425.1 | Gm15966 | AC140300.3 | Gm5091 | Zfp54 |
| AC135964.2 | Gm21970 | AC140300.1 | AC154507.3 | Zfp51 |
| AC135964.1 | AC150035.3 | Gm29050 | A930024N18Rik | Gm26753 |
| AC129178.1 | Gm15964 | 3300005D01Rik | AC154378.1 | 9330136K24Rik |
| AC165344.1 | Atp5o | Gm26848 | CT033750.2 | Zfp52 |

| | | | | |
|---|---|---|---|---|
| Zfp983 | Prss34 | 4833413E03Rik | Gm20522 | Olfr97 |
| Gm10509 | Prss28 | Gm17276 | Gm17705 | Olfr98 |
| Gm10226 | Narfl | Gm26858 | Gm16181 | Olfr99 |
| Zfp760 | Fam173a | Cyp4f17 | Gm11131 | Olfr101 |
| Zfp820 | Gm26694 | Cyp4f37 | Gm19553 | Olfr102 |
| Zfp995 | Gm20683 | Zfp871 | Gm9573 | Olfr103 |
| Zfp942 | A930017K11Rik | Gm17115 | Dpcr1 | Olfr104-ps |
| Zfp943 | D630044L22Rik | Gm26693 | Gm20483 | Olfr105-ps |
| Gm9772 | 1700022N22Rik | Zfp870 | Gm4577 | Olfr106-ps |
| Zfp947 | Gm8186 | Zfp472 | Gm20443 | Olfr107 |
| Zfp994 | Tmem8 | Zfp952 | Gm20442 | Olfr108 |
| Zfp944 | Gm8225 | Zfp763 | 4833427F10Rik | Olfr109 |
| Zfp758 | 1700049J03Rik | 4921501E09Rik | Ppp1r18os | Olfr110 |
| Zfp946 | Gm17218 | Zfp563 | Gm16279 | Olfr111 |
| Vmn2r111 | Gm17382 | Olfr55 | Gm20508 | Olfr112 |
| Vmn2r112 | Gm20468 | Olfr239 | A930015D03Rik | Olfr113 |
| Gm9805 | Itpr3os | Olfr1564 | H2-T24 | Olfr114 |
| Gm5493 | Gm26724 | Zfp955a | H2-T22 | Olfr115 |
| Gm16386 | AC125141.2 | Olfr63 | Gm6034 | Olfr116 |
| Zfp945 | AC125141.1 | Zfp955b | Gm11127 | Olfr117 |
| Vmn2r113 | AC125141.3 | Zfp81 | Gm7030 | Olfr118 |
| Vmn2r-ps130 | AC125141.4 | Zfp101 | BC023719 | Olfr119 |
| Zfp40 | AC127341.3 | 2-Mar | Gm19684 | Olfr120 |
| Vmn2r114 | AI413582 | 4931413I07Rik | 2410017I17Rik | Olfr121 |
| Vmn2r115 | AC127341.5 | CT030732.1 | Gm8909 | Olfr122 |
| Vmn2r116 | Gm15420 | Platr17 | Gm20478 | Olfr123 |
| Vmn2r117 | Gm15458 | BC051226 | H2-T3 | Olfr124 |
| Gm26695 | D17Wsu92e | Gm19412 | Gm20546 | Olfr125 |
| Gm49092 | Uhrf1bp1 | Gm26940 | H2-M10.3 | Olfr126 |
| 1520401A03Rik | Gm15597 | H2-Ke6 | H2-M11 | Olfr127 |
| 9530082P21Rik | Gm15598 | BC051537 | H2-M9 | Olfr128 |
| Gm49163 | Gm20109 | Gm20496 | H2-M1 | Olfr761 |
| Gm16275 | E230001N04Rik | Gm15821 | H2-M10.5 | Olfr129 |
| D930048G16Rik | Pnpla1os | Gm20506 | H2-M10.6 | Olfr130 |
| Prss32 | 4930539E08Rik | Gm20513 | 1700031A10Rik | Olfr131 |
| Gm15947 | 1700030A11Rik | Btnl1 | Znrd1 | Olfr132 |
| Dcpp1 | Gm16191 | BC051142 | Znrd1as | Olfr133 |
| Dcpp2 | Gm16196 | Btnl4 | 2410137M14Rik | Olfr134 |
| Dcpp3 | Gm16195 | Btnl6 | H2-M5 | Olfr135 |
| Gm5225 | Gm16194 | Gm20463 | Olfr90 | Olfr138 |
| Gm43796 | Gm26885 | Gm20461 | Olfr91 | Olfr137 |
| Rab26os | Gm17657 | Skiv2l | Olfr92 | Olfr136 |
| Slc9a3r2 | Tbc1d22bos | Gm20547 | Olfr93 | Esp36 |
| Cramp1l | Gm28043 | Gm20481 | Olfr94 | Esp34 |
| Gm38655 | 1700097N02Rik | Gm10501 | Olfr95 | Esp31 |
| BC003965 | Gm15318 | Vars | Olfr96 | Esp24 |

| | | | | |
|---|---|---|---|---|
| Gm21903 | 1700008K24Rik | Gm26749 | Gm28357 | Gm21721 |
| Esp23 | Gm16555 | Gm10093 | Gm29351 | Gm29329 |
| Esp18 | Gm16554 | Gm26637 | Gm29349 | Gm21812 |
| Esp16 | AY702103 | Gm17315 | Gm20918 | Gm21874 |
| Esp15 | Gm7334 | Gm10190 | Gm29353 | Gm20821 |
| Gm26917 | Gm19585 | C230072F16Rik | Gm21820 | Gm21310 |
| Gm42418 | Gm27217 | Gm11096 | Gm29158 | Gm28509 |
| AY036118 | 4932415M13Rik | Gm6594 | Gm21854 | Gm20834 |
| Esp38 | 1700025K24Rik | Gm29052 | Gm20914 | Gm20737 |
| Crisp3 | Gm26547 | C430042M11Rik | Gm28171 | Gm21775 |
| Esp8 | Vmn2r118 | 1110020A21Rik | Gm28173 | Gm21900 |
| Esp6 | Zfp119a | Gm28528 | Gm29194 | Gm29527 |
| Gm44501 | Zfp959 | Gm29418 | Gm21778 | Gm29522 |
| Esp4 | Zfp119b | 2010106C02Rik | Gm29198 | Gm20777 |
| Esp3 | Gm16712 | Gm10309 | Gm29193 | Gm28955 |
| Esp1 | Ccdc94 | Gm29168 | Gm28430 | Gm28954 |
| Mut | Gm20219 | 4833418N02Rik | Gm21746 | Gm20828 |
| 1700071M16Rik | 2410015M20Rik | Gm15978 | Gm28593 | Gm28656 |
| Runx2os3 | 1700061G19Rik | 0610012D04Rik | Gm20873 | Gm20812 |
| Runx2os2 | Gm17949 | Gm28676 | Gm21719 | Gm37739 |
| Runx2os1 | Gm17168 | Gm26612 | Gm28444 | Gm20807 |
| B230354K17Rik | Gm11110 | Gm10184 | Gm20830 | Gm38084 |
| Gm17080 | Vmn2r120 | Gm10308 | Gm28442 | Gm37231 |
| Gm16172 | Gm21834 | Gm6741 | Gm28445 | Gm21891 |
| 1600014C23Rik | Rpl7a-ps5 | Gm15404 | Gm28999 | Gm21425 |
| F630040K05Rik | Nudt12os | Gm1976 | Gm21244 | Gm37252 |
| Gm26785 | Gm29051 | Gm26734 | Gm28398 | Gm21828 |
| Gm5093 | A930002H24Rik | Gm20939 | Gm28395 | Gm21440 |
| Gm26904 | AU016765 | Gm29277 | Gm21788 | Gm21454 |
| Gm47119 | Gm17133 | Gm29089 | Gm28394 | Gm37147 |
| 2310039H08Rik | 4930583I09Rik | Zfy1 | Gm28393 | Gm38159 |
| Gm16494 | 1700016K05Rik | Uba1y | Gm20826 | Gm21725 |
| Gm5814 | Gm47471 | Gm28588 | Gm28575 | Gm37870 |
| 1700001C19Rik | Gm9984 | Gm28587 | Gm28571 | Gm37378 |
| Gm20517 | 2410021H03Rik | Gm29650 | Gm28570 | Gm21866 |
| Gm21981 | A930029G22Rik | Zfy2 | Gm29049 | Gm20773 |
| Tomm6os | C030034I22Rik | Gm4064 | Gm20825 | Gm37561 |
| Frs3os | 5031415H12Rik | Gm10256 | Gm29046 | Gm37263 |
| Gm15556 | Gm20703 | Gm10352 | Gm29043 | Gm21767 |
| 1700122O11Rik | Gm28727 | Gm29289 | Gm29044 | Gm37130 |
| 1700067P10Rik | Gm26510 | Gm21677 | Gm28147 | Gm37467 |
| 9830107B12Rik | Gm26561 | Gm21693 | Gm28331 | Gm38363 |
| A530064D06Rik | Gm16519 | Gm21704 | Gm20815 | Gm30174 |
| Trem3 | Gm4707 | Gm21708 | Gm29364 | Gm30353 |
| B430306N03Rik | Gm15641 | Gm3376 | Gm29363 | Gm30686 |
| Gm45330 | BC027072 | Gm28242 | Gm21292 | Gm37654 |

| | | | | |
|---|---|---|---|---|
| Gm37572 | Gm36950 | Gm28246 | Gm21562 | Gm28264 |
| Gm37344 | Gm20909 | Gm28249 | Gm28298 | Gm28463 |
| Gm37059 | Gm20865 | Gm28250 | Gm21572 | Gm28585 |
| Gm31942 | Gm37462 | Gm28244 | Gm28297 | Gm20896 |
| Gm37440 | Gm38370 | Gm29021 | Gm21723 | Gm28470 |
| Gm37577 | Gm38054 | Gm28880 | Gm28296 | Gm28469 |
| Gm20822 | Gm37690 | Gm28454 | Gm21821 | Gm28332 |
| Gm21809 | Gm32114 | Gm28878 | Gm28295 | Gm28333 |
| Gm20877 | Gm37075 | Gm29399 | Gm21842 | Gm28336 |
| Gm38003 | Gm38371 | Gm28799 | Gm28170 | Gm28338 |
| Rbm31y | Gm37657 | Gm28798 | Gm28764 | Gm28811 |
| Gm37865 | Gm37721 | Gm20894 | Gm28762 | Gm28810 |
| Gm36941 | Gm21366 | Gm28938 | Gm28763 | Gm20835 |
| Gm21904 | Gm33954 | Gm29122 | Gm21588 | Gm29027 |
| Gm37937 | Gm37538 | Gm28985 | Gm28761 | Gm29028 |
| Gm20772 | Gm34217 | Gm28725 | Gm21599 | Gm29023 |
| Gm20831 | Gm37952 | Gm28984 | Gm28765 | Gm29025 |
| Gm37286 | Gm37454 | Gm28987 | Gm21916 | Gm28212 |
| Srsy | Gm37840 | Gm28986 | Gm29191 | Gm20905 |
| Gm38028 | Gm37574 | Gm28129 | Gm28210 | Gm28820 |
| Gm21764 | Gm35670 | Gm28127 | Gm29074 | Gm29671 |
| Ssty1 | Gm37687 | Gm28128 | Gm28518 | Gm20738 |
| Gm38072 | Gm38296 | Gm28132 | Gm28519 | Gm28545 |
| Gm37473 | Gm36345 | Gm28130 | Gm28520 | Gm28547 |
| Gm37656 | Gm37157 | Gm28131 | Gm21626 | Gm28771 |
| Gm38127 | Gm20809 | Gm21679 | Gm28521 | Gm28772 |
| Gm38361 | Gm37434 | Gm29316 | Gm21633 | Gm20897 |
| Gm38209 | Gm30045 | Gm29315 | Gm28522 | Gm28774 |
| Gm21822 | Gm21921 | Gm28216 | Gm28517 | Gm21732 |
| Gm37734 | Gm37112 | Gm28217 | Gm29532 | Gm28775 |
| Gm38013 | Gm37071 | Gm28092 | Gm29531 | Gm28176 |
| Gm37948 | Gm37547 | Gm28091 | Gm28464 | Gm28786 |
| Gm34550 | Gm30705 | Gm29580 | Gm29656 | Gm29360 |
| Gm37544 | Gm21773 | Gm28089 | Gm29655 | Gm29497 |
| Gm34716 | Gm37451 | Gm28088 | Gm29653 | Gm29498 |
| Gm37635 | Gm37875 | Gm29579 | Gm29654 | Gm36782 |
| Gm35070 | Gm31422 | Gm21529 | Gm29080 | Gm29839 |
| Gm37998 | Gm37798 | Gm29581 | Gm29081 | Gm28595 |
| Gm38168 | Gm37346 | Gm21539 | Gm29077 | Gm29433 |
| Gm37740 | Gm38136 | Gm29578 | Gm29078 | Gm29215 |
| Gm36261 | Gm37808 | Gm21780 | Gm29225 | Gm21882 |
| Gm37898 | Gm32181 | Gm28486 | Gm29226 | Gm21801 |
| Gm36467 | Gm28853 | Gm28487 | Gm20855 | Gm28850 |
| Gm37927 | Gm28852 | Gm28488 | Gm29386 | Gm21914 |
| Gm36929 | Gm28858 | Gm28491 | Gm29384 | Gm28617 |
| Gm38174 | Gm28245 | Gm28490 | Gm28260 | Gm28619 |

| | | | | |
|---|---|---|---|---|
| Gm21852 | Gm28554 | Gm20929 | Gm29130 | Gm29380 |
| Gm28993 | Gm29056 | Gm28276 | Gm20978 | Gm29381 |
| Gm28994 | Gm29098 | Gm28274 | Gm29132 | Gm20852 |
| Gm21797 | Gm21170 | Gm28291 | Gm21450 | Gm29003 |
| Gm20795 | Gm29373 | Gm28293 | Gm28889 | Ssty2 |
| Gm28427 | Gm29370 | Gm29305 | Gm28890 | Gm28612 |
| Gm21800 | Gm29166 | Gm29303 | Gm28691 | Gm28613 |
| Gm28426 | Gm28194 | Gm29557 | Gm28692 | Gm29219 |
| Gm28425 | Gm28754 | Gm29301 | Gm28690 | Gm29632 |
| Gm28834 | Gm28753 | Gm20931 | Gm28689 | Gm29082 |
| Gm29537 | Gm29070 | Gm29555 | Gm28687 | Gm29217 |
| Gm29274 | Gm28726 | Gm28355 | Gm20987 | Gm29636 |
| Gm29275 | Gm29270 | Gm28352 | 1700040F15Rik | Gm29662 |
| Gm29645 | Gm29271 | Gm28201 | Gm29207 | Gm29660 |
| Gm28312 | Gm28681 | Gm28202 | Gm29204 | Gm29450 |
| Gm29042 | Gm28684 | Gm28206 | Gm29206 | Gm29449 |
| Gm28311 | Gm28679 | Gm28207 | Gm29203 | Gm20816 |
| Gm28310 | Gm29511 | Gm28208 | Gm29612 | Gm29446 |
| Gm28568 | Gm29117 | Gm29222 | Gm29343 | Gm28633 |
| Gm29286 | Gm20883 | Gm29221 | Gm29342 | Gm28462 |
| Gm29285 | Gm29116 | Gm29220 | Gm28600 | Gm29409 |
| Gm28280 | Gm29321 | Gm29405 | Gm20823 | Gm28632 |
| Gm28278 | Gm28317 | Gm29406 | Gm29547 | Gm33815 |
| Gm28279 | Gm28318 | Gm29404 | Gm29209 | Gm28824 |
| Gm28886 | Gm28313 | Gm28964 | Gm29549 | Gm28081 |
| Gm29426 | Gm28315 | Gm28709 | Gm29421 | Gm28079 |
| Gm28887 | Gm28316 | Gm28965 | Gm21627 | Gm28082 |
| Gm29444 | Gm28944 | Gm21943 | Gm29060 | Gm29644 |
| Gm21783 | Gm28945 | Gm28532 | Gm29061 | Gm21118 |
| Gm29445 | Gm28947 | Gm28966 | Gm29625 | Gm29423 |
| Gm28457 | Gm20920 | Gm28962 | Gm29628 | Gm29425 |
| Gm28458 | Gm28948 | Gm28963 | Gm29622 | Gm28238 |
| Gm28461 | Gm28950 | Gm29466 | Gm29265 | Gm28431 |
| Gm28460 | Gm29063 | Gm29467 | Gm20908 | Gm28432 |
| Gm28851 | Gm28197 | Gm20963 | Gm28252 | Gm28842 |
| Gm28456 | Gm28735 | Gm28326 | Gm28254 | Gm28839 |
| Gm29648 | Gm28732 | Gm28325 | Gm20924 | Gm28668 |
| Gm29646 | Gm28733 | Gm28284 | Gm29457 | Gm20869 |
| Gm28549 | Gm28233 | Gm29368 | Gm28561 | Gm28482 |
| Gm29210 | Gm28235 | Gm21413 | Gm29584 | Gm28485 |
| Gm29213 | Gm28236 | Gm28073 | Gm28134 | Gm20870 |
| Gm28550 | Gm20747 | Gm21419 | Gm28135 | Gm29182 |
| Gm28604 | Gm29250 | Gm28074 | Gm28133 | Gm29255 |
| Gm28565 | Gm29252 | Gm21428 | Gm28137 | Gm20903 |
| Gm28566 | Gm29248 | Gm29672 | Gm28138 | Gm28259 |
| Gm28697 | Gm28908 | Gm29131 | Gm29379 | Gm29616 |

| | | | | |
|---|---|---|---|---|
| Gm29108 | Gm21317 | Gm29393 | 0710001A04Rik | Gm15337 |
| Gm21913 | Gm29472 | Gm28672 | Gm26658 | Pabpc2 |
| Gm28174 | Gm29473 | Gm28670 | B930094E09Rik | 2900055J20Rik |
| Gm29416 | Gm28226 | Gm28673 | Gm26533 | Gm5689 |
| Gm28789 | Gm28225 | Gm28674 | Gm6665 | Lars |
| Gm20814 | Gm28832 | Gm28930 | Gm26823 | 4933407I08Rik |
| Gm29278 | Gm28472 | Gm29504 | Gm16344 | Gm37797 |
| Gm28606 | Gm28475 | Gm20837 | Gm26717 | Gm10267 |
| Gm20850 | Gm21394 | Gm28300 | A830052D11Rik | Spinkl |
| Gm21776 | Gm29338 | Gm28301 | Gm35060 | Spink11 |
| Gm29313 | Gm21409 | Gm21860 | 4930455D15Rik | Gm10542 |
| Gm29311 | Gm28758 | Gm47283 | 2310026I22Rik | A930012L18Rik |
| Gm29309 | Gm29071 | Gm21748 | Gm10549 | Gm43425 |
| Gm28702 | Gm28348 | Vmn1r238 | Epb41l4aos | A330093E20Rik |
| Gm28701 | Gm28972 | Gm6225 | 4933408B17Rik | 1700044K03Rik |
| Gm28704 | Gm28970 | G430049J08Rik | Gm3550 | 1700065O20Rik |
| Gm28540 | Gm28971 | Gm26865 | 2010110K18Rik | Gm4950 |
| Gm21650 | Gm20854 | Gm10556 | Gm26538 | Gykl1 |
| Gm28541 | Gm28346 | Gm26682 | Gm5239 | Redrum |
| Gm28538 | Gm28345 | Gm17036 | Gm28285 | Gm26742 |
| Gm29569 | Gm29090 | Rpl27-ps3 | 1700066B19Rik | Gm4221 |
| Gm20867 | Gm28152 | Gm28529 | Tmem173 | Gramd3 |
| Gm29568 | Gm28823 | 4921524L21Rik | Gm29417 | Tex43 |
| Gm29566 | Gm28220 | Armc4 | E230025N22Rik | 3-Mar |
| Gm29565 | Gm28218 | Gm5819 | Hars | Gm15345 |
| Gm29146 | Gm29302 | Gm17430 | Gm37751 | 1700011I03Rik |
| Gm28664 | Gm28440 | 4930563E18Rik | Gm38225 | Gm26507 |
| Gm28663 | Gm29091 | Mir133a-1hg | Gm42416 | A730017C20Rik |
| Gm28817 | Gm29297 | 1010001N08Rik | Gm37753 | Gm4951 |
| Gm28816 | Gm28365 | Gm6277 | Gm37013 | Gm4841 |
| Gm20806 | Gm21477 | 3110002H16Rik | Gm37388 | F830016B08Rik |
| Gm28813 | Gm28367 | Gm15956 | Gm10545 | Iigp1 |
| Gm20917 | Gm20906 | Gm29200 | Pcdha11.1 | Bvht |
| Gm28421 | Gm28109 | Gm5160 | Gm38097 | Gm38165 |
| Gm28422 | Gm28108 | Gm10036 | Gm37446 | 1500015A07Rik |
| Gm20916 | Gm28104 | Gm15485 | Pcdhb14 | Gm9949 |
| Gm29606 | Gm28103 | Gm15328 | 4930517L18Rik | Spink10 |
| Gm29190 | Gm28718 | 1700001G01Rik | 3222401L13Rik | Gm36368 |
| Gm28157 | Gm29162 | Gm16090 | Gm37118 | Gm17732 |
| Gm28696 | Gm28866 | Gm10269 | Gm37165 | 2700046A07Rik |
| Gm28898 | Gm28405 | 4930426D05Rik | BC037039 | Gm26972 |
| Gm29324 | Gm28406 | Gm47898 | Gm29994 | St8sia3os |
| Gm28741 | Gm29436 | Gm15972 | 1700086O06Rik | Nars |
| Gm28743 | Gm28977 | Zfp35 | 0610009O20Rik | A330084C13Rik |
| Gm28124 | Gm28407 | Gm9955 | Gm15334 | Raxos1 |
| Gm28126 | Gm29392 | Gm3227 | Gm15336 | Gm17669 |

| | | | | |
|---|---|---|---|---|
| Gm26910 | Gm14966 | Olfr262 | Olfr1494 | A830019P07Rik |
| 4930546C10Rik | Gm14964 | Olfr235 | Olfr1495 | Gm47735 |
| Gm31294 | Gm14967 | Olfr1434 | Olfr1496 | F530104D19Rik |
| 1700061H18Rik | Gm14968 | Olfr1436 | Olfr1497 | A330032B11Rik |
| 4930448D08Rik | Gm17227 | Olfr1437 | Olfr1499 | 5-Mar |
| D730045A05Rik | 1700105P06Rik | Olfr1438-ps1 | Olfr1500 | I830134H01Rik |
| Gm9925 | 2700081O15Rik | Olfr1440 | Olfr1501 | Gm42723 |
| 1700120E14Rik | Pla2g16 | Olfr1441 | Olfr1502 | Gm28991 |
| Gm10532 | Hrasls5 | Gm47243 | Olfr1504 | Cyp2c70 |
| 1700034B16Rik | Gm45736 | Gm47242 | Olfr1505 | A930028N01Rik |
| Gm7276 | 1700092M07Rik | Olfr1442 | C130060C02Rik | Gm27042 |
| Rnf165 | Lbhd1.1 | Olfr1443 | 1500015L24Rik | E030044B06Rik |
| 8030462N17Rik | Ints5.1 | Keg1 | C730002L08Rik | Gm340 |
| A330094K24Rik | Gm10353 | Gm15962 | Tmem2 | AI606181 |
| Atp5a1 | Pcna-ps2 | Olfr1444 | Gm27151 | Gm16541 |
| Gm21886 | Gm26859 | Olfr1445 | C330002G04Rik | Gm47936 |
| Gm16286 | Ppp1r32 | Olfr1446 | 2410080I02Rik | Gm47938 |
| Pqlc1 | 2210404E10Rik | Olfr1447 | 1700028P14Rik | Gm20467 |
| Gm26676 | Gm28347 | Olfr1448 | Gm9493 | Gm20538 |
| Gm47272 | AW112010 | Olfr1449 | Gm6563 | Gm26644 |
| Gm27239 | 1700017D01Rik | Olfr1450 | Gm9938 | Gm20395 |
| 4930594M17Rik | 1700025F22Rik | Olfr1451 | Fam189a2 | 1700039E22Rik |
| 2210420H20Rik | 4930526L06Rik | Olfr1453 | 1700021P04Rik | Tlx1os |
| 4930445N18Rik | Gm28935 | Olfr1454 | Pip5k1bos | Gm29595 |
| 1700095A13Rik | Ms4a4c | Olfr1457 | Fam122a | Gm29593 |
| Zadh2 | Ms4a4b | Olfr1458 | Gm10053 | Gm29594 |
| Fam69c | Gm37387 | Olfr1459 | Cbwd1 | Gm28578 |
| Gm16146 | Gm8369 | Olfr1461 | 2610016A17Rik | 1700016H03Rik |
| Gm5096 | Gm19261 | Olfr1462 | Gm48775 | Gm17018 |
| Gm45871 | Ms4a4d | Olfr1463 | Gm815 | Gm15491 |
| 1700030N03Rik | Oosp3 | Olfr1465 | Gm28228 | Mgea5 |
| Gm48683 | Gif | Olfr1466 | D930032P07Rik | 9130011E15Rik |
| Nudt8.1 | Olfr1417 | Olfr1467 | 4430402I18Rik | 4930505N22Rik |
| Gm16312 | Olfr1418 | Olfr1469 | 1700018L02Rik | Gm26792 |
| Gm17552 | Olfr1419 | Olfr1471 | A930007I19Rik | 2310034G01Rik |
| Gm960 | Olfr1420 | Olfr1472 | 9930021J03Rik | Usmg5 |
| Gm21992 | Olfr1423 | Olfr1474 | 2700046G09Rik | Gm6970 |
| Gm21844 | Olfr1424 | Olfr1475 | Gm7237 | Gm19557 |
| 4930481A15Rik | Olfr1425 | Olfr1477 | Lipo5 | Gm16068 |
| 1700020D05Rik | Olfr1426 | Olfr1480 | Lipo4 | Rpl13a-ps1 |
| Gm16538 | Olfr1427 | Olfr1484 | Lipo3 | Gm45352 |
| Sssca1 | Olfr1428 | Olfr1487 | Lipo2 | Gm26629 |
| Frmd8os | Olfr76 | Olfr1489 | Lipo1 | 1700054A03Rik |
| Gm42067 | Olfr1555-ps1 | Olfr1490 | Gm5519 | Gm10197 |
| Gm10814 | Olfr1431 | Olfr1491 | Gm26902 | Mirt1 |
| Gm28374 | Olfr1432 | Olfr1493-ps1 | Gm5248 | 4833407H14Rik |

| |
|---|
| Nutf2-ps1 |
| Gm16299 |
| 4930552P12Rik |
| Gm17197 |
| B230217O12Rik |
| Fam160b1 |
| Gm26874 |
| Gm16277 |
| 1810007D17Rik |
| 1700019N19Rik |
| Gm29261 |
| Rps12-ps3 |
| Gm33756 |
| 2700089I24Rik |
| Gm17203 |
| E330013P04Rik |
| Gm28351 |
| Fam45a |
| Gm7102 |
| Gm6020 |
| Gm21060 |
| AC123873.3 |
| AC123873.1 |
| AC123873.2 |
| AC126035.1 |
| Gm16367 |
| AC163611.1 |
| AC163611.2 |
| AC140365.1 |
| AC124606.2 |
| AC124606.1 |
| AC133095.2 |
| AC133095.1 |
| CAAA01165726.1 |
| AC133103.4 |
| AC133103.6 |
| AC133103.7 |
| AC133103.5 |
| AC133103.1 |
| AC133103.3 |
| Ccl21b.1 |
| AC087559.3 |
| Gm13298 |
| AC087559.2 |
| Ccl21c |
| Il11ra2.1 |
| Ccl27 |
| CR974586.3 |
| Ccl21c.1 |
| CR974586.4 |
| CR974586.6 |
| CR974586.5 |
| CR974586.2 |
| CR974586.1 |
| AC132444.1 |
| AC132444.3 |
| AC132444.5 |
| AC132444.4 |
| AC132444.2 |
| AC132444.6 |
| AC165294.1 |
| AC165294.2 |
| AC165294.3 |
| AC164084.2 |
| AC164084.3 |
| AC164084.1 |
| AC140325.2 |
| AC140325.1 |
| Gm3286.1 |
| AC140325.3 |
| AC140325.4 |
| Ccl27.1 |
| Il11ra2.2 |
| Ccl19.1 |
| Ccl21a.1 |
| Gm10931 |
| CT868723.1 |
| AC125178.1 |
| AC125178.3 |
| AC125178.2 |
| Vmn1r186 |
| AC102264.1 |
| AC125149.3 |
| AC125149.5 |
| AC125149.1 |
| AC125149.2 |
| AC125149.4 |
| AC234645.1 |
| AC168977.2 |
| AC168977.1 |
| AC149090.1 |
| CAAA01118383.1 |
| Vmn2r122 |
| CAAA01147332.1 |
