## Supplemental Table 2 for "Human-mouse cross-species comparison identifies common and unique aspects of intestinal mesenchyme development"

| Species | Stage and Region | Gene Filter | UMI Count Filter |
|---|---|---|---|
| Mouse | E13.5 | <1000 and >8000 | <1000 and >50000 |
| Mouse | E14.5 | <1000 and >7000 | <1000 and >40000 |
| Mouse | E15.5 | <1000 and >7500 | <1000 and >40000 |
| Mouse | E16 | <1000 and >10000 | <1000 and >60000 |
| Mouse | E17.5 | <1000 and >7500 | <1000 and >40000 |
| Human | 70dpc Duodenum | <300 and >4000 | <300 and >30000 |
| Human | 72dpc Duodenum | <1000 and >10000 | <1000 and >100000 |
| Human | 80dpc Duodenum | <1000 and >7500 | <1000 and >50000 |
| Human | 80dpc Jejunum | <800 and >8000 | <800 and >50000 |
| Human | 80dpc ileum | <1000 and >8000 | <800 and >50000 |
| Human | 101dpc Duodenum | <1000 and >6000 | <500 and >40000 |
| Human | 101dpc Ileum | , <1000 and >6000 | <500 and >40000) |
